# Male-male interactions shape mate selection in *Drosophila*

**DOI:** 10.1101/2023.11.03.565582

**Authors:** Tom Hindmarsh Sten, Rufei Li, Florian Hollunder, Shadé Eleazer, Vanessa Ruta

**Author notes:** These authors contributed equally, and are listed in alphabetical order (Tom Hindmarsh Sten, Rufei Li).

## Abstract

Males of many species have evolved behavioral traits to both attract females and repel rivals. Here, we explore mate selection in *Drosophila* from both the male and female perspective to shed light on how these key components of sexual selection — female choice and male-male competition — work in concert to guide reproductive strategies. We find that male flies fend off competing suitors by interleaving their courtship of a female with aggressive wing flicks, which both repel competitors and generate a ‘song’ that obscures the female’s auditory perception of other potential mates. Two higher-order circuit nodes – P1a and pC1x neurons – are coordinately recruited to allow males to flexibly interleave these agonistic actions with courtship displays, assuring they persistently pursue females until their rival falters. Together, our results suggest that female mating decisions are shaped by male-male interactions, underscoring how a male’s ability to subvert his rivals is central to his reproductive success.

---

In many species, lengthy and elaborate courtship rituals serve as a prelude to copulation^1^. Females are generally believed to be the choosier sex due to the metabolic costs of maternal investment^2–4^. Male courtship displays are therefore thought to have evolved under sexual selection as a form of behavioral ornamentation that advertises a male’s quality as a suitable mate^1–5^. Yet how females assess the behavioral performance of different males to guide their mate choice has often remained elusive^6–8^.

In *Drosophila*, males court females with a complex behavioral display during which they chase, tap, and lick the female while extending and vibrating a single wing to produce a species-specific courtship song^9–11^. Female mating decisions have primarily been studied in the context of single courting pairs and point to the importance of a male’s courtship song as a potent aphrodisiac to ‘charm’^3^ the female and ultimately trigger her sexual acceptance^9,12–17^. In the wild, however, *Drosophila* meet and mate on fermenting fruits where a diversity of species and sexes congregate^18–20^. Mating decisions therefore naturally unfold in a complex social environment where a female is often pursued by an array of prospective sexual partners, creating a fertile ground for competition. Female flies are highly adept at species discrimination and will rarely accept the promiscuous advances of heterospecific males^10,21–25^, yet how a female selects between competing conspecifics remains unclear. Females may choose a mate based on his preferable morphological or behavioral traits, aligned with the proposal that male characteristics and female preferences coevolve in a self-reinforcing fashion through sexual selection^3,26–29^. However, female preferences may also be indirectly shaped by male-male rivalry, in which each male seeks to maximize his own chances of reproductive success while minimizing those of his opponents, reinforcing traits that facilitate successful male competition^3,26,30–32^. At an extreme, competing males may engage in scramble competition, attempting to copulate with a female irrespective of her preferences, eroding her agency to choose^2,33^. Elucidating the logic of female mating strategies thus requires parallel insight into female sensory preferences, how male courtship is altered by the presence of a rival male, and in turn, how male-male interactions shape female sensory perception.

Here we examine the behavioral dynamics of competitive courtship in *D. melanogaster* to shed light on the mechanisms of mate choice. We reveal that while females must hear the species-specific courtship song produced by conspecific males to become sexually receptive, they do not appear to explicitly compare the behavioral performance of competing males to arbitrate their mating decisions. Rather, mate selection is honed by the emergence of antagonistic behaviors^34^ that confer males with a transient competitive advantage, serving to fend off other males as well as generating an agonistic ‘song’^35^ that jams the female’s auditory pathways, hindering her recognition of other potential mates. Reproductive success in the face of competition therefore requires that males rapidly interleave their courtship of a female with aggressive subversions of the courtship displays of their rivals. We find that this flexible alternation between male courtship and agonistic behaviors is mediated by the transient recruitment of a population of higher-order pC1x neurons that operate in parallel to the courtship-promoting P1a neurons^36–42^, enabling a male to adaptively tailor his actions to compete in a complex social landscape. Together, our work suggests how mate choice unfolds through the complex interplay between female preferences and male-male interactions.

## Conspecific mate selection in females

To investigate how females evaluate potential mates based on their courtship performance, we paired a single female with two equivalently-reared isogenic males and performed high-resolution postural tracking^43^ of each member of the triad until copulation (**Figure 1A**; **Video S1**). This simplified assay recapitulates many aspects of the rich social interplay that a female encounters in the wild^1,10,18–20^, where she is simultaneously pursued by multiple males, while the restricted geometry of the chamber assures that the female eventually accepts a sexual partner by precluding her from decamping. To explore which attributes of a male may predict his mating success, we trained behavioral classifiers^44^ to quantitatively characterize male behavioral actions in malemale-female (MMF) triads (**S1A-C**; **Video S1**). However, while we observed substantial variation in the behavioral performance of competing males – including their average speed, the fidelity of their pursuit of the female, and the frequency of their wing extensions or copulation attempts – none of these characteristics were predictive of which male eventually would copulate with the female (**Figure 1B**). Indeed, the ‘winning’ male could only be reliably distinguished in the final seconds prior to copulation, as he assumed the optimal copulatory position immediately behind the female’s genitalia, moved closer to her, and performed a final unilateral wing extension while attempting to mount (**Figures 1C-D**). Thus, although both males vigorously pursued the female for many minutes (average time to copulation = 414.3sec, range: 83-1267 sec), copulations occurred suddenly and could not be readily predicted by the overt actions or attributes of either male.

**Figure 1.**
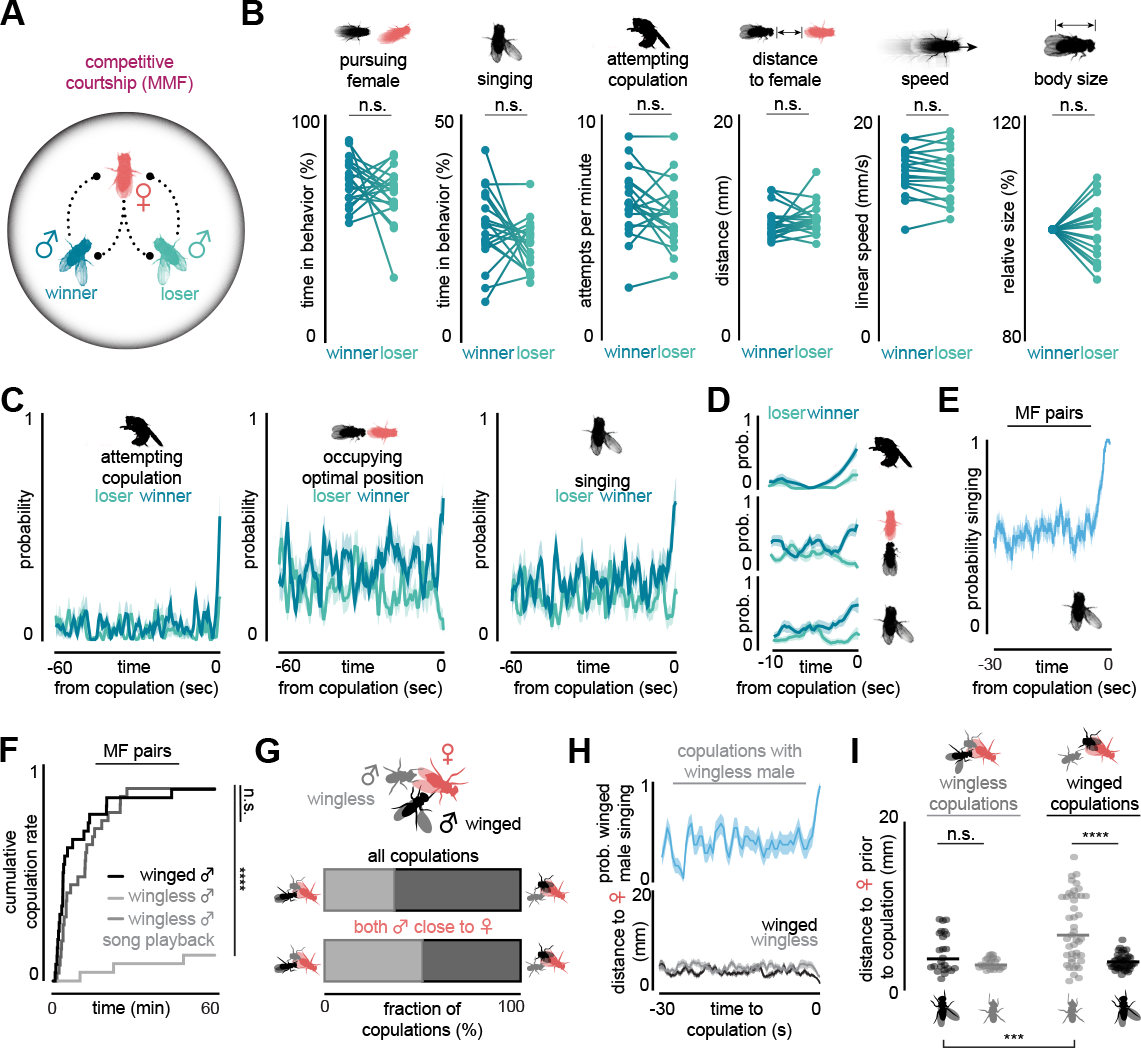
| Courtship cues signal species identity but not individual identity. (**A**)Schematic of competitive courtship triads, in which two isogenic males are paired with and compete for a single virgin female. Comparison of behavioral and morphological metrics for the male who succeeded in copulating with the female (winner) and the other male (loser) in competitive courtship triads. Individual points represent the average across an entire assay. (**C** and **D**) Average behavioral metrics comparing the winner and the loser male in the last 60 seconds before copulation (C) and focusing on the last 10 seconds before copulation for those same behaviors (D). (**E**) Probability of males performing unilateral wing extensions across the last 30 seconds before copulation in male-female (MF) pairs. (**F**) Cumulative fraction of copulations in male-female (MF) pairs during 60-minute assays, in which one female was paired with either a single intact winged male (black), a mute wingless male (light gray), or a mute wingless male while synthetic *D. melanogaster* pulse song was played back to the pair (dark gray). (**G**) Top: Fraction of all competitive courtship assays (MMF, n = 68 triads) in which the female copulated with the wingless male (gray) or the winged male (black). Bottom: Same as top but including only the subset of assays when both males were close (average distance < 5mm) to the female in the 0.5-1 sec before copulation (n = 30 triads). (**H**) Probability of the intact male singing (top) and the distance of both males to the female (bottom) over the last 30 seconds before copulation in trials in which the wingless male copulated with the female. (**I**) Average distance of the intact male (black) and the wingless male (gray) to the female immediately (0.5-1 sec) prior to copulation in assays in which the wingless male (left) or intact male (right) copulated with the female. Shaded lines show mean ± s.e.m.; dots are individual animals. n.s., P >= 0.05; ***P<0.001; ****P < 0.0001. Details of statistical analyses and sample sizes are given in **TableS2**. See also **Figure S1** and **Video S1**.

*Drosophila* use several modes of communication during courtship that cannot be readily captured by video alone. In particular, females have been proposed to arbitrate their mating decisions based on the statistics and quality of a male’s song^17,23,45^, which is patterned into bouts of sinusoidal humming and pulse trains^46,47^. Male courtship songs, in turn, have rapidly diversified across the *Drosophila* lineage, giving rise to species-specific spectral properties that females use to recognize a conspecific mate^47–49^. Consistent with the importance of courtship song in promoting a female’s sexual receptivity^9,12–15,17^, we found that males invariably performed a unilateral wing extension immediately prior to copulation in male-female (MF) pairs^50^ (**Figure 1E**). Moreover, as shown previously^12,17,51^, the sexual receptivity of a female was strongly suppressed when she was paired with a mute male whose wings had been surgically removed, but could be restored by playing synthetic *D. melanogaster* pulse song through a speaker beneath the floor of a perforated assay chamber (**Figure 1F**). Song is therefore critical to promote female receptivity, raising the possibility that females may actively select between competing mates based on their acoustic performance. To assess this, we examined the preferences of females offered a choice between an intact male and a mute wingless male. Unexpectedly, females copulated with the wingless male in a significant fraction of trials (**Figure 1G**, 35.3% copulation rate; 532.3±317.3 sec to copulation, mean±st.d.; n=68 triads) despite rarely mating with mute males when paired with them alone (**Figure 1F**, 13.6% copulation rate; 1733±1167 sec to copulation, mean±st.d.) or in combination with a second wingless male (**Figure S1D**). Copulations with the wingless male predominantly occurred when the winged male was close to the female and singing (**Figures 1H-I**, 87.5% of copulations accompanied by wing extensions), suggesting that the mute male may exploit mating opportunities created by the winged male’s song (**Video S2**). In contrast, winged males could successfully copulate with the female even when their mute rival was several body lengths away (**Figure 1H-I**), indicating that the ability to independently serenade the female confers them with their distinct reproductive advantage. Indeed, in trials where both males were near the female (< 5mm) immediately prior to copulation, the winged male’s odds of being successful were reduced to chance (**Figure 1G**), suggesting that she may not be able to discern which male is singing to her. Females therefore do not appear to directly compare the acoustic performance of competing males but rather rely on courtship song as an instructive cue to signal the presence of an appropriate mate. Consistent with this, in other *Drosophila* species females have been shown to mate with mute heterospecific males if serenaded with their conspecific song^24,25^. Courtship song thus appears to signal species identity but not the individual identity of rival males, suggesting additional criteria are used to shape mate selection.

## Competing males interleave courtship and agonistic displays

In many species, male-male interactions operate in parallel with female preferences to determine mating success^2,3,32^. To assess the role of male competition in shaping mate selection, we leveraged our classifiers to explore the interplay between competing males in MMF triads (**Figures 2A-B, S1A-C, S2A-C**; **Video S1**). In contrast to MF courtship trials, where the sole male can pursue the female at a close distance for many minutes uninterrupted, both males of a triad jockeyed for the position behind the female, closest to her genitalia (**Figures 2A, 2C, S2E**; **Video S3**). Each male rapidly alternated between closely following and singing to the female, jostling the other male, or circling the assay chamber far from the female before quickly rejoining the group (**Figures 2C-D**). In the brief moments when a male disengaged from courtship pursuit, he was typically immediately replaced by his competitor (**Figures 2C-D, S2E**). As a consequence, the female was almost constantly pursued by at least one suitor (**Figure 2C, S2A**), despite the fact that each male spent significantly less time overall chasing and singing to the female than in non-competitive courtship assays (**Figure 2B**). Bouts of song performed synchronously by both males were rare and consistent with chance overlap, suggesting that males did not appear to explicitly avoid singing at the same time (**Figure S2C**). All animals in the triad also exhibited higher linear speeds (**Figures 2B, S2A**) compared to the more slowly unfolding courtship dynamics observed in MF pairs (**Figures 2B, S2A**).

**Figure 2.**
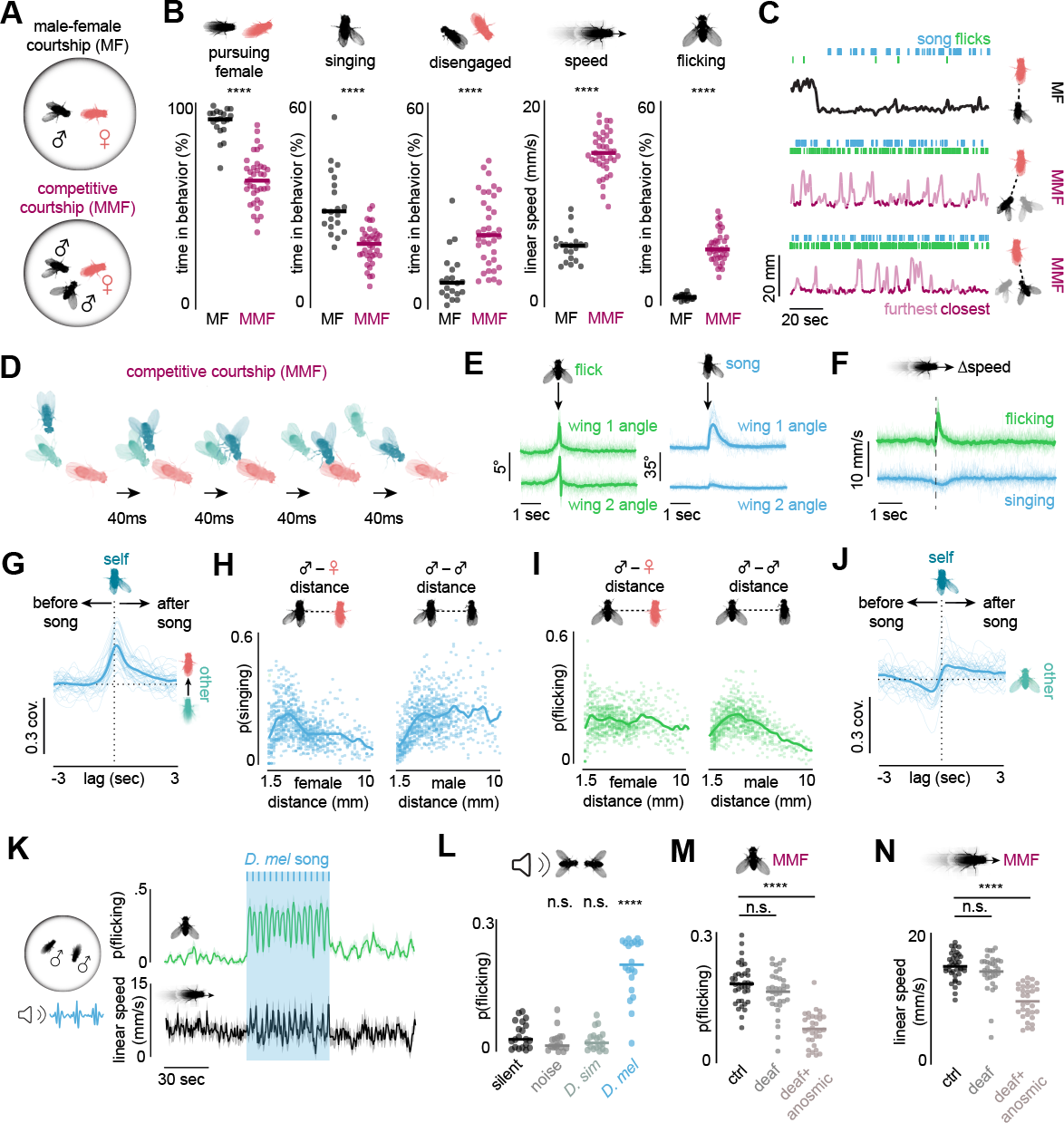
|Male sensory cues evoke agonistic actions during competitive courtship. (**A**) Schematic of male-female (MF) and competitive courtship assays (MMF). (**B**) Comparison of behavioral metrics of males in MF and MMF assays. Dots represent individual males, lines denote average. (**C**) Representative examples showing distinct behavioral dynamics in a MF (top) and a MMF assay (middle, bottom). Green hatch marks denote bilateral wing flicks, and blue hatch marks denote unilateral wing extensions. In MF plot, the black line shows male-female distance. MMF plots show the distance of each of the two males to the female (top vs. bottom), with dark purple regions indicating periods where the male occupied the position closest to the female. (**D**) Snapshot of a typical behavioral sequence in which one male (dark blue) overtakes his rival (cyan) to assume the position closest to the female, while concurrently transitioning from performing a bilateral wing flick to a unilateral wing extension. (**E** and **F**) Average wing angles (E), and linear speed (F) of males, aligned to the onset of wing extensions (blue) and wing flicks (green). Wing 1 denotes the wing whose angle changed most following onset. (**G**) Cross-covariance between one male’s unilateral wing extensions and the other male’s directed approach towards the female. Positive lags indicate that unilateral wing extensions lead in time. (**H** and **I**) Probability of males performing unilateral wing extensions (H) and bilateral wing flicks (I) as a function of their distance to the female (right) and their rival male (left). Dots denote individual animals at each distance, line denotes average. (**J**) Cross-covariance between one male’s unilateral wing extensions and the other male’s bilateral wing flicks. Positive lags indicate that unilateral wing extension lead in time. (**K**) Probability of males exhibiting bilateral wing flicks (top) and male linear speed (bottom) prior to, during, and after 1 minute of synthetic *D. melanogaster* song played to two males in a perforated chamber. Shaded region denotes playback period, darker blue hatches denote individual trains of pulse song. (**L**) Probability of males exhibiting bilateral wing flicks during periods of no sound (black), or epochs of playback of white noise (gray), synthetic *D. simulans* song (light gray), and synthetic *D. melanogaster* song (blue). (**M** and **N**) Probability of males exhibiting bilateral wing flicks (M) and average male linear speed (N) in MMF assays when both males were intact (ctrl, black), had their aristae removed (deaf, gray), or had the third antennal segment removed. Lines with dots denote mean and individual animals; shaded lines show mean ± s.e.m.; thin lines denote individual animals. n.s., P > 0.05; ****P < 0.0001. Details of statistical analyses and sample sizes are given in **TableS2**. See also **Figures S1, S2** and **S3** and **Videos S2, S3, S4, S5**, and **S7**.

Beyond changes to the structure of their courtship pursuit (**Figures 2B-D, S1C, S2A-C**), competing males also performed a distinct wing display that was rarely observed in MF courtship, in which they flicked both their wings to a 30-45° angle over the course of 150-300ms (**Figures 2A-E, S2C**; **Video S4**). During wing flicks, males concurrently increased their linear speed (**Figure 2F**), creating a behavioral pattern reminiscent of the agonistic “charges” or wing threats that underlie the aggressive interactions male flies employ to defend scarce resources from encroaching rivals^35,52–54^. The selective emergence of wing flicks in the context of competitive courtship (**Figures 2B-C**) reinforces that they represent a form of male-male aggression, performed in tandem with courtship displays as males attempt to both attract and defend potential mates.

To gain insight into how competing males alternate between these agonistic and courtship displays, we characterized the spatiotemporal relationships between the behaviors performed by each member of the triad. Computing the cross-covariance between different male component behaviors (**Figure S2G)** revealed a positive correlation between when one male sang to the female and when his rival subsequently oriented towards and approached the courting pair (**Figures 2G, S2H-K, Video S5**), suggesting that males may eavesdrop on the courtship song of another male to recognize the presence of a potential mate. Opponent males performed unilateral wing extensions predominantly when they were within 2-3 mm of the female (**Figures 2H, S2D**), consistent with observations that males preferentially sing only in close proximity to a female^9,10^, but suppressed their singing in the immediate vicinity of a competitor (< 2 mm) (**Figures 2H, S2D**). Conversely, wing flicks were more likely to be elicited when the two males were within a few millimeters of one another, independent of their position relative to the female (**Figures 2I, S2D**), suggesting that these agonistic actions are promoted by sensory cues emanating from other males. Indeed, the onset of one male’s unilateral wing extension was correlated with a subsequent increase in the probability that his rival performed a wing flick (**Figure 2J, S2L-M**). Hearing the song of another male may therefore signify the presence of a competitor as well as a potential mate, provoking males to become aggressive. To test this, we played synthetic *D. melanogaster* pulse song to pairs of males (**Figure S3A**) and found that with each bout of song, males vigorously flicked their wings and charged across the floor of the chamber, even in the absence of a female target to vie for (**Figure 2K**; **Video S4**). These behaviors were not observed in the period prior to song onset or during playback of white-noise or heterospecific courtship song (**Figures 2L, S3B-C**), suggesting that males selectively engage in these competitive interactions upon detecting a conspecific rival courting nearby. These agonistic responses to song (**Figures 2L, S3D**) contrast with prior observations that song playback evokes courtship between wingless males^17,55–59^, indicating that song can also promote male sexual arousal but that the presence of wing flicks may actively suppress intrasexual courtship^1,17^.

Playback of song to a MF pair did not trigger males to abandon their pursuit of the female (**Figure S3E-F**). Rather, males continued to court but began to rapidly interleave their unilateral wing extensions with bilateral wing flicks, and both members of the pair increased their linear speed for the duration of song playback, mimicking the behavioral dynamics of triads (**Figure S3G-H**). Auditory cues emanating from a rival male are thus sufficient to transform the behavioral patterns of courting males, promoting the emergence of agonistic wing flicks without diminishing their courtship. While deafening both males in a triad by surgically removing their aristae had little impact on their rate of flicking, removal of their third antennal segment, which houses both the aristae and most olfactory sensory neurons^60,61^, suppressed their wing flicks and restored the slower dynamics of non-competitive courtship pursuit (**Figures 2M-N**; **Video S7**). The agonistic interactions between competing males therefore appear to be regulated by multiple sensory signals emanating from other males, with song serving a potentially redundant role to olfactory cues.

## Rival males produce an agonistic ‘song’

Agonistic interactions could confer males with a competitive advantage by altering female receptivity, impairing the actions of rival males, or both. To differentiate between these possibilities, we searched a published behavioral database^62^ to identify genetic drivers that trigger bilateral wing flicks, but not unilateral wing extensions. R85A12-GAL4 emerged as a potential candidate as optogenetic activation of neurons labeled by this driver evoked robust wing flicking in males (**Figures 3A-B, S4A, S4C**; **Video S6**). We did not observe wing flicks when expression of R85A12-Gal4 was limited to the brain by genetic intersection, indicating that neurons driving flicks reside within the ventral nerve cord (VNC) (**Figures S4C**; **Video S6**). Consistent with the sexually dimorphic nature of these neurons in the VNC (**Figures 3A, S4A-B**), activation of R85A12 in females also did not produce any overt changes in behavior (**Figure S4C**; **Video S6**).

**Figure 3.**
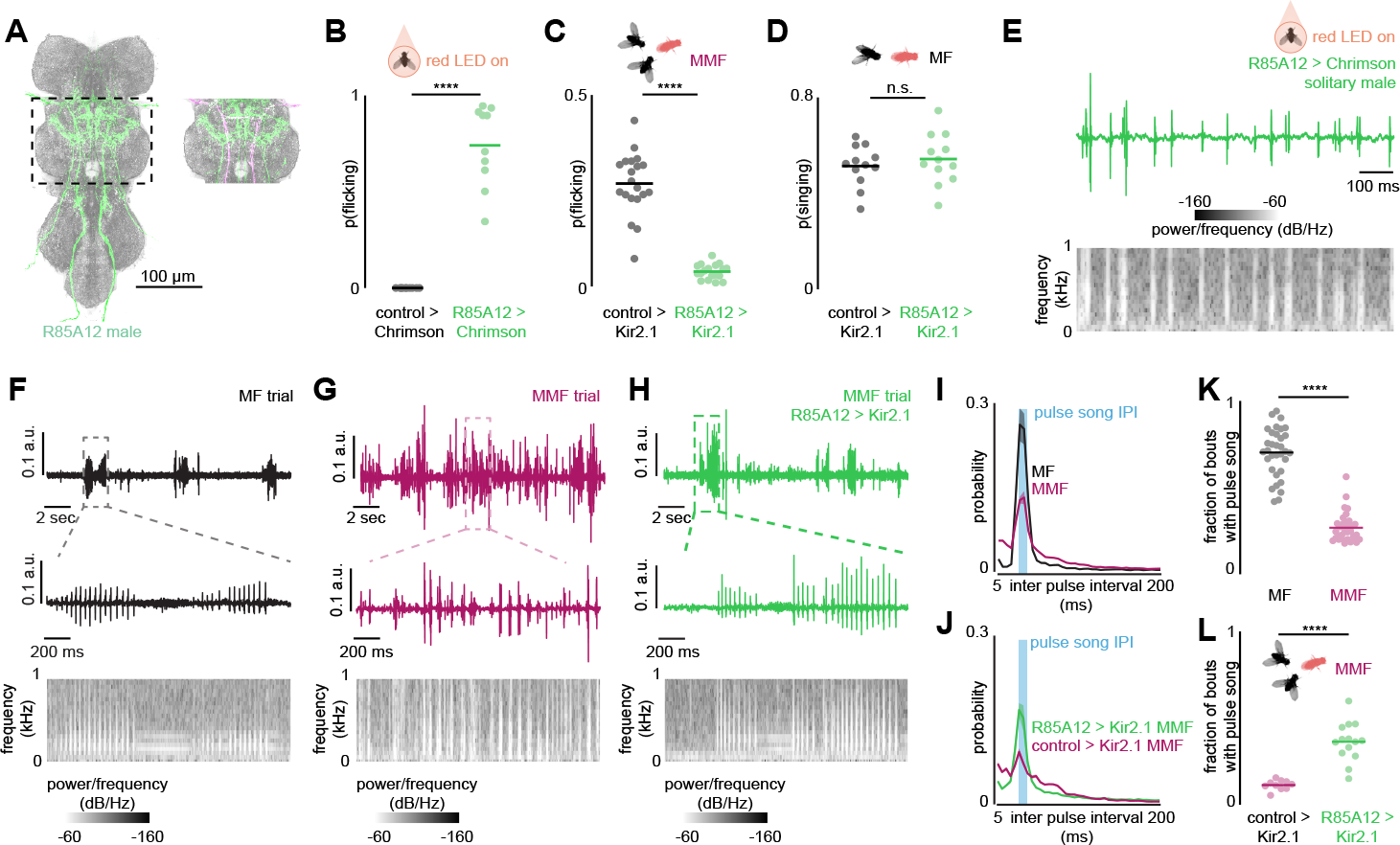
|Wing flicks produce an agonistic song that obscures the courtship song of rivals. (**A**) Morphology of neurons labeled by R85A12-Gal4 in the ventral nerve cord (VNC) of males (left). Neurites with sexually monomorphic expression, common to both sexes, are pseudo-colored by magenta in inset (right). (**B**) Probability of control or R85A12>CsChrimson males performing bilateral wing flicks during red-light-mediated activation. (**C**) Probability of control or R85A12>Kir2.1 males performing bilateral wing flicks in MMF assays. (**D**) Probability of control or R85A12>Kir2.1 males exhibiting unilateral wing extensions in MF assays. (**E**) Representative acoustic recording from a single male during optogenetic activation of R85A12 neurons. Top shows the raw recording trace, bottom shows the spectral decomposition of evoked song. (**F**-**H**) Same as E, but for a wild-type MF assay (F), a wild-type MMF assay (G), and an MMF assay where R85A12 neurons were constitutively silenced via Kir.2.1 expression in both males (H). (**I** and **J**) Distribution of inter-pulse-intervals (IPIs) from acoustic recordings comparing MF and MMF wild-type assays (I), and MMF assays where R85A12 neurons were silenced versus MMF assays with genetic controls (J). (**K** and **L**) Fraction of acoustic bouts classified as pulse song based on their frequency content in MF versus MMF wild-type assays (K) and in MMF assays where R85A12 neurons were silenced versus MMF assays with genetic controls (L). Lines with dots represent mean and individual animals; Shaded lines show mean ± s.e.m.; thin lines denote individual animals. n.s., P > 0.05; ****P < 0.0001. Details of statistical analyses and sample sizes are given in **TableS2**. See also **Figures S4, S5**, and **S7** and **Videos S6** and **S7**.

To confirm that the wing flicks produced by R85A12GAL4 activation represent the same agonistic behaviors performed by competing males, we silenced neurons labeled by this driver in both males of a triad via expression of an inward-rectifying potassium channel. This perturbation largely suppressed the bilateral flicks of males while preserving their ability to court females and perform the unilateral wing extensions that generate courtship song (**Figures 3C-D, S7A-G**; **Video S7**), indicating that these two wing displays are mediated by distinct and segregated motor pathways. Indeed, optogenetic inhibition of the pIP10 descending neuron that mediates production of pulse song^37,63^ suppressed unilateral wing extensions in courting pairs but had no effect on the vigorous flicks evoked by song playback in pairs of males (**Figures S4F-G**).

In many *Drosophila* species, wing displays underlie acoustic signaling^58^. Auditory recordings from isolated males induced to flick via optogenetic activation of R85A12-Gal4 neurons revealed that they produced an audible signal with characteristics that resembled the ‘agonistic song’ that males emit as they compete for food resources^35^ (**Figures 3E, S4D-E**). This agonistic song is distinguished from courtship song by its higher carrier frequency (∼257±22Hz) and longer inter-pulse-interval (∼91±19ms)^35^ (**Figures S4D-E**). Bilateral wing flicks and unilateral wing extensions thus comprise distinct channels for acoustic communication, both of which appear to be almost continuously engaged (**Figure 2C**) during competition for a potential mate.

## Acoustic interference of female receptivity pathways

Given the importance of courtship song in stimulating a female’s sexual receptivity^9,12–17^ (**Figures 1E-F**), we considered whether one role for the incessant flicks performed by competing males may be to complicate the acoustic environment of a female and confound her ability to recognize the song of rival males. To assess this possibility, we compared auditory recordings of MF pairs and MMF triads. In contrast to the typical pattern of alternating pulse and sine song emitted from a courting pair^46^ (**Figures 3F**), the acoustic environment in triads was louder and more complex^64^ (**Figures 3G, S4H-I**), with the characteristic carrier frequencies of the *D. melanogaster* courtship pulse song^46,48,65^ (∼220Hz) largely obscured by a broadband signal centered around the frequency domain of agonistic song (∼275Hz) (**Figure 3G, S5A-C**). Indeed, individual bouts of pulse song^47,48^ (IPI ∼35ms) were frequently contaminated by the presence of higher frequency pulses interspersed throughout each MMF trial (**Figures 3I, S5A)** such that significantly fewer epochs of unadulterated pulse song could be detected during competitive courtship (**Figures 3K, S5D-H**). Acoustic signals in MMF assays that lacked the spectral properties of either pulse or sine song were concentrated around the characteristic frequencies emitted by wing flicks (**Figure S5I**), consistent with the notion that agonistic song is the predominant source of this corruption. Silencing R85A12 neurons in both males of a triad significantly attenuated this broadband signal, largely restoring the distinctive IPI and frequency profile of pulse song recorded from MF pairs (**Figures 3H, 3J, 3L, S5B-C**).

To explore how the complex acoustic environment of triads may impact a female’s mating decisions, we recorded from the auditory pathways controlling vaginal plate openings, a final motor step prior to copulation in which the female offers a male access to her reproductive organs^9,14,66^ (**Video S8**). Vaginal plate openings are governed by a pair of female-specific descending neurons (vpoDNs) that receive auditory input from two parallel auditory pathways — excitatory vpoEN neurons that are selectively tuned to *D. melanogaster* pulse song IPI and inhibitory vpoIN neurons that are sensitive to a broader spectrum of courtship songs^14^ (**Figure 4A**). The balance of this excitatory and inhibitory synaptic input is thought to underlie the selective tuning of vpoDNs and assure that females only become sexually receptive when serenaded by a conspecific male. Consistent with the distinct tuning of these auditory pathways, we found that while vpoENs responded robustly to each bout of pulse song recorded from a MF courting pair, their activity was far weaker in response to acoustic recordings from MMF triads or pure bouts of agonistic song (**Figures 4B-D, S6A-B**). Conversely, vpoINs were preferentially activated by the complex acoustic environment of triads and agonistic song and displayed weaker responses to courtship song (**Figures 4B-D, S6A-B**). The acoustic output of the agonistic wing flicks performed by competing males therefore appears to shift the balance of excitation and inhibition onto vpoDNs (**Figure 4C-D, S6A-B**).

**Figure 4.**
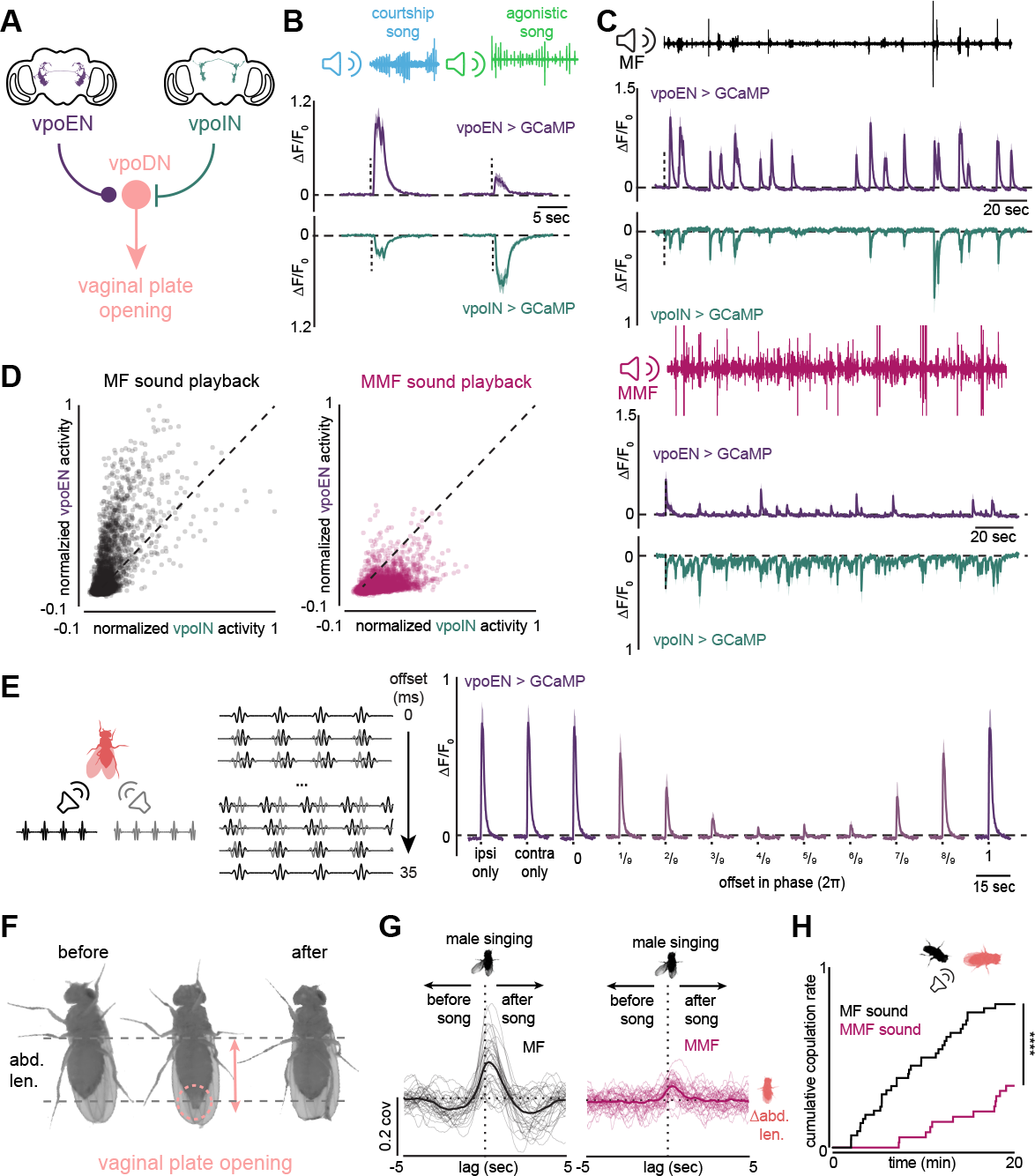
|Acoustic interference of auditory pathways controlling female receptivity. (**A**) Schematic of the excitatory (vpoEN) and inhibitory (vpoIN) auditory input onto the vpoDN descending neurons that control vaginal plate openings in females. (**B**) Average evoked vpoEN (middle) and vpoIN (bottom) activity (∆F/F_0_) during playback of courtship song (left top) or agonistic song elicited by R85A12>CsChrimson activation (right top). Horizontal dashed line traces zero; vertical dashed line denotes stimulus onset. (**C**) Average activity (∆F/F_0_) of vpoEN (purple) and vpoIN (green) neurons in tethered females during playback of acoustic recordings from an MF pair (top) and an MMF triad (bottom). Black and purple traces represent the acoustic recording played back to the female. Horizontal dashed line traces zero, vertical dashed line denotes the onset of sound playback. (**D**) Normalized vpoEN responses and vpoIN responses during playback of MF (left) or MMF (right) acoustic recordings. Normalization was performed across animals; dashed lines denote the unity line; dots denote individual time points from all MF or MMF recordings. Note that the balance of vpoEN and vpoIN responses shifts towards inhibition during MMF playback. (**E**) Schematic representing shifting phase of pulse song stimuli played back to females (left). Brief (1 sec) synthetic D. melanogaster pulse trains were played simultaneously from both the ipsilateral and contralateral side of the female with respect to the imaging ROI, with varying phase offsets between them (middle). vpoEN activity (∆F/F_0_) to pulse song when played from either side alone, and to simultaneous playback of two songs with indicated phase offsets (right). (**F**) Representative snapshots of a female before, during, and after vaginal plate opening following a male’s unilateral wing extension in an MF pair. Dashed lines show average abdomen length. (**G**) Cross-covariance between male unilateral wing extensions and increases in female abdomen length in MF (left) and MMF (right) assays. Positive lags indicate that unilateral wing extension leads in time. (**H**) Cumulative fraction of assays in which the female copulated with a wingless male across 20 minutes when the acoustic recording from an MF pair (black) or an MMF triad (purple) was played back. Shaded lines show mean ± s.e.m.; thin lines denote individual animals. ****P < 0.0001. Details of statistical analyses and sample sizes are given in **TableS2**. See also **Figure S6** and **Videos S8** and **S9**.

Females could potentially differentiate between the acoustic signals emitted by rival males and reduce their susceptibility to interference if they were able to selectively detect the courtship and agonistic song produced from two different spatial sources and process them independently. To assess this possibility, we evaluated the sensitivity of vpoENs to acoustic interference. While males rarely sing synchronously during competitive courtship (**Figure S2C**), we leveraged the stereotyped IPI of pulse song and played bouts of synthetic pulse song to the female from two different speakers, angled at her left or right antenna, while progressively shifting the phase between them (**Figure 4E**). Consistent with the suggestion that higher-order auditory populations can integrate acoustic signals bilaterally^67,68^, vpoEN responses were equivalent when song was played to the female from either speaker or synchronously through both (**Figures 4E**). These auditory pathways thus appear unable to discriminate the spatial origin of a male’s courtship song, aligned with behavioral evidence that females are willing to copulate with a mute male if a rival male is singing nearby (**Figure 1G**) or when song is played back in a delocalized manner through a speaker beneath the chamber floor (**Figure 1F**). However, introducing even small temporal offsets between the songs emanating from the two speakers suppressed vpoEN activity, such that these excitatory pathways were nearly unresponsive when the songs were perfectly out of phase with one another (**Figure 4E**), consistent with the narrow spectral tuning of this population. Auditory circuits controlling vaginal plate opening thus appear vulnerable to acoustic overlap from spatially segregated sound sources.

To explore how acoustic interference from competing males impacts a female’s sexual receptivity at the behavioral level, we compared the ability of a bout of male song to trigger female vaginal plate openings in MF courting pairs and MMF triads, leveraging the fact that vaginal plate openings induce a characteristic lengthening of the female abdomen^9,14,69^ (**Figures 4F, S6C-D**; **Video S8**). When only a single male serenaded a female, we observed a tight correlation between his unilateral wing extensions, the elongation of the female’s abdomen as the vaginal plates opened, and his subsequent copulation attempts (**Figures 4G, S6C-E, S6G; Video S8-S9**). However, this feedback loop between male and female behaviors was strongly attenuated in triads, such that females were significantly less likely to open their vaginal plate in response to a male’s unilateral wing extension in the presence of a competitor (**Figures 4G, S6F-G; Video S9**). Indeed, females took far longer to copulate with a mute male when the complex acoustic environment recorded from triads was played back to her than when the simpler soundscape of a courting pair was replayed (**Figure 4F**). Agonistic wing flicks therefore appear to effectively ‘jam’ the female’s auditory pathways by shifting the excitatory-inhibitory balance onto vpoDNs, obscuring her perception of a male’s courtship song and diminishing her willingness to accept a mate.

## Male-male interactions regulate mating success

Given the constant acoustic interference by competing males, how do males ever succeed in copulating with a female? To gain insight into this question, we analyzed the statistics of the acoustic environment of triads in the minute prior to copulation (**Figure 5A**). We found that there was a sharp increase in the probability of detecting unadulterated pulse song, free from interference by agonistic song, in the two seconds immediately preceding coitus (**Figure 5A**), an epoch during which a male typically performs a final unilateral wing extension before preceding to copulate with the female^50^ (**Figure 1D-E**). Indeed, a bout of pure pulse song was apparent in the majority of triads (84.5%) in the final two seconds, such that it represented the predominant frequency in the power spectrum (**Figures 5B**). Over this same period, the losing male became spatially separated from the copulating pair (**Figure 5C**), suggesting that mating preferentially occurs when one male succeeds in capturing a moment alone with the female and serenades her with a bout of his uncontaminated song, triggering her vaginal plates to open.

**Figure 5.**
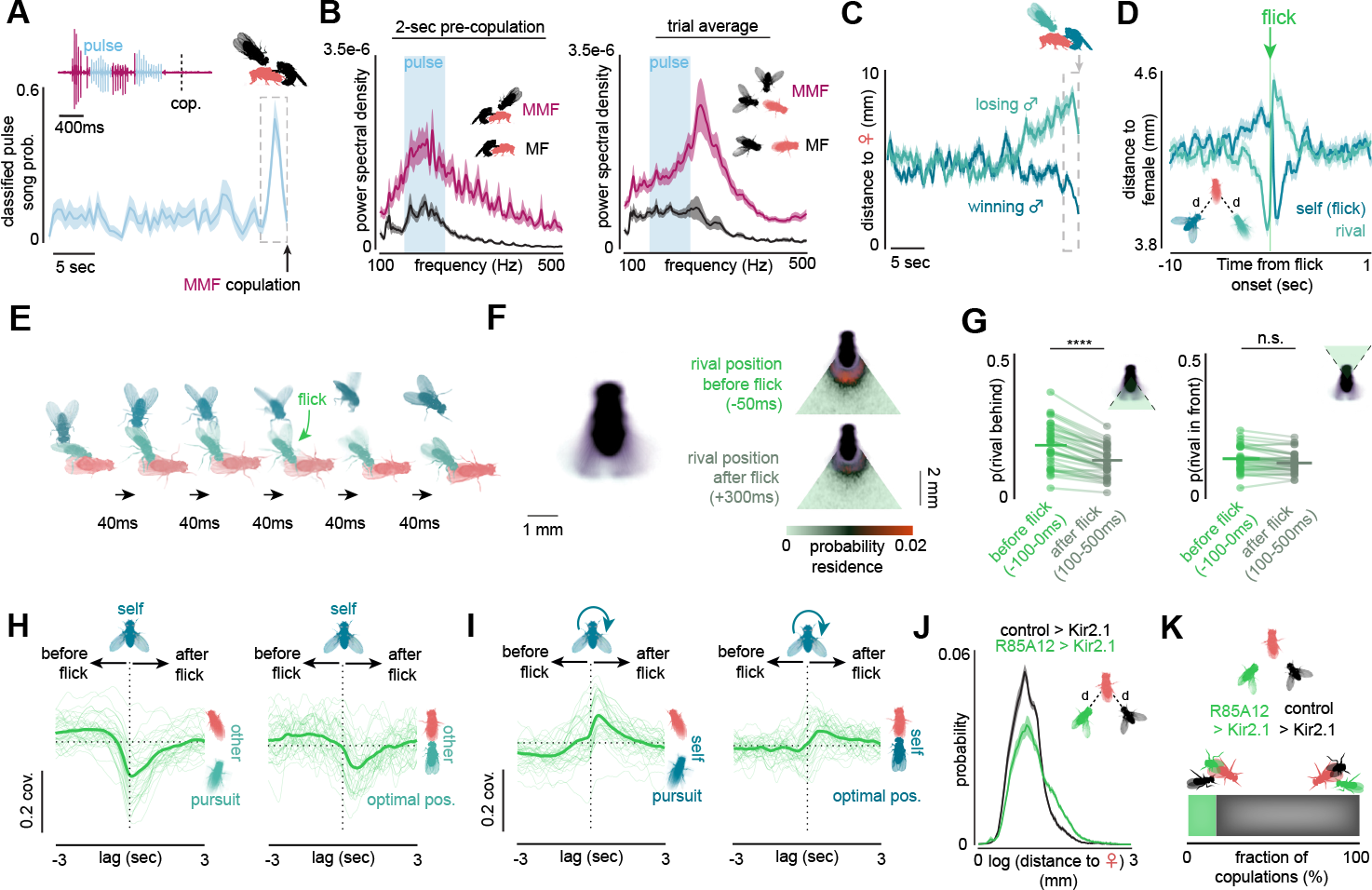
|Agonistic actions contribute to male reproductive success in competitive mating environments. (**A**) Top: A representative example of an acoustic recording from the 2 seconds prior to copulation in a MMF triad. Uninterrupted pulse trains are highlighted in blue. Bottom: Probability of pulse song classified using a frequency-based method (see Figure S5) in the last 30 seconds before copulation in MMF triads. Dashed box highlights the final two seconds before copulation.(**B**) Average power spectra of acoustic recordings in the 2 seconds prior to and 0.5 seconds after copulations (left), or from randomly selected 2.5-second time windows from MF (black) or MMF (purple) assays. Blue regions denote the approximate canonical carrier frequency range for D. melanogaster pulse song (∼220±50 Hz)^46,48,65^ . (**C**) Average distance between the female and the winning (blue) or the losing (green) male across the last 30 seconds before copulation in MMF triads. The dashed box indicates 2 seconds before copulation. (**D**) Average distance of a flicking male (blue) and his rival (cyan) to the female, aligned to the onset of flicking bouts. Only flicks that occurred at least one second after the end of the previous flick were included in analysis. (**E**) Representative behavioral sequence from high-resolution recordings, in which the male in green physically dispels his rival (blue) by performing a wing flick, increasing the distance between the rival and the courting pair. (**F**) Left: Image showing the average of frames in which a male performed wing flicks across one MMF assay (black, n=2295 frames, 301 bouts of wing flicks). Wing-flick motion is concentrated in the quadrant directly behind the male (45° on each side of the male). Right: Heat maps comparing the occupancy of the rival male within the 5-mm quadrant behind the flicking male prior to (50ms, top) and after (300ms, bottom) the central male performed a wing flick. (**G**) Average probability that the rival male resides in the 5mm quadrant behind the male (left), where wing flick motions are concentrated, or in the 5mm quadrant in front of the male (right) prior to (−100 - 0 ms) and after (100-500 ms) the central male performed a wing flick. (**H**) Cross-covariance in MMF triads between one male’s bilateral wing flicks and the other male’s pursuit of the female (left) or the other male’s occupation of the optimal copulatory position behind the female (right). Positive lags indicate that bilateral wing flicks lead. (**I**) Same as (H), but between one male’s bilateral wing flicks and his own pursuit (left) or position relative to the female. (**J**) Distribution of distance from the female to each of the two males in an MMF triad, in which one male is a control male and the other male has R85A12 neurons constitutively silenced by exogeneous expression of Kir2.1 inward-rectifying potassium channels. (**K**) Fraction of all competitive courtship assays (n = 64 triads) in which the female copulated with a male unable to perform wing flicks due to silencing of R85A12 neurons (green) versus a control male (black). Shaded lines show mean ± s.e.m.; thin lines denote individual animals. Details of statistical analyses and sample sizes are given in **TableS2**. See also **Figure S7** and **Video S10**.

Notably, we found that a male’s agonistic actions may facilitate his ability to transiently sequester the female and gain access to these precious mating opportunities^18–20^. Indeed, a male’s performance of wing flicks appeared to rapidly propel his rival away from the female, altering both their relative distances to her in his favor (**Figures 5D-E, S7I**). As a consequence, the onset of a male’s wing flicks was associated with an acute decrease in the probability that his rival would pursue the female and occupy the optimal copulatory position behind her and an increase in the probability that he himself occupied this position (**Figure 5H-I**). These flick-evoked alterations in the spatial relationship of the two males to the female were maintained even when both were deafened (**Figures S7J-L**). Beyond their role as an acoustic signal used to jam a female’s auditory pathways, wing flicks may therefore also directly shape male-male interactions by serving to physically repel competitors. In support of this possibility, a male’s wing flicks could effectively displace a rival encroaching from behind him, where wing motion is concentrated, but not in front of him (**Figures 5F-G**). Wing flicks may thus enhance a male’s chances for reproductive success by physically constraining the ability of opponent males to gain access to the female. Consistent with this notion, while 85A12-silenced males produced pulse song and copulated with females at the same rate as control males in MF pairs (**Figures 3D, S6A-G**), they remained at a distinct disadvantage in triads, spending significantly less time near the female throughout the trial and ultimately losing to control males in most (82.8%, n=64 triads) competitions (**Figures 5J-K, S7H**; **Video S10**). Notably, males that were selectively impaired in their ability to perform agonistic flicks appear to be at a greater competitive disadvantage than mute males that can neither sing nor flick (**Figure 1G**), presumably because wingless males can silently ‘sneak’ in to exploit the courtship song generated by their rivals^58^ without provoking them to become further aroused or countersignal.

## P1a and pC1x neurons coordinate courtship and agonistic actions

Our data suggests that a male’s reproductive success during competition requires that he continually alternate his ardent pursuit of a female with agonistic interactions to fend off his rivals. To gain insight into how males can flexibly pattern their ongoing behavior during competitive courtship, we leveraged the fact that tethered males will spontaneously court a fly-sized visual target projected onto a panorama, faithfully tracking its motion while singing to it ^40,42^ (**Figures 6A-C, S8A**; **Video S11**). We found that playback of pulse song to a tethered courting male was sufficient to provoke him to flick both wings while charging towards the visual target with elevated linear speeds (**Figures 6B-D, S8A-G**; **Video S11**), replicating the agonistic displays performed during competitive interactions with a rival. In contrast to the persistent nature of spontaneous courtship pursuit^39–41,63^, these agonistic behaviors were transient and time-locked to epochs of song playback (**Figures 6B-D, S8A-G**; **Video S11)**. Indeed, in the brief (3 sec) period between bouts of song playback, males resumed tracking the visual target at the slower linear speeds characteristic of courtship pursuit (**Figures 6B, 6D, S8A-G**) while frequently performing unilateral wing extensions (**Figures 6B-D, S8A, S8F**; **Video S11**). Males can thus rapidly interleave courtship and agonistic behaviors towards the same visual target depending on the presence of additional sensory signals, mirroring the flexible alternation of motor displays they perform in the context of mating competitions (**Figure 2D**).

**Figure 6.**
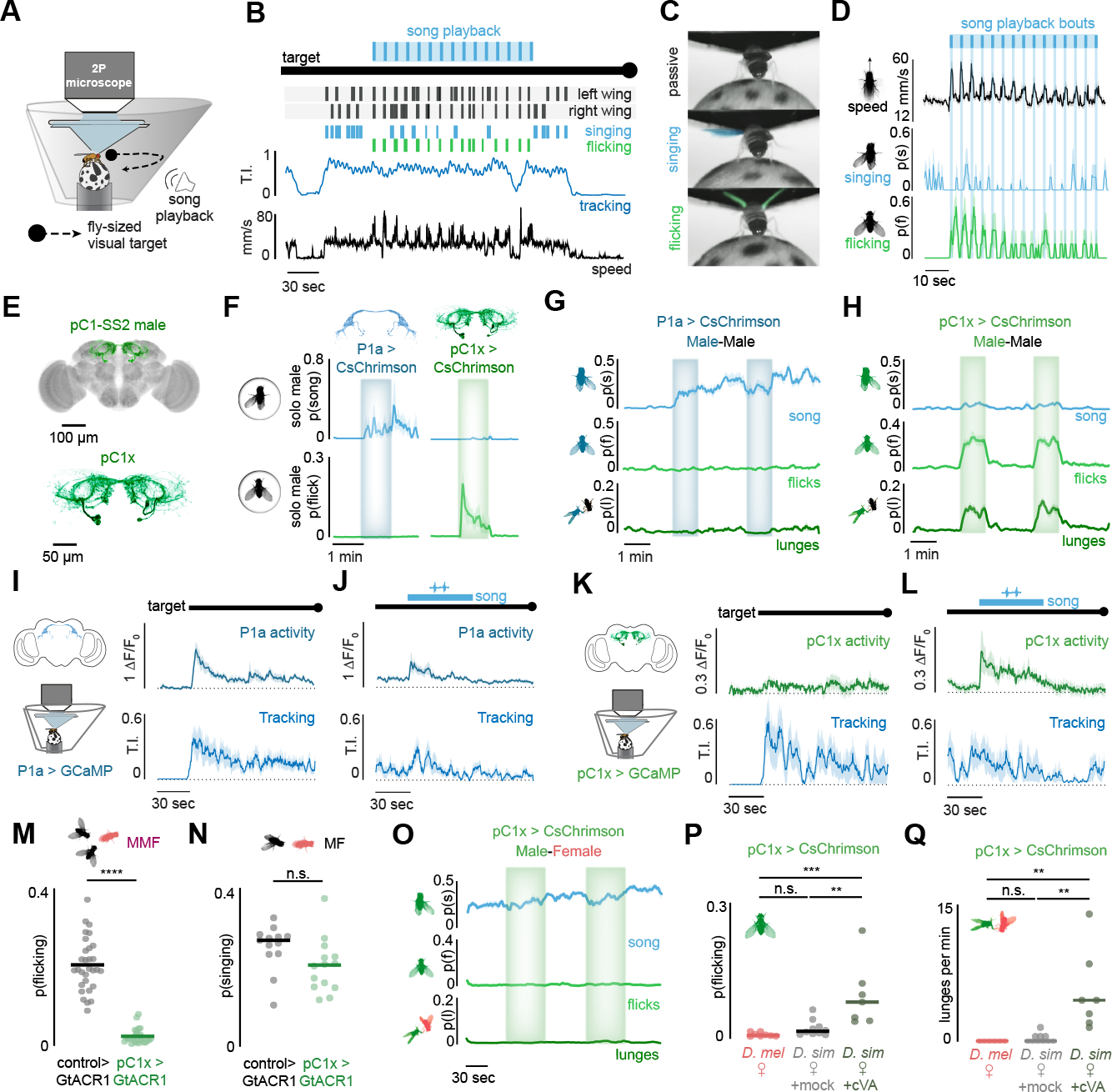
|P1a and pC1x neurons coordinate the expression of courtship and agonistic behaviors. (**A**) Schematic of the tethered behavioral setup, where a fly-sized visual target is projected onto a panorama in front of a headfixed male and oscillates back and forth, while neural activity is recorded via a two-photon microscope. A speaker is positioned near the male for the delivery of synthetic D. melanogaster pulse trains. (**B**) Representative behavior of a tethered male prior to, during, and after the playback of conspecific courtship song. Black hatch marks denote left and right wing extensions; light blue hatch marks indicate bouts of unilateral wing-extensions (singing); green hatch marks indicate bouts of bilateral wing flicks. The male’s tracking index (T.I., see methods) is plotted in blue, and his linear velocity in black as he transitions from courting the fictive female by singing to charging towards it while flicking upon song-playback. Note that wing extensions are interleaved with wing flicks during the playback. (**C**) Pseudo-colored example images of a tethered male viewed from behind, during tracking while not performing any wing behaviors (top), performing a unilateral wing-extension (middle), or performing a bilateral wingflicks (green). (**D**) Average male linear speed (top), probability of males performing unilateral wing extensions (middle), and probability of males performing bilateral wing flicks (bottom) aligned to the playback of pulse trains. Light blue box denotes period of sound playback, darker hatch marks denote individual pulse trains. (**E**) Morphology of neurons labeled by pC1-SS2 in a male’s brain with (top) and without (bottom) background neuropil staining. (**F**) Average probability of solitary males performing unilateral wing extensions (song, blue) or bilateral wing flicks (flick, green) during optogenetic activation of P1a versus pC1x neurons. (**G** and **H**) Average probability of males performing unilateral wing extensions (top), bilateral wing flicks (middle), or lunge attacks (bottom) towards a wingless mute male prior to, during, and after two 1 min epochs of optogenetic activation of P1a neurons (G, shaded regions) or pC1x neurons (H, shaded regions). (**I** and **J**) Average P1a activity (∆F/F_0_, top) and tracking index of tethered males that that courted (wing extensions, tracking) the visual target, aligned to either the onset of visual motion (I; black line on top denotes when the visual target was oscillating) or to the onset of pulse song playback (J; blue line shows period of song playback). (**K** and **L**) Same as I and J, but showing the average activity of pC1x neurons. (**M**) Probability of control males versus pC1x-silenced males performing bilateral wing flicks in MMF triads. (**N**) Probability of control males versus pC1x-silenced males performing unilateral wing extensions when paired with a single female. (**O**) Same as (G and H), but for males paired with conspecific virgin females during optogenetic activation of pC1x neurons. (**P** and **Q**) Probability of males performing bilateral wing flicks (P) and the frequency of lunge attacks (Q) during optogenetic activation of pC1x neurons in males paired with a *D. melanogaster* female (red), a *D. simulans* female (light gray), or a D. simulans female perfumed with the male pheromone cVA (dark gray). Shaded lines show mean ± s.e.m.; Lines with dots represent mean and individual animals. n.s., P > 0.05; *P < 0.05; **P < 0.01; ***P < 0.001; ****P < 0.0001. Details of statistical analyses and sample sizes are given in **TableS2**. See **Figures S8, S9**, and **S10** and **Videos S7, S11, S12, S13**, and **S14**.

Male-specific P1a neurons have been proposed to play a role in promoting both courtship and aggression^39,70,71^ when animals encounter each other on a food source, suggesting they may coordinately control the distinct behaviors that males express towards the female and rival male in a triad. However, in the absence of food, we found that optogenetic activation of this population in males paired with a wingless target male, that was unable to countersignal by flicking, drove persistent courtship pursuit without eliciting any apparent agonistic actions (**Figures 6G, S9D**; **Video S12**). The absence of aggression towards a wingless male parallels observations that P1a neuron activation selectively triggers courtship towards inanimate visual targets^38,40–42^, highlighting the importance of sensory and behavioral feedback cues in shaping a male’s ongoing actions. We therefore searched for additional circuit nodes sufficient to drive wing flicks, focusing on neurons marked by expression of the Doublesex transcription factor due to its role in specifying sexually-dimorphic patterns of aggression^72–76^. We identified a previously described intersectional genetic driver, pC1-SS2^14,77^, that labels four neurons within the central brain as a potential candidate (**Figures 6E, S9A-C**). We call this neuronal subset pC1x.

While pC1x neurons form part of the sexually dimorphic pC1 cluster from which the P1a neurons also derive, optogenetic activation of these morphologically similar populations elicited distinct patterns of behaviors, pointing to divergent roles in promoting courtship and agonistic displays. First, activation of pC1x neurons in isolated males triggered them to perform bilateral wing flicks, in contrast to the sustained unilateral wing extensions evoked by P1a neuron activation in the absence of a visual target (**Figure 6F**). Second, pC1x neuron activation selectively drove males to pursue, flick their wings towards, and even escalate to lunging at wingless male targets (**Figures 6H, S9D**; **Video S13**), indicating that, unlike P1a neurons (**Figures 6G**; **Video S12**), they are sufficient to rapidly evoke aggressive behaviors. Third, while transient P1 neuron activation triggered persistent courtship pursuit^39–41,63,78^ (**Figure 6G**), activation of pC1x neurons evoked agonistic actions only for the duration of the stimulation (**Figure 6H**; **Video S13**), mirroring the acute recruitment of these behaviors during song playback (**Figures 2K, 6B, 6D, S8A-H**). Finally, optogenetic silencing of pC1x neurons in both males of a triad largely suppressed their wing flicks (**Figure 6M**; **Video S7**), with no significant impact on their courtship of a female (**Figures 6N, S9G-K**), supporting they form an essential node that selectively governs the agonistic behaviors that males display in competition.

To assess how pC1x neural dynamics correlated with these transient agonistic actions, we monitored functional responses within their arbors in the lateral protocerebral complex (**Figure S9C**) as tethered males spontaneously initiated courtship and then transitioned to flicking their wings and charging towards the visual target during song playback. While the activity of P1a neurons was tightly correlated with the onset of courtship pursuit^40^ (**Figures 6I, S10A-B, S10H-I**), pC1x neurons responded only weakly as males began to track the target (**Figures 6K, S10C-D, S10K-N**). However, pC1x neurons were robustly activated as males switched from courtship to agonistic displays during song playback (**Figures 6L, S10D**), reinforcing that this circuit node is recruited by sensory cues emanating from rival males to acutely drive competitive behaviors. Notably, pC1x neuron activity was sustained throughout the entire epoch of song playback (**Figures 6L, S10D**) and not just in the brief moments a male performed wing flicks (**Figures S10D, S10J**) suggesting that, analogous to P1a neurons^39–41,63,78^, this central node does not convey acute motor commands to drive behavior (**Figures S10B**,**E**). Moreover, pC1x neurons remained near baseline in males that tracked the visual target but failed to flick during song playback, underscoring that this population promotes the emergence of agonistic actions without directly triggering them (**Figures S10D, S10J-L**).

To gain insight into how the sensory signals emanating from a rival male gate the acute expression of wing flicks, we compared pC1x-evoked behaviors towards different target flies. Optogenetic activation of pC1x neurons in males paired with a conspecific female continued their courtship pursuit without displaying any apparent agonistic actions, suggesting that females either convey cues that suppress agonistic behaviors or lack male cues that promote them (**Figures 6O-Q, S9D**; **Video S13**). Notably, *D. simulans* females rarely elicited aggression upon pC1x activation (**Figures 6O, S9E**; **Video S14**), suggesting that they are not interpreted as rival males despite that they carry the same cuticular pheromones as *D. melanogaster* males^79,80^. However, perfuming a *D. simulans* female with the male-specific olfactory pheromone cis-vaccenyl acetate^54,81–84^ (cVA) was sufficient to transform her into a perceived adversary, such that pC1x-activated males flicked their wings and escalated to lunging at her (**Figure 6P**,**Q, S9F**; **Video S14**). Shortrange olfactory pheromones thus appear to contribute to sex-recognition^54,75,83,85,86^, acting in parallel to song, to rapidly tune a male’s social interactions by gating the aggressive motor programs promoted by pC1x neurons.

The ability of P1a and pC1x neurons to bias males toward opposing social behaviors raises the possibility that they may function in a mutually antagonistic manner, analogous to the cross-inhibitory ‘nodes’ from classic ethology that control the expression of distinct behavioral programs^87^. Recording from P1a neurons, however, revealed that their activity was not suppressed as males transitioned from courtship pursuit to agonistic displays (**Figures 6J, S10B**). Rather, P1a neuron activity continued to correlate with the intensity of a male’s pursuit^40^ and became even further elevated during periods of song playback (**Figures 6J, S10B**), reflecting the increased vigor of their courtship (**Figures S8B, S8F-H, S10I**). Courtship song therefore appears to coordinately signal both the presence of a prospective mate and rival male, serving to invigorate a male’s arousal, potentially through direct auditory inputs to the P1a neurons^57^, while also acutely recruiting pC1x neurons to trigger brief bouts of aggression. P1a neurons thus remain continuously active as males rapidly interleave courtship and aggressive displays, enabling males to remain aroused as they transiently layer agonistic behaviors onto their ongoing performance of the courtship ritual.

### Discussion

During *Drosophila* courtship, male and female behaviors are tightly intertwined through multiple rapidly unfolding feedback loops: females adapt their behavior in response to male song, while males pattern their song based on female behavior^9–11,13,50,88^. In the wild, however, these courtship rituals typically occur against a more complex social backdrop, where many males vie for the opportunity to mate with the same female^18–20^. Here, we investigated how mating decisions emerge in the context of competitive social environments. Our findings suggest that while courtship song is essential to charm a *D. melanogaster* female and render her sexually receptive, females appear unable to discriminate between competing conspecific mates based on their acoustic performance alone. Rather, antagonistic male-male interactions break the tight feedback loops between a courting pair, shifting the dynamics of courtship towards a “struggle between males for possession of females”^3^. Male-male competition thus appears central to mechanisms of mate choice in *Drosophila*.

To curtail the mating opportunities of their rivals, we find that males interleave their courtship of a female with the production of agonistic wing flicks. These wing flicks emerge selectively in competition and enhance a male’s opportunities for reproductive success by thwarting the ability of his opponents to gain optimal access to a female’s genitalia, likely through a direct physical engagement. In parallel, the performance of wing flicks also generates an agonistic song that exploits several characteristics of the circuits controlling a female’s acceptance of a sexual partner to obscure her perception of other potential mates. Indeed, while the strict spectral tuning of female auditory pathways is essential for species discrimination and assures females only become receptive in response to conspecific song^14^, it also renders these pathways highly susceptible to acoustic interference. This vulnerability is further exacerbated by the fact that while the peripheral detection of sound waves by the fly’s antenna is thought to be inherently directional^60,89^, vpoENs appear unable to segregate the acoustic signals emanating from different spatial sources, preventing a female from selectively attending to the courtship song from a single male among many. As a consequence, males can readily mask the song of their competitors by corrupting its characteristic spectral content. This signal jamming resembles the competitive strategies observed in many species that rely on acoustic communication for mate choice^90–95^. In turn, males of these species have evolved mechanisms to subvert acoustic competition and achieve reproductive success^90–95^, for example by suppressing the calls of their rivals^94^ or emerging as the first to sing amid a chorus of competing males^95^.

In *Drosophila*, a male may overcome the acoustic interference of his rivals by transiently isolating a female, such that for a moment he is the sole male to serenade her and trigger her sexual acceptance. Males can therefore only effectively compete if they continually balance their courtship and aggressive displays, rapidly alternating between the agonistic wing flicks necessary to repel their rivals and the continued courtship pursuit essential to persuade a female to mate. The importance of maintaining this balance is underscored by the fact that males that only engage in courtship and are unable to perform agonistic wing flicks are at a competitive disadvantage (**Figure 5K**). Likewise, aggression rarely escalates beyond wing flicks in the presence of a female and never proceeds to the higher intensity components like lunging observed between males in isolation^52,53,86^, suggesting that males must keep their aggression in check to subserve their reproductive goals. In the wild, these male strategies may also be paralleled by female tactics, such as evasive maneuvers or speed changes, that similarly serve to isolate a single male to allow her sample his song.

The ability of a male to persevere in his courtship pursuit is likely even more critical in the context of these natural social settings, to prevent yet other males from swooping in during extended bouts of fighting. Thus, while females do not appear to directly compare competing males, by relying on male-male interactions for access to mating opportunities, they may nevertheless be indirectly selecting for the most vigorous and persistent sexual partners who can remain resilient in their pursuit while dispelling their rivals. Indeed, while females in the wild could potentially decamp when ‘harassed’ by multiple suitors, we found that females in larger arenas often remain on a dense food patch while being pursued by several males until copulation (**Video S15**). The long-lasting internal arousal state triggered by P1a neurons^39–41,63,78^ may therefore have been reinforced under sexual-selection, not merely to satisfy females, but to facilitate mating within crowded competitive environments.

Our work sheds light on how the circuit architecture underlying *Drosophila* social dynamics tightly regulates the performance of courtship and agonistic actions. Our data reveal how two parallel higher-order circuit nodes – P1a and pC1x neurons – regulate distinct sensorimotor pathways controlling singing and flicking, and appear to work in concert to pattern a male’s ongoing behavior. Such a modular circuit organization may allow for the differential expression of each behavior in other social contexts. Males, for example, perform wing flicks not only when competing for access to mates but also in the acquisition and defense of food^35,52^, raising the possibility that pC1x neurons may be recruited independently of P1a neurons through a distinct set of multisensory signals. Likewise, P1a neurons can signal the intensity of a male’s courtship drive even in the absence of competition, assuring that males faithfully pursue a female without the expression of aberrant agonistic actions. Sexually-aroused males nevertheless appear to be prone to aggression^39^, suggesting P1a or other courtship promoting neurons may indirectly impinge on pC1x neurons or their downstream targets, priming males to defend their mating opportunities. pC1x neurons likely also interface with other previously described aggression-promoting pathways to tune a male’s agnostic behaviors^47,75^.

By recording from tethered males as they engage with a fictive target fly, we reveal that P1a and pC1x neurons remain co-active as they rapidly alternate between courtship and agonistic displays. Thus, while singing and flicking appear to be mutually exclusive motor programs, the circuit nodes gating the expression of these behaviors are not and can be simultaneously engaged to orchestrate a male’s social interactions on a moment-to-moment timescale. Such a circuit logic suggests how males can sustain their sexual arousal even while flexibly transitioning between courtship towards a female and agonistic displays towards a rival male. Indeed, we find that while a fly-sized moving visual target is inherently perceived as a female and sufficient to arouse a male to court^38,40,96^, additional sensory cues are required to gate the expression of sex-specific aggressive displays^54,75,83,85,86^. For example, hearing the song of a rival both activates P1a neurons to invigorate a male’s sexual arousal and enhance the fidelity of his continued pursuit but also temporarily transforms the same visual target into an adversary, driving the acute engagement of pC1x neurons to trigger transient bouts of agonistic behavior. The low volatility male-specific pheromone cVA appears poised to further sculpt a male’s social interactions, acting redundantly with song. A male will not, for example, attack a target fly in the absence of cVA, even with pC1x neurons exogenously activated, underscoring how once a male is aggressively aroused, he may rely on the short-range nature of this pheromone to spatially localize which member of a triad is his rival^97^ and appropriately direct his antagonistic actions. The neural circuits mediating a male’s social interactions therefore function like a switchboard, allowing the same incoming visual information to be flexibly rerouted to different motor circuits to trigger singing or agonistic flicks depending on ongoing changes to his motivational state and immediate sensory context.

The dynamic interplay between individuals in the triad makes it challenging to disentangle how the discrete actions of any individual lead to mating success or to assign unique agency to any member of the group. Indeed, taken together our data suggest that mate choice in *Drosophila* does not simply reflect a ‘decision’ by the female to actively sequester her preferred suitor, the superior ability of the winning male to repel his rival, or a brief lapse by the losing male that provides his opponent with a window of opportunity. Rather, mate choice appears to arise from a distributed and intricate web of social interactions that dynamically unfold across the group, reflecting how female preferences and male-male interactions coordinately orchestrate reproductive strategies.

## ACKNOWLEDGEMENTS

We thank D. Anderson, D. Stern, B. Datta, L. Luo, G. Rubin, B. Noro, C. Schretter, P. Brand, and R. Coleman for comments on the manuscript; D. Anderson, Y. Aso, A. Clark, B. Dickson, V. Jayaraman, G. Maimon, J. Pool, M. Reiser, G. Rubin, D. Stern, the Janelia Fly Bank, and the Bloomington *Drosophila* Stock Center (NIH P40OD018537) for providing fly stocks; S. Sawtelle and D. Stern for generously sharing their designs for an integrated, parallelized acoustic recording system for acoustic communication in *Drosophila* in advance of publication, and G. Maimon and J. Weisman for sharing their designs for an integrated, projector-based system for presenting tethered, walking flies with visual stimuli; G. Rubin, C. Schretter, and C. Christoforou and G. Ihrke of the HHMI Janelia Project Technical Resources Group for generously assisting with immunohistochemical labeling and imaging of R85A12-GAL4 neurons; J. Zhao for performing preliminary experiments characterizing the behavioral effects of pC1 activation; A. Otopalik, R. Coleman, P. Brand, Y. Chiba, A. Kamikouchi, J. Petrillo, and A. Siliciano for technical advice; and L. Abbott, D. Anderson, C. Bargmann, W. Freiwald, G. Maimon, P. Rajasethypathy, G. Rubin, C. Schretter, and members of the Ruta laboratory for invaluable insights and discussion. This work was supported by an NIH NINDS grant (5R35NS111611) and the Simons Foundation Collaboration for the Global Brain to V.R., a David Rockefeller Fellowship and a National Science Foundation Graduate Research Fellowship (1946429) to T.H.S, and a Boehringer Ingelheim Fonds Ph.D. Fellowship to F.H. V.R. is a Howard Hughes Medical Institute Investigator.

## AUTHOR CONTRIBUTIONS

R.L., T.H.S., and V.R. conceived the project, designed experiments, interpreted the results, and co-wrote the manuscript. F.H. collected data for Figure 6F and contributed to preliminary behavioral data collection related to pC1x. S.E. contributed preliminary behavioral data related to Figure 6 and S4. R.L. and T.H.S were equal contributors and are listed alphabetically.

## COMPETING INTERESTS

The authors declare no competing interests.

## OPEN ACCESS STATEMENT

This article is subject to HHMI’s Open Access to Publications policy. HHMI lab heads have previously granted a nonexclusive CC BY 4.0 license to the public and a sublicensable license to HHMI in their research articles. Pursuant to those licenses, the author-accepted manuscript of this article can be made freely available under a CC BY 4.0 license immediately upon publication.

## RESOURCE AVAILABILITY

### Lead contact

Further information and requests for resources and reagents should be directed to and will be fulfilled by the lead contact, Vanessa Ruta.

### Materials availability

This study did not generate new unique reagents.

### Data and code availability

Data: Raw behavior videos, sound recordings, and calcium imaging data will be provided upon request from the lead contact.

Code: Data analysis code is available from the lead contact upon request.

Additional information: Any additional information required to reanalyze the data reported in this work is available from the lead contact upon request.

### Experimental model and subject details

Flies were housed under standard conditions at 25°C on a 12h light-dark cycle, except for flies for optogenetic experiments (i.e. expressing CsChrimson or GtACR1 and their controls) which were dark-reared. All flies were raised on molasses food unless otherwise noted. Male flies used for imaging experiments were raised on Würzburg food^98^ for at least five generations to increase behavioral robustness. All crosses for optogenetic experiments were grown on sugar-yeast food (100 g Brewer’s yeast, 50 g sucrose, 15 g agar, 3 ml propionic acid, 3 g p-hydroxy-benzoic acid methyl ester per 1 Liter H2O) to avoid low levels of retinal metabolized from vitamin A in more nutritious food. Progeny from these crosses were also raised on sugar-yeast food before flies were transferred to food containing 400μM all-trans-retinal (Sigma-Aldrich R2500) at least 48h before the experiments.

Fly stocks used were as follows: Canton-S, UAS-Kir2.1 (G. Maimon, The Rockefeller University), CO-4N (J. Pool, UW-Madison), ZH42 (A. Clark, Cornell University), UAS-CsChrimson-mCherry (B. Dickson, Janelia), Ruta Lab *D. simulans*, Otd-nls:FLPe, Split-P1a (D. Anderson, Caltech), UAS>STOP>CsChrimson (M. Rieser, Janelia), 2xUAS-jGCaMP7f (Y. Aso and V. Jayaraman, Janelia), split-pIP10 (D. Stern, Janelia). The following stocks were obtained from the Bloomington Drosophila Stock Center: UAS-CsChrimson-mVenus (BDSC: 55134), UAS-GtACR1 (BDSC: 92983), UAS-jGCaMP7s (BDSC: 79032), Gal4 control (BDSC: 68384), Split Gal4 control (BDSC: 79603), R85A12-Gal4 (BDSC: 47967). vpoIN-SS2, vpoEN-SS2, vpoDN-SS1, DNp13-SS1, and SS59911 were obtained from the Janelia Fly Bank.

Key resources table and **TableS1** provide detailed descriptions of all genotypes used in each experiment. The age, sex, and additional housing conditions of the flies are described in the method details.

## Method details

### Free behavioral assays

All assays were performed with virgin male and virgin female flies 3-6 days post-eclosion. Flies were isolated 2-8 hours post-eclosion and reared with flies of the same sex at low density (5-10 animals) in food vials (d = 3cm, h = 9cm), except that males in wild-type behavioral assays and tethered assays were raised in isolation. For amputation experiments (e.g., **Figures 2M and 2N**), aristae or the third segment of antennae were surgically removed by a needle (BD PrecisionGlide Hypodermic Needles 18G x 1” BD 305195) connected to a syringe.

Amputations were performed at least 12h before behavioral experiments to allow flies to recover. Free behavior experiments were performed in custom-milled Delrin chambers (d = 20mm, h = 2.5mm or 3.5mm, protolabs), or custom 3D-printed (printer: ProJet MJP 3600, material: VisiJet M3 Crystal) perforated chambers (d = 20mm, h = 2.5 or 3.5mm) for efficient sound transmission. The floor of the perforated chamber was punctured at regular intervals by 200μm holes, yielding a smooth mesh that flies appeared to walk comfortably on. All assay chambers were designed with sloped edges to decrease the chances of flies walking on edges or both sides^99^. A thin layer of transparent acrylic board was used as a lid for the chamber. We added flies to the chamber by aspiration without anesthetization.

#### Non-optogenetic free behavioral assays

For characterization of male singing behavior prior to copulation in male-female assays (**Figure 1E**), wing-wingless competition assays (**Figures 1G, 1H, 1I**, and **S1D**), and male-female distances before copulation (**Figure 5C**), we recorded fly behaviors for 1h or till copulation at 40 frames/s. A light pad (Logan Electric Slim Edge-Light Pad A-5A, 5400K, 6 klx) was used for backlighting and videos were taken through a FLIR Grasshopper3 camera (GS3-U3-32S4M) and lens (Fujinon HF12.5SA-1). The video resolution was around 17.5 pixels/mm, sufficient for individual position tracking. No assays were excluded from copulation latency experiments.

For R85A12 silencing competition assays (**Figure 5K**) and copulation latency assessments (**Figure S7G**), we recorded fly behaviors for 1h or till copulation at 40 frames/s. White LED strips (HitLights L0512V-402-1630-U) were used for backlighting and videos were collected via a FLIR Grasshopper3 camera (GS3-U3-41C6C-C) and lens (Fujinon CF16ZA-1S) or a FLIR Grasshopper3 (GS3-U3-32S4M) and lens (Fujinon HF12.5SA-1). For competition assays, we randomly painted either R85A12-silenced or control males with a white dot on the dorsal side of their thoraces with acrylic paint (Magicfly) for tracking their identities, at least 12h before behavioral experiments to allow them to recover. We did not detect any correlation between painting and copulation success in competition assays (data not shown). A subset of competition assays was collected via high-resolution recording (see below) till copulation to allow us to better quantify behavioral patterns (**Figures 5J, S7H**).

All other assays were performed in upside-down conditions with high resolution for postural tracking, where the sloped chamber was used as the top and the acrylic lid was used as the bottom. We painted the chamber with Fluon (byFormica) to constrain flies to walk on the transparent acrylic side. White LED strips (HitLights L0512V-402-1630-U) were used for backlighting. A FLIR Flea3 camera (FL3-U3-32S2M-CS) with lens (Computar A4Z2812CS-MPIR) was placed under the bottom side to record behaviors from the ventral side of flies, enabling us to robustly and consistently quantify female abdomen lengths. Unless otherwise noted, we recorded the assays for 20 minutes or till copulation at 60 frames/s. The video resolution was around 72.5 pixels/mm. For wild-type high-resolution behavioral assays, assays with copulation latency longer than 20 minutes or shorter than 45s were discarded to ensure that females were receptive and that both males had time to participate in the chase. No assays were excluded from copulation latency experiments.

#### Optogenetic free behavioral assays

For optogenetic free behavioral assays, white LEDs intensities were turned much lower through a dimmer to minimize aberrant excitation of opsins. IR LED strips (850nm, Waveform Lighting 7031.85) were used for camera recording. High-powered red (627nm, LuxeonStarLEDs SP-03-D9) and green (530nm, LuxeonStarLEDs SP-03-G4) LEDs were controlled through an Arduino board (Arduino UNO REV3 [A000066]). Custom MATLAB scripts synced the video recording, LEDs paradigm, and sound delivery.

For most experiments, videos were taken from a FLIR Grasshopper3 (GS3-U3-32S4M) and lens (Fujinon HF12.5SA-1) at 60 frames/s. The video resolution was around 40 pixels/mm. A Near-IR bandpass filter (MidOpt BP850-40.5) was used to ensure constant lighting for the recording. LEDs were placed above the chamber. To view dynamic changes in fly behaviors spurred by optogenetic activation or inactivation, we typically designed red or green LED paradigms as interleaving 2-minute off and 1-minute on blocks (8min in total) unless otherwise noted. For Male-Female assays, we interleaved 1-minute LED-off and LED-on blocks (5min in total) to allow us to sufficiently sample behavioral patterns before copulation occurred. For Optogenetic Male-Female assays, assays with copulation before the full 5 minutes were discarded. pIP10 silencing experiments were performed under green LEDs with power intensities of 25.2μW/ mm^2^. pC1x activation experiments in social contexts were performed under red LEDs with power intensities of 4.7μW/mm^2^. To estimate the baseline behaviors elicited by activation of P1a neurons or pC1x neurons, solitary males were placed in free-behavior assay chambers, and red LEDs with power intensities of 5.6 μW/ mm^2^ and 11.8 μW/mm^2^, respectively, were used for activation. Assays with solitary males were 3 minutes long, and consisted of a 1-min LED-ON block flanked by two control blocks where the LED was turned off.

For R85A12 activation, pC1x silencing, and pIP10 silencing Male-Female experiments, assays were performed upside-down as described above in the *non-optogenetic free behavioral assays* section. For R85A12 activation experiments, a piece of blue filter paper (Rosco #071 Tokyo Blue) over the camera was used to ensure constant light intensity for the recording. The red LED paradigm was designed as interleaving 5s-on and 5s-off blocks (65s in total). Experiments were performed under red LEDs with power intensities of 5.6μW/mm2, except that for activation in females and expression-restricted males, activation was also performed under red LEDs with higher power intensities (27.6μw/mm^2^) to exclude the possibility that we didn’t observe behavioral phenotypes because of insufficient light power. For pC1x silencing and pIP10 silencing Male-Female experiments, assays were performed under constant green light. pC1x silencing experiments were performed under green LEDs with power intensities of 28-33μW/mm^2^. Because the accumulated heat from high-powered LEDs could interfere with fly behaviors, assays were only recorded for 5 minutes from the start of all males being engaged in courtship. Assays in which flies copulated before the end of the assay were discarded. pIP10 silencing experiments were performed under green LEDs with power intensities of 13.5-18μW/mm^2^ and recorded till copulation. Assays with copulation latencies longer than 20min were discarded.

#### Sound playback free behavioral assays

For the playback of synthetic courtship songs, sound patterns were generated by custom MATLAB scripts. For *D. melanogaster* pulse song, we generated trains of Gaussian-modulated sinusoidal RF pulses with a carrier frequency of 200Hz and 60% bandwidth separated by 35ms (IPI)^16^. For *D. simulans* pulse song, we instead used a carrier frequency of 320Hz and separated pulses by 62ms^16,48^. White noise was generated pseudorandomly from a standard normal distribution. For natural courtship songs, we selected MF and MMF sound recordings from the start of all males being engaged in courtship till copulation. For consistency, we selected MF and MMF recordings that were both approximately 3 minutes long.

Sound was delivered via an 8-inch woofer (Peerless by Tymphany 830869 8” Nomex Cone HDS) placed directly under the experimental chambers at around 10cm distance, and amplified through an amplifier (Pyle PMSA20 50 Watt). For examining male wing flicks in response to playback (e.g., **Figures 2K, 2L**, and **S3**), we interleaved 1-second trains of continuous playback with 3-second periods of silence. Each trial was 5-minute long and the sound was turned on between minutes 2-3, except for the pIP10 silencing experiment (**Figure S4G**) in which each trial was 8-minute long, the sound was turned on between minutes 2-3 and 6-7, and green LEDs were turned on solely between minutes 6-7, allowing us to compare the effects of sound playback depending on whether pIP10 neurons were silenced within animals. For wingless Male-Female copulation latency assessment with synthetic song playback (**Figure 1F**), we interleaved 1-second trains of continuous playback with 2-second periods of silence for 1h. For wingless Male-Female copulation latency assessment with natural songs playback (**Figure 4H**), we looped the MF or MMF recording for 20 minutes. The sound volume for each stimulus was consistent across trials.

#### Perfuming assays

For perfuming experiments (**Figures 6P-Q, S9E**-**F**), we dissolved cis-vaccenyl acetate (cVA, Cayman Chemicals 10010101) in 100% ethanol to yield a concentration of 50μg/ml. We isolated virgin *D. simulans* females and housed them with same-sex flies for 2-3 days before experiments. We subsequently applied 1μL of the cVA solution (or plain ethanol for mock-perfuming) to the abdomen of CO_2_ anesthetized females, and let them recover in food vials for at least four hours before pairing them with pC1x>CsChrimson males. We observed some heterogeneity in the vigor with which males attacked perfumed *D. simulans females*, likely due to uneven distribution of the poorly-soluble cVA across their abdomen.

### >Free behavior analysis

#### Tracking

For wing-wingless competition assays (**Figures 1H**-**1I**), and male-female distances before copulation assays (**Figure 5C**), we tracked the positions of each fly in the last 30 seconds before copulation with Caltech Flytracker^100^.

For other behavioral assays, we trained two DeepLabCut 2.2/2.3rc3 models^43,101^, one for upside-down high-resolution videos and one for perforated chambers to track the postures of interacting flies. Specifically, we manually labeled eight body markers (head, thorax, top of the abdomen, left and right sides of the abdomen, genitalia, and the left/right wingtips). For model 1, we labeled 295 frames taken from 25 videos (95% was used for training). For model 2, we labeled 478 frames taken from 16 videos (95% was used for training). We used the ResNet-50 network architecture with default parameters for 200,000 training iterations. We validated with 1 shuffle. Model 1 exhibited a test error of 3.18px and a train error of 6.78px (image size: 1552×1552px); Model 2 exhibited a test error of 3.19px and a train error of 4.09px (image size: 1088×1088px). We then used a p-cutoff of 0.6 to condition the X and Y coordinates for future analysis. Animals were reconstructed from body parts using the ellipse tracking method. We subsequently manually inspected each video and corrected identity swaps that occurred (typically 2-3 per 36,000 frames). For wild-type MMF assays, we corrected all identity swaps for the comparison between winners and losers. For the rest of the MMF assays, we only corrected male-female identity swaps as the behaviors of both males were pooled in the following analysis.

We also *post hoc* filtered out aberrant single-frame ‘jumps’ in the position of any body part larger than 200 pixels, which most likely represent detection errors in which the body part of one animal is attributed to another. We computed various relationships between the body parts of individual animals, including the centroid X and Y position as the average density of marked points, the abdomen length as the distance between the marker placed at the top of the abdomen and the marker placed at the genitalia, the heading direction as the angle of the head with respect to the thorax, the wing angles as the angle of the wing markers with respect to the thorax, as relative to the male’s heading direction. All initial analysis was performed in Python 3.8, with SciPy v1.7.3, Pandas v1.3.5, and NumPy v1.21.2, and subsequently exported as CSV files for further analysis in MATLAB (2019b, 2021b-2023a).

#### Classification

For male actions before copulation in male-female assays (**Figure 1E**) and wing-wingless competition assays (**Figure 1H**), we manually labeled each male’s wing extension bouts in the last 30 seconds before copulation because of low-resolution recordings. For R85A12 activations in single animals (**Figures 3B** and **S4C**), we manually labeled all wing flicks events for efficiency. We also manually scored vaginal plate opening/abdomen lengthening events in **Figure S6D**.

For other behavioral assays, we imported the parameters extracted from our DeepLabCut postural tracking into the Janelia Automatic Animal Behavior Annotator (JAABA^44^) using custom scripts and a custom parameter configuration file. To train classifiers, we labeled example epochs of behaviors and trained classifiers for singing, flicking, copulation attempts, pursuing females, pursuing males, general pursuit, directed approaches, occupation of the optimal copulatory position immediately behind the female, and lunges (**Figure S1A**). We further validated that all classifiers exhibited a sensitivity (d’) greater than 2.0 in both MF and MMF assays by labeling a ground-truth dataset consisting of 20,000 frames for each behavior in each condition (MF and MMF) from two videos the model did not see during training for each condition (**Figure S1B**). These behaviors were classified by JAABA classifiers without duration thresholds unless otherwise noted. For quantifying the absolute number of behavioral events rather than rates (**Figure 6P-Q**) we manually verified all predicted behavioral epochs. To quantify the frequency of specific male component behaviors over time (e.g., **Figures 1C, 6G-H, S9D-F**), we binned the binary classifier results over time (∆t = 0.25-2.5sec).

#### Co-variance analysis

To understand how the onset of one behavior influences, and is influenced by, the actions of different individuals, we computed the cross-co-variance between the time series of behavioral signals

Hindmarsh Sten and Li *et al*., 2023 21 or spatial relationships between individuals. As behavioral patterns unfold very rapidly, we used a maximum lag of 300 frames (5 seconds) in our analysis and normalized the cross-covariance values by the autocovariance of the source signal at zero lag. During analysis, all missing frames were filled by linear interpolation. To compute covariance matrices between the behaviors performed within and across males (**Figure S2G**), we computed the maximum magnitude (positive or negative) of the cross-covariance within one second of the source behavior.

To validate the fidelity of covariances between a subset of behavioral features across trials and animals (e.g., **Figures S2H-M, S6E-G, S7I**) we computed triggered averages aligned to the onset of specific actions (e.g., singing or flicking). To estimate the frequency at which males approached the female after his rival performed a unilateral wing extension (**Figures S2H-K**), we selected song bouts performed by the rival when the affected male was classified as disengaged from courtship in the second preceding onset of song (accounting for ∼11% of all unilateral wing extensions), and computed the distance and facing angle of the affected male relative to the female. To ensure that the selected unilateral wing extensions were not the result of erroneous detections, we only included wing extensions that lasted for at least 250ms. Moreover, to ensure that detected wing extensions were not split into multiple bouts by dropped frames and instead constituted the onset of a new behavior, we only included wing extensions that occurred at least 1 second from the last detected wing extension. Heatmaps of distance-to-female and facing angles on individual trials were sorted by a modulation index, computed as the average value in the second before the onset of rival song minus the average value in the second after the onset of rival song, divided by the total average (i.e., (Pre – Post)/(Pre + Post)). Similarly, to compute fidelity with which the onset of rival song triggered males to perform bilateral wing flicks (**Figures S2L-M**) and changes in the abdomen length of males and females following song (**Figures S6E-G**), we selected song bouts that lasted for at least 250ms and were separated from previous bouts by at least 1 second, but included bouts that occurred when both males were engaged in courtship. Finally, to compute changes in the distance between flicking males, rival males, and the female relative to the onset of flicks (**Figures 5D, S7I**), we selected flicking bouts that lasted at least 125ms and were separated from pervious bouts by at least 1 second.

#### Wing angle analysis

To characterize the change in wing angle occurring at the onset of flicking and singing, respectively, we defined “wing 1” as the dominant wing that exhibited a larger open angle with respect to the central body axis, and “wing 2” as the subordinate wing (**Figures 2E** and **2F**). For each detection instance, we normalized the wing angle by its position immediately before the onset of behavior.

#### Relative position analysis

We defined the distance between two flies as the distance between their centroids. To visualize a female’s position relative to the male (**Figures S2D, S7F**, and **S9K**), we transformed the female’s position at each frame into a coordinate system defined by the male’s heading direction and position, such that she was located at 90° when faced by the male. Likewise, to estimate the position of rival males immediately before and after a wing-flick was performed, we computed the position of the rival male based on the coordinate system defined by the flicking male’s centroid and heading direction. For heatmaps of the spatial position of rivals or the female, we binned the date into 0.25mm x 0.25mm bins.

### Sound recording and analysis

#### Sound recording

For all sound recordings, we used a high-throughput multi-channel recording apparatus generously shared by D. Stern and S. Sawtelle (personal communication, manuscript in preparation). Briefly, the apparatus consists of a backplane, multiple eight-channel microphone boards, a battery supply, an AC-DC power supply, and chambers. We used four rows of eight-channel microphone boards, such that 32 assays at most could be recorded at the same time. Microphones were connected via the backplane with a 3.5V battery, and an FPGA board. The FPGA board provides the host computer with the control and data signals to the microphone array via the backplane board. Above each microphone, we placed a small 3D-printed behavioral chamber (top diameter: 10mm, height: 3.1mm) covered by a thin layer of acrylic board (1/16’’ thick) on the top side and a thin layer of mesh (McMaster Carr 9318T21) on the bottom side facing the microphone.

Flies were aspirated into the behavioral chambers without anesthetization, and we recorded acoustic outputs via a custom software interfacing with the FPGA board at 5kHz. In parallel, we also recorded video of the behavioral chambers using a FLIR Flea3 camera (FL3U3-32S2M-CS) and lens (Computar A6Z8516CS-MP). Acoustic outputs and videos were synced through a GPIO cable. All assays were recorded for 20 minutes unless otherwise noted. We then manually scored the onset of copulation and assays with copulation latencies longer than 20 minutes were discarded.

For R85A12 activation recordings, a bifurcated fiber-optic cable with Ø1.2mm ferrules (Thorlabs BFYL4LS01) connected to a fiber-coupled LED (625nm, ThorLabs M625F2) was directed at the chambers. A T-Cube LED driver (ThorLabs LEDD1B) controlled LEDs, and we titrated the power output to maximize the number of flicks males exhibited per minute. Each assay lasted 10 minutes and we used the first 5 minutes for analysis.

#### Signal identification in recordings

Sound energy was calculated by estimating the signal envelope using the Hilbert transformation^102^, and we subsequently segmented acoustic bouts where the sound energy exceeded 0.015 (95^th^ quantile in MMF assays) for at least 30ms. To compute the amplitude of each recording (**Figure S4I**), we averaged the maximum sound energies of all sound bouts.

#### Inter-pulse-interval (IPI) computation

For agonistic song and pulse song in MF assays (**Figures S4D-E, S7A-S7D**), pulse locations were selected based on trained classifiers in Deep Audio Segmenter^103^. Then we computed the differences between neighboring pulse locations to find pulse trains. Considering the potential IPI, only pulses with distances less than 200ms were in the same pulse train. All IPIs in every train were used for further computation. Based on the IPI characterized in this way, large values like ∼200ms, likely from detection errors, would greatly shift the mean values. Therefore, we used the median value of IPI in each trial (**Figure S7A**). Recordings with less than 50 detected IPIs were discarded.

For comparing IPI distributions in different recordings (**Figures 3I** and **3J**), pulses were selected in a in a more stringent manner to filter out background noise present in triads. We first filtered the recording by a bandpass filter of 100Hz-1000Hz and then binned the filtered recording into bouts of 30ms without overlap for pulse selection. Only pulse bouts with maximum sound energy above 0.015 were selected. We chose the peak in each pulse bout as the pulse location. Then we computed the differences between neighboring pulse locations as IPI. Considering the potential IPI, we discarded IPIs shorter than 5ms, which are likely from polycyclic pulses, and longer than 200ms, which are likely from pulses in different pulse trains. Then we computed the distribution of IPIs in each recording.

#### Carrier frequency computation

For agonistic and pulse song characteristics (**Figures S4E, S7C**, and **S7D**), 30ms around each detected pulse peak were defined as pulse segments. Given that the raw spectra of pulses are relatively broad, the peak frequency of the spectrum is an unreliable measure of the pulse carrier frequency. Instead, we defined the carrier frequency of each pulse as the center of mass of the power spectra thresholded at (e-1)^102^.

#### Pulse classification based on frequency

To detect segments of unadulterated pulse song across MF and MMF assays (**Figures 3K, 3L, 5A**, and **S5D-I**), we randomly selected a few manually detected 500ms pulse trains, sine, and agonistic songs and computed the power spectrum of each example by using short-time Fourier transformation. Power spectra from each category were averaged, normalized, and fit a two-term Gaussian model. We subsequently created a pulse filter from the fitted spectra by subtracting the sine and agonist power spectra from the pulse spectrum, such that the filter was positive in the region around the characteristic carrier frequency of pulse song and negative elsewhere and normalized it between −100 and +100. We derived a standard weighted pulse spectrum by multiplying the pulse filter with the normalized power spectrum of an example trace. For the recording of an entire assay, we filtered it by a bandpass filter of 100Hz-1000Hz and then segmented the whole recording into 100ms bouts with 80ms overlap. We applied a short-time Fourier transformation to each bout, normalized the resultant power between −1 and 1, and multiplied it by the pulse filter to get a weighted spectrum for each bout. We defined a pulse score for each bout as the minimum Euclidean distance between the weighted spectrum and the standard weighted pulse spectrum by using dynamic time warping, allowing us to account for the shape of the power spectra. The pulse score for every 100ms was defined by the average pulse scores of bouts including it, and we subsequently binarily classified pulses based on scores exceeding the mean score by 0.75st.d. To assess the fraction of sound bouts that consisted of uninterrupted pulse trains, we identified envelopes longer than 200ms in which the average peak sound signal exceeded an energy threshold (0.01). We classified sine songs in a similar manner. We validated the classifier performance against manually labeled sound traces from 5 MF assays (**Figure S5G**).

#### Soundscape before copulation

We detected classified pulse probability by using the frequency-based method in the last 60 seconds before copulation and observed a sharp increase in the last 2-3 seconds before copulation (**Figure 5A**). To estimate the soundscape leading up to copulation events, we computed the power spectra of the final two seconds before manually scored copulation events. To account for the fact that the moment of coitus is difficult to precisely estimate, we also included 500ms following the onset of copulation in these soundscape calculations. To control for the possibility that shorter segments could be biased towards pulse song, we also randomly selected 100 peaks in regions with sound energy above 0.015 from each assay and computed the soundscape in the 2.5-second window around these peaks (**Figure 5B**).

#### Frequency probability distribution

We first detected sound bouts in MF and MMF recordings by setting the sound energy threshold and bout length threshold (> 200ms). We got the power threshold by z-scoring powers in all frequency domains in MF sound bouts. The Hindmarsh Sten and Li *et al*., 2023 23 frequency domains used in each sound bout were determined by whether the power is above this threshold. Then we computed the probability of each frequency domain for all MF and MMF recordings (**Figure S5C**).

### Immunohistochemistry and confocal imaging

Immunohistochemistry and confocal imaging were performed by Janelia’s core facility as previously described (Aso 2014, Schretter 2021). All confocal stacks were collected under 20X objectives. Each GAL4 line was crossed to the same UAS-CsChrimson:mVenus effector used for behavioral assays. A full step-by-step protocol can be found at: https://www.janelia.org/project-team/flylight/protocols.

To better visualize the morphology of R85A12 and pC1SS2 in both sexes (**Figures 3A, 6E, S4**, and **S9**), we used VVD viewer (Janelia) to skeletonize its arbors in the brain and VNC. We only included regions where we could confidently trace the neurites – thus, it is possible that the morphology of these neurons is slightly more extensive.

### Fly tethering and dissection

Virgin male flies were collected during 2-8h following eclosions and single-housed for 42-48h to increase their motivation to court^40,42,63,104,105^. Experiments were performed at Zeitgeber 0-3h. Virgin females were collected 2-8h following eclosion and group-housed at low densities (3-8 flies) for 3-6 days to ensure that they are mature but not too old.

For tethered behavioral experiments and two-photon functional imaging, flies were briefly anesthetized on CO_2_ and tethered to a custom-milled plate similar to those used in previous studies^40,106^. Flies were held in place by a string across the neck and fixed to the holder by both eyes and the back of the thorax using UV-curable glue (Loctite 3106). To minimize brain motion during functional imaging, the proboscis was also glued to the mouthparts. The string was subsequently removed, and flies were left to recover in a warm, humidified chamber (25°C, 50-70% humidity) in the dark. For behavioral experiments, flies were transferred to the ball after 2-6 hours. For functional imaging experiments, flies were left in the dark until immediately before the experiment, at which point the cuticle was removed to give optical access to the central brain without anesthesia. To do so, the tethering plate was filled with saline (108mM NaCl, 5mM KCl, 2mM CaCl_2_, 8.2 mM MgCl_2_, 4 mM NaHCO_3_, 1 mM NaH_2_PO_4_, 5 mM trehalose, 10 mM sucrose, 5 mM HEPES, pH 7.5 with osmolarity adjusted to 265 mOsm) to cover the fly’s head, and the cuticle between the eyes was cut with a needle (BD PrecisionGlide Hypodermic Needles 18G x 1” BD 305195) and removed with forceps. The trachea covering the top of the central brain was removed from both hemispheres with forceps. Flies were subsequently transferred to the ball and left to recover in darkness for at least 30 minutes.

### Virtual reality preparation

Virtual reality courtship experiments were performed as previously described (see Hindmarsh Sten et al., 2021 for details), with the addition of sound pulses for inducing agonistic behavioral patterns, and briefly summarized below.

#### Hardware

For our virtual courtship preparation, we adapted an existing hardware design for presenting tethered flies with visual stimuli (Jazz L. Weisman and Gaby Maimon, in preparation; Hindmarsh Sten *et al*., 2021). Male flies rested and walked on a small 6.35mm diameter ball, which was shaped from foam (General Plastics Last-A-Foam FR-4618) and manually painted with uneven black spots using a Sharpie. The foam ball was held by a custom-milled aluminum base with a concave hemisphere of 6.75mm. A 1mm tract drilled through the base was connected to air supplied at ∼0.8 L/min such that the ball could move smoothly. The aluminum base was held in place by a 3D-printed (printer: Carbon 3D, material: UMA-90) contraption. The ball was illuminated by infrared LED flood lights, and imaged with a Point Grey FLIR Firefly camera (FMVU-03MTM-CS) with a 94mm/1x WD Video Lens (InfiniStix) by way of a mirror (ThorLabs ME05-G01). The ball was surrounded by a 270° conical screen with a large diameter of ∼220 mm, a small diameter of ∼40mm, and a height of ∼60mm. The screen was laser cut from matte white 80lb cardstock (Desktop Publishing Services, Part #59421-50) and fitted into a custom 3D-printed screen holder (printer: Carbon 3D, material: UMA-90) with a tilted slit for placing and forming the screen shape. One computer controlled all sensory stimuli and stored fly positions through MATLAB scripts.

#### Parameters readout

As males walked on the foam ball, all three rotational axes of the ball were read out by the FicTrac2.0 software^107^ at 60Hz in real-time. FicTrac was linked by a socket to MATLAB, which read out the estimated angular position of the ball at each frame of an experiment. The animal’s updated position was read out in MATLAB and stored in the output matrix.

Videos collected by the camera which read ball positions were analyzed for wing extensions and wing flicks posthoc. Videos were time-stamped to the tracking data for behavior-stimulus alignment.

#### Visual stimuli

The visual stimulus was projected around the male from a DLP 3010 Light Control Evaluation Module (Texas Instruments). The light was projected by way of a 4×4×0.25’’ mirror (First Surface Mirrors) from below the fly. The red and green LEDs in the projector were turned off, leaving only the blue LEDs. The lens of the projector was also covered by a piece of blue filter paper (Rosco #071 Tokyo Blue). Together, this minimized the light from the projector collected by the microscope.

The projector received input from the same computer running FicTrac via an HDMI cable and was controlled via a mini-USB cable to the same computer. Visual stimuli were generated in the MATLAB-based ViRMEn software^108^ and projected onto the screen using custom perspective transformation functions. The net visual refresh rate of the visual stimulus ranged from 47.6Hz to 58.9Hz.

Each trial was initiated by the presentation of a stationary visual target for 60 sec to examine the animal’s baseline locomotion, after which the visual target began to oscillate. The visual target oscillated in a 107° arc around the animal with a constant angular velocity of ∼75°/s, but the angular size of the dot was continuously altered to mimic the dynamics of a natural female during courtship. The angular size was altered by changing the distance between the male and the target in the ViRMEn world. The distance between the male and the target was taken from the inter-fly-distance (IFD) in a courting pair over the course of two minutes of courtship, and at each frame, the angular position of the target was scaled by this IFD to give rise to a more dynamic female path. Angular sizes ranged between ∼8-50°, with the average size being 22.5°. Each stimulus frame was thus unique for 2 minutes of time, and subsequently repeated until the end of the trial when it intersected its original position. Each trial lasted 10 minutes.

#### Acoustic stimuli

To induce males to transition between courtship and agonistic wing flicks, we placed a speaker (ScanSpeak Discovery 10F/4424G, 4” Midrange 4 Ohm) at a 45° angle to the males, approximately 10cm from the ball, and delivery trains of *D. melanogaster* pulse song. The pulse song was generated in MATLAB with the same parameters as in the *sound playback free behavioral experiments* section, namely 35ms IPI and 200Hz carrier frequency. In each 10-minute visual stimulus trial, the pulse song was on in minutes 2-3. The sound was amplified through an amplifier (Crown XLS1502 2-Channel 525W Power Amplifier). The sound volume for each stimulus was consistent across trials.

### Tethered behavioral analysis

#### Fidelity, Vigor, and Tracking Index

To estimate how well animals were actively tracking the stimulus, we computed both the vigor and fidelity of their pursuit in a sliding time-bin of 180 frames (∼3.7 sec). We defined the fidelity of a male’s pursuit as the correlation between the position of the visual target and the change in a male’s heading (rad/s), and the vigor as the net amount of turning the male exhibited in the direction ipsilateral to the visual target. Because neither of these metrics fully captured male behaviors, we defined a Tracking Index (T.I.) as the product of the fidelity and the within-animal normalized vigor of pursuit (vigor in the current cycle divided by the maximum vigor observed). This normalization step was done to bound the T.I. between −1 and 1 and to correct for any difference in males’ ability to turn on the ball. The TI was set to zero in the first and last 90 frames, as we did not have a sufficient number of frames to compute it.

#### Male wing motor programs classification

To detect bouts of unilateral wing extensions and bilateral wing flicks we manually drew two ROIs adjacent to the wings, and integrated the pixel intensity in these regions over time. During wing extensions and wing-flicks, the lightly IR-reflective wings moved to briefly cover the dark background in these regions, resulting in a transient increase in pixel identity. If both ROIs synchronously exhibited increased intensity (2st.d. > mean intensity) we flagged a putative bilateral wing flick, while we flagged a putative unilateral wing extension when the intensity of one ROI alone was increased. All wing extensions and bilateral wing flicks were subsequently manually validated by the experimenters.

#### Courtship and agonistic behaviors classification

We only used males whose average linear speeds exceeded 5mm/s, or whose angular speeds exceeded 2rad/s to exclude males with locomotion defects. We defined non-courting males as those whose tracking index (T.I.) never exceeded 0.3 within the described periods (Figs). We defined non-aggressive males as those who never displayed any wing flicks or exhibited a pulsatile increase in their linear speed during the song playback.

The threshold value for the T.I. was selected so as to be above fluctuations in the T.I. during random running^40^. The threshold for angular and linear speed was selected to be well above the noise of a fly standing still caused by small vibrations in the floating ball^40^.

#### Heat maps of turning, locomotion, and behaviors

Turning was computed on a frame-by-frame basis as the circular distance between the animal’s current heading and the animal’s heading in the next frame using the MATLAB circular statistics toolbox (v. 1.21.0.0)^109^. A male’s linear speeds were computed based on the changes in a male’s angular positions read out from FicTrac. The classification of wing motor programs was as described above.

Heat maps were constructed by computing the phase length (in frames, PL) of the stimulus and multiplying it by 3 (3PL). All frames were fit into a matrix of size Nx-3PL. A very small number of remnant frames (typically < 0.5% of frames) at the end of the trial, caused by the total frames not being divisible by 3PLN, were discarded from heat maps but included in all other analyses.

### Two-photon functional imaging

Because of the low expression level of the driver lines we were using, we chose bright calcium sensors (2xjGCaMP7f and jGCaMP7s) for functional imaging. Functional imaging experiments were performed with an Ultima Investigator or Ultima Investigator Plus two-photon laser scanning microscope (Bruker Nanosystems) with a Chameleon Ultra II Ti:Sapphire laser. All samples were excited at a wavelength of 920nm, and emitted fluorescence was detected with a GaAsP photodiode detector (Hamamatsu). All images were acquired with a 40X Olympus water-immersion objective with 0.8 NA. All images were collected using PrairieView Software (Version 5.5 or 5.7) at 1024 pixel × 1024 pixel resolution for females or 512x512 pixel for males. To reduce noise caused by emitted light from the projector during imaging, we turned off red and green LEDs in the projector, placed a piece of blue-light filter paper (Rosco #071 Tokyo Blue) in front of the projector lens, and 3D printed a custom light shield (printer: Carbon 3D, material: UMA-90) that fit over the objective to prevent light from entering the brain from above.

After cuticular removal as described above, flies were carefully lowered onto the ball using a micromanipulator (Scientifica), and the shrouded objective was lowered over the brain. We subsequently identified the brain region of interest and centered a small ROI over it, yielding an imaging rate of 10-14Hz for females and 7-12Hz for males. Flies were left in the dark to acclimate after ROI selection. Power was kept low to minimize heating fly brains and we ensured that no pixels were saturated. On rare occasions, flies were discarded because the glue holding the proboscis broke loose from the mouthparts, causing severe motion artifacts in the z-direction, or because the expression of GCaMP was too weak to detect the neuropil of interest at low imaging power.

### Imaging ROI selection

The hemisphere targeted for imaging was altered between experiments.

#### vpoEN

Given how deep major vpoEN neurites resided in the dorsal-ventral axis, we selected an ROI that was relatively close to the dorsal brain surface and was at the lateral side where many neurites converged^14^.

#### vpoIN

Given how deep major vpoIN neurites resided in the dorsal-ventral axis, we selected an ROI that was relatively close to the dorsal brain surface and was at the lateral side where many neurites converged^14^.

#### P1a

An ROI was selected to cover the lateral junction in the medio-lateral and dorsal-ventral axes, below the P1a arch, and above the protrusion of the P1a ring^40,110^.

#### pC1x

An intricate ROI was selected to cover the sparse neurites of pC1x neurons near the lateral junction in the medio-lateral and dorsal-ventral axes, near the axon terminals of P1 neurons^110^. The imaging plane is illustrated in **Figure S9C**.

##### Acoustic stimuli

Acoustic stimuli to tethered males during imaging were described in the *virtual reality preparation* section.

For delivering acoustic stimuli to females during imaging, we adapted the sound delivery system described in Tootoonian, et al. 2012^111^. Sound from the computer was amplified through an amplifier (Samson SAQH4), which was connected to a pair of headphones (Koss ‘The Plug’). Each headphone was in turn connected to an ∼80mm long tubing (I.D. 0.125”, O.D. 0.25”, Thermo Scientific 80100125). Tubings on each side were directed towards the left and right aristae of the female respectively from the back, with the end of each tubing approximately 5mm away from the aristae. The sound volume for each stimulus was consistent across trials.

For testing responses of vpoEN and vpoIN neurons to unadulterated MF courtship song and agonistic song (**Figure 4B**), a short segment (∼1s) of the courtship song was selected from MF recordings, and a short segment (∼1s) of the agonistic song was selected from R85A12 activation recordings. Given the amplitudes of agonistic song recorded from optogenetic activation are not natural, we ensured that the sound volumes of maximum amplitude in the courtship and agonistic song were the same. A one-second synthetic *D. melanogaster* pulse train was delivered at the beginning and the end of each imaging session as positive controls to exclude the possibilities of auditory defects and sensory adaptation over time. Courtship and agonistic song stimuli were both delivered three times and the order of stimuli is randomized. The interval between neighboring acoustic stimuli is 15 seconds. We left 10 seconds before the first stimulus in each session for baseline fluorescence computation.

For synthetic pulse songs with offsets (**Figure4E**), we synthesized *D. melanogaster* pulse songs with parameters as described in the *sound playback free behavioral assays* section. Each pulse train lasted 1s, and the pulse trains in the two channels were offset. We chose the channel with the leading stimulus based on the imaging hemisphere. Given that the pulse trains followed a cycle of 35ms, we designed the offsets to range from 0 to 35ms to cover different possibilities of overlapping pulse songs (from 0 to 2π). We also included stimuli with synthetic pulse song only from one channel to assess the ability to distinguish sound from different locations on the neuronal level. A one-second synthetic *D. melanogaster* pulse train was delivered at the beginning and the end of each imaging session as positive controls to exclude the possibilities of auditory defects and sensory adaptation over time. The order of all stimuli is randomized and the interval between neighboring acoustic stimuli is 20 seconds. The stimuli panel was repeated 1-4 times on each fly. We left 10 seconds before the first stimulus in each session for baseline fluorescence computation.

For natural songs (**Figures 4C, S6A**, and **S6B**), we selected three MF and MMF recordings that exhibited copulation latencies of about three minutes, and assigned them to one of three stimulus groups. Each fly received at least one group of MF and MMF recordings for paired analysis. During analysis, we only used the regions of acoustic stimuli from the start of all males engaging in courtship till copulation. The amplitude differences between MF and MMF recordings were preserved during playback. We left 10 seconds before the start of the acoustic stimulus in each session for baseline fluorescence computation.

### Imaging analysis

Image stacks were motion corrected using Non-Rigid Motion Correction (NoRMCorre^112^) and were subsequently manually validated for motion artifacts. For each experimental recording, an ROI was drawn in FIJI (ImageJ, NIH) across the entire population of interest containing neuropil, and the average fluorescence for each imaging frame was extracted. Fluorescence was normalized in MATLAB by assuming that the pre-acous-tic-stimulus (for females) or pre-visual-stimulus (for males) represented the baseline fluorescence of the populations of interest. The average fluorescence of the first 20 frames (∼2s, for females) and 100 frames (∼10s, for males) of recording was used as the baseline (F_0_), and an ∆F/F_0_ was defined as: Fi-F_0_/F_0_, where i denotes the current frame.

All female imaging data were downsampled to 10Hz by using an FIR Antialiasing Lowpass Filter and compensating for the delay introduced by the filter. To visualize relations between excitatory and inhibitory auditory inputs to vpoDN (**Figure 4D**), we normalized vpoEN and vpoIN activities (∆F/F_0_) by the maximum activities of vpoEN and vpoIN observed in all natural songs (MF and MMF recordings) playback trials.

For male imaging-behavior correlations (**Figure S10**), because imaging data were collected at a lower frame rate than behavioral data, we downsampled the behavioral data using linear interpolation at the imaging time points to allow us to compute correlations between behavior and imaging.

### Data Reporting

Preliminary experiments were used to assess variance and optimize behavioral conditions. Experiments were not randomized, and the experimenters were not blind to the conditions. For tethered courtship assays, unless otherwise noted, only experiments during which animals exhibited courtship towards the visual targets were included for analysis, selected based on a tracking index > 0.3 for at least 1 second and the presence of at least one unilateral wing extension. For analysis of functional responses to sound playback, unless otherwise noted, only animals that flicked in response to the stimulus were included. For tracking index computation and wing flicks classification, see *tethered behavioral analysis*. Some free behavioral assays with long copulation latencies were excluded for consistency (see *free behavioral assays*).

### Statistics and reproducibility

All statistical analyses were performed in MATLAB (2019b, 2021b-2023a) or GraphPad Prism 9. Data sets were tested for normality using Shapiro-Wilk, and appropriate statistical tests were applied (e.g. t-test for normally distributed data, Mann Whitney U-test for non-parametric data, Log-Rank test for survival distributions of two samples). All statistical tests used were two-tailed. Details of statistical analyses and sample sizes are given in Supplementary Table 2. Experimenters were not blind to the conditions of the experiments during data collection and analysis.

## KEY RESOURCES TABLE

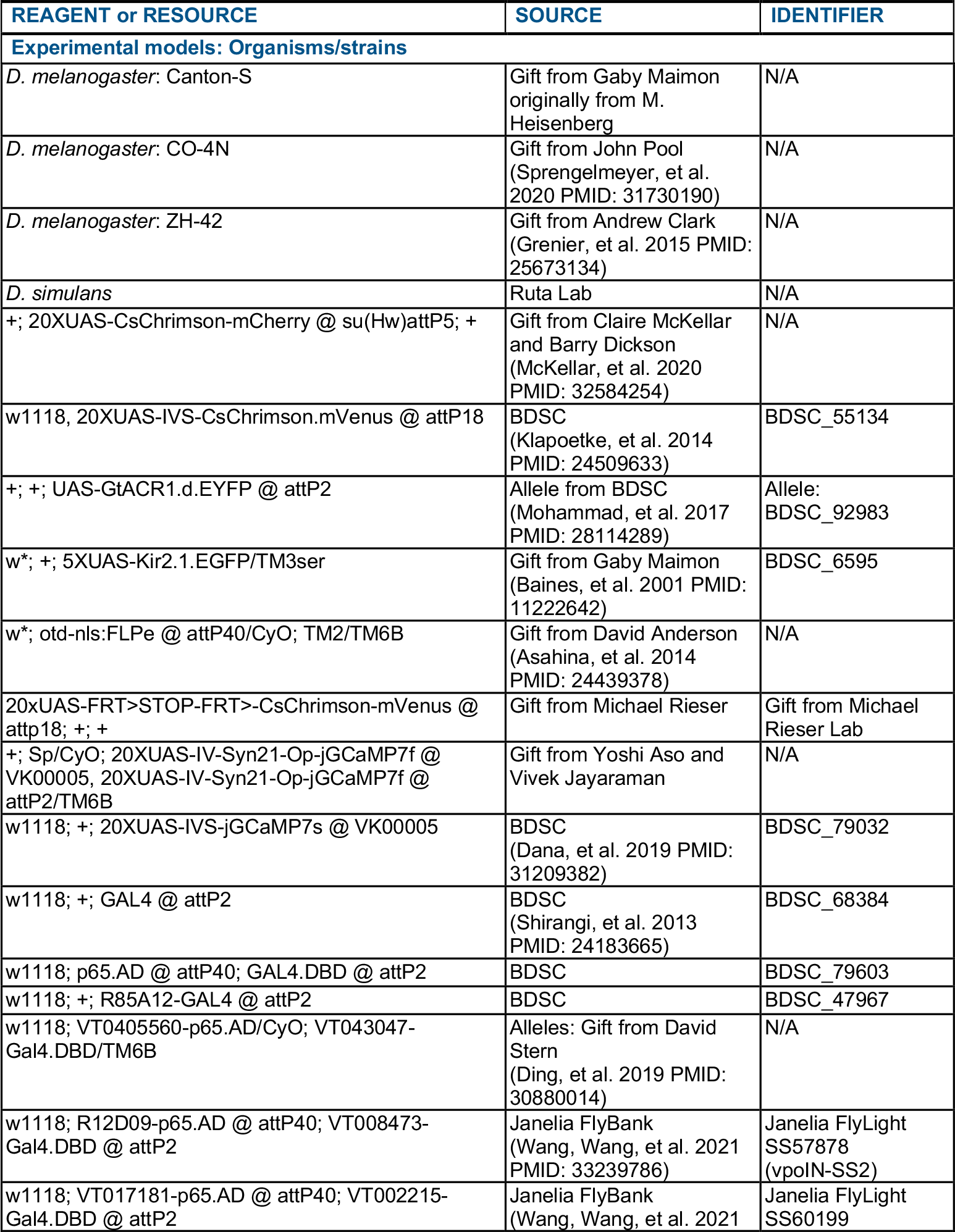
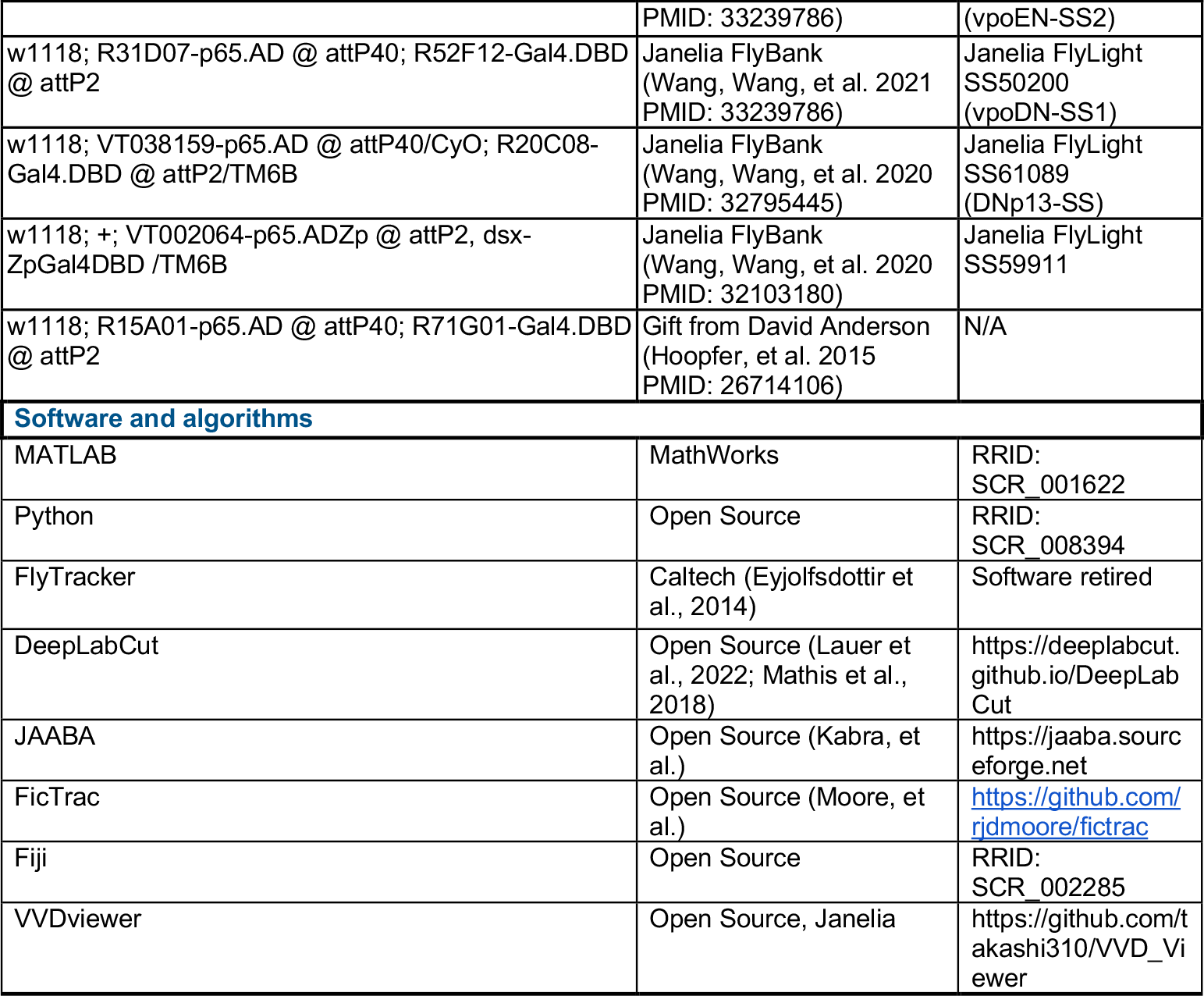

**Figure S1.**
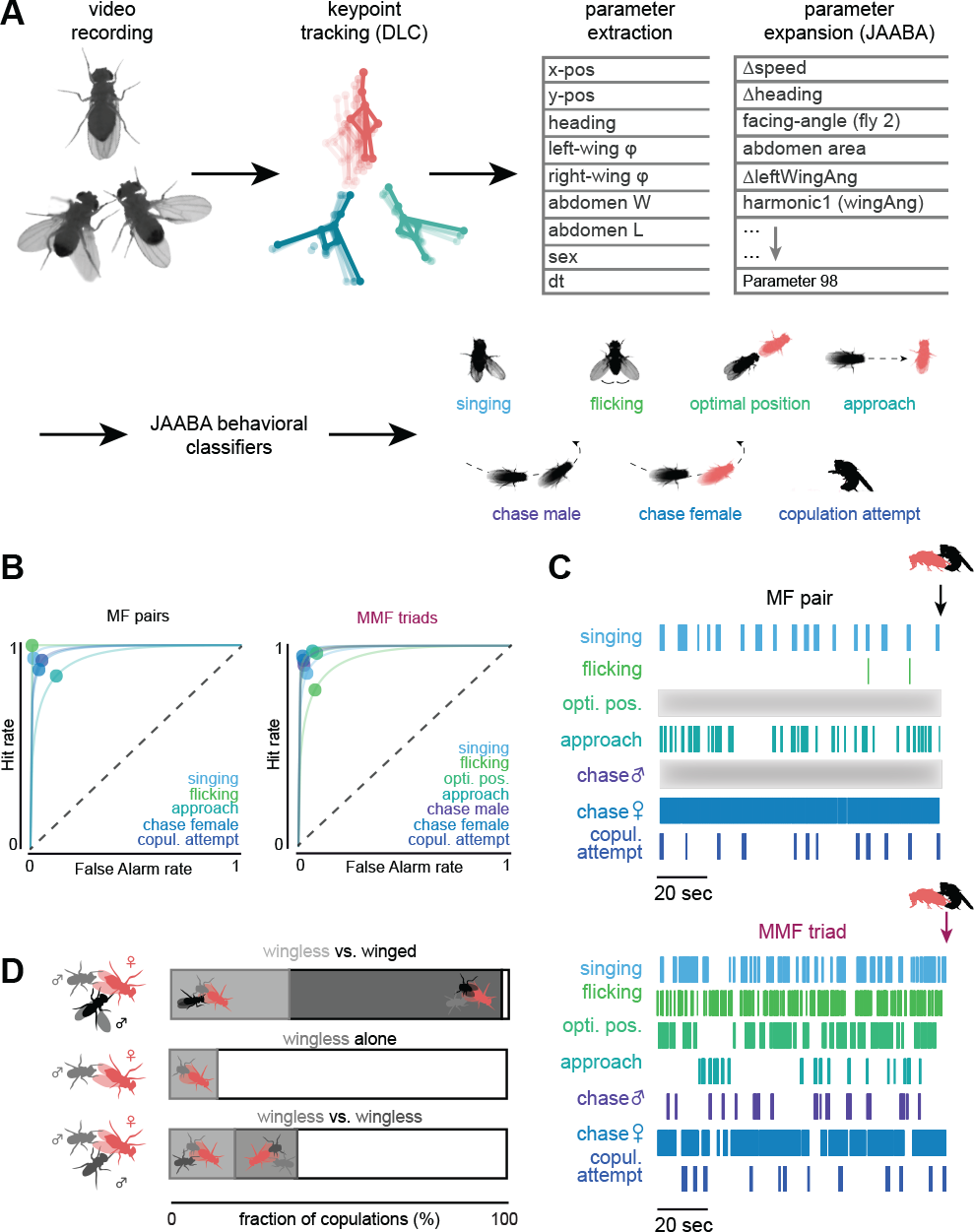
Classification of behavioral dynamics in competitive courtship, related to Figure 2. (A)Analysis pipeline for behavioral quantification. For each assay, we performed supervised tracking using eight key points for each animal (head, thorax, top of the abdomen, left and right sides of the abdomen, genitalia, and the left/right wing-tips) using DeepLabCut, and quantified coarse postural metrics. Postural metrics were transformed and expanded to ∼100 behavior metrics in JAABA, including derivatives of postural parameters across time, yielding N(number of parameters) X T (number of frames) matrices. Classifiers for core component behaviors of competitive courtship were built by JAABA using these metrics from example videos in a supervised manner. We used these classifiers to classify epochs of behaviors in MF pairs and MMF triads. (**B**) Classifier sensitivity and specificity in MF pairs (left) and MMF triads (right). Dots represent the average classification accuracy across behaviors, lines are inferred isosensitivity (d’) curves. (**C**) Ethograms of male behaviors in a characteristic MF pair (left) and MMF triads (right) across one minute prior to copulation. (**D**) Comparison of copulation rates with wingless males for females offered one intact and one mute wingless male (top, 35.2%, n = 68; same data as shown in Figure 1G); a single mute wingless male (middle, 14.7%, n = 38) or two wingless males (bottom, 18.3%, n = 41). Note that the fraction of females copulating with wingless male is higher in the presence of a winged male.

**Figure S2.**
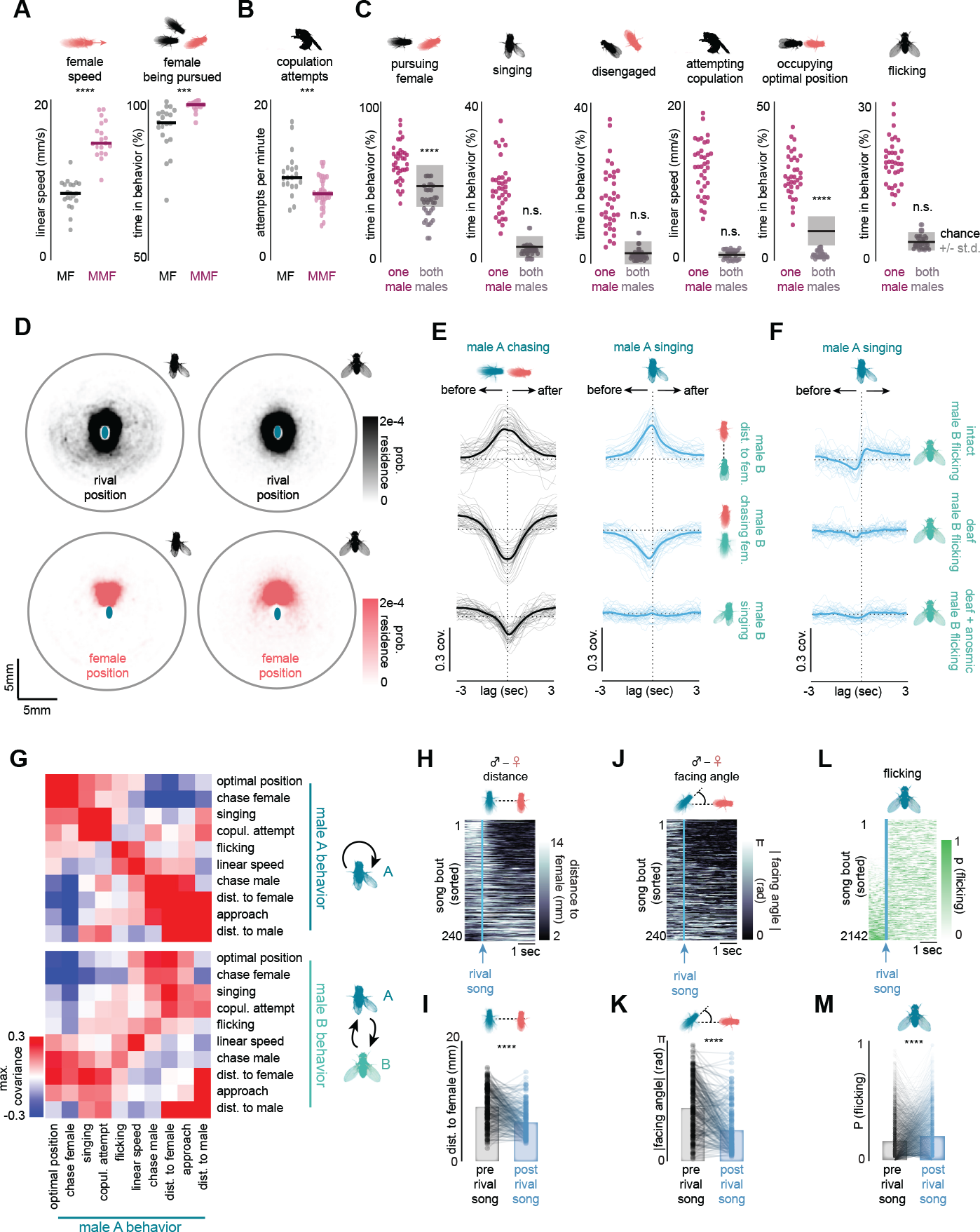
Spatiotemporal relationships between male component behaviors in MMF triads, related to Figure 2. (A)Average female speed (left) and the fraction of time females were pursued by a male (right) in MF (black) MMF assays (purple). (**B**) Number of copulation attempts performed per minute by males in MF versus MMF assays. (**C**) Fractions of time when at least one male (purple) or both males simultaneously (gray, individual dots) performed indicated core component behaviors during MMF assays. Lines with shaded regions denote the null probability (mean +/-std) of both males performing the same behavior simultaneously by chance, based on the frequency of each behavioral component across the entire assay. Statistics denote significant deviations from the null hypothesis that male behaviors are independent. (**D**) Heat maps showing the position that the female (bottom row) or rival male (top row) occupies relative to a male while he is performing unilateral wing extensions (left) or bilateral wing flicks (right). (**E**) Cross-covariance between one male’s chasing of the female (left) or performance of unilateral wing extensions (right) and the other male’s distance to the female (top), the other male’s chasing of the female (middle), or the other male’s performance of unilateral wing extensions (bottom). Positive lags indicate that male A leads. Note that the anti-correlation between positional variables indicates that males alternate between occupying the position closest to the female, and preferentially sing when their rival is further from the female. However, the unilateral extension of one male does not appear to directly promote or suppress the unilateral wing extensions of the other male (see C). (**F**) Cross-covariance between one male’s unilateral wing extensions and the other male’s bilateral wing flicks for intact males (top, same as Figure 2J), deaf males (aristae removed), and males that were both deaf and anosmic (third antennal segment removed). Positive lags indicate that unilateral wing extension leads. (**G**) Maximum covariance within a 1-second lag between the behaviors performed by one male and either the behaviors performed by his rival (bottom) or the behaviors performed by the same male. (**H**) Heatmap of the distance between non-courting males and the female, aligned to when a rival performs a unilateral wing extension. (**I**) Average distance of non-courting males to the female 1 second before and 1 second after a rival performed a unilateral wing extension. Dots denote individual wing extensions from the rival. (**J** and **K**) Same as H and I, but showing the absolute facing angle of the non-courting male relative to the female’s position in a triad. (**L**) Heatmap of the probability that a male is performing a bilateral wing flick, aligned to the onset of a rival’s unilateral wing extension. Note that males that were not flicking prior to his rival singing began to do so. (**M**) Probability of a male performing bilateral wing flicks across the second prior to versus after a rival performed a unilateral wing extension. Dots denote individual wing extensions from the rival. Thin lines denote individual animals; dot plots with line denote individual males and average; dot plots with bars denote individual behavioral segments; n.s., P > 0.05; ***P < 0.001; ****P < 0.0001. Details of statistical analyses and sample sizes are given in **TableS2**.

**Figure S3.**
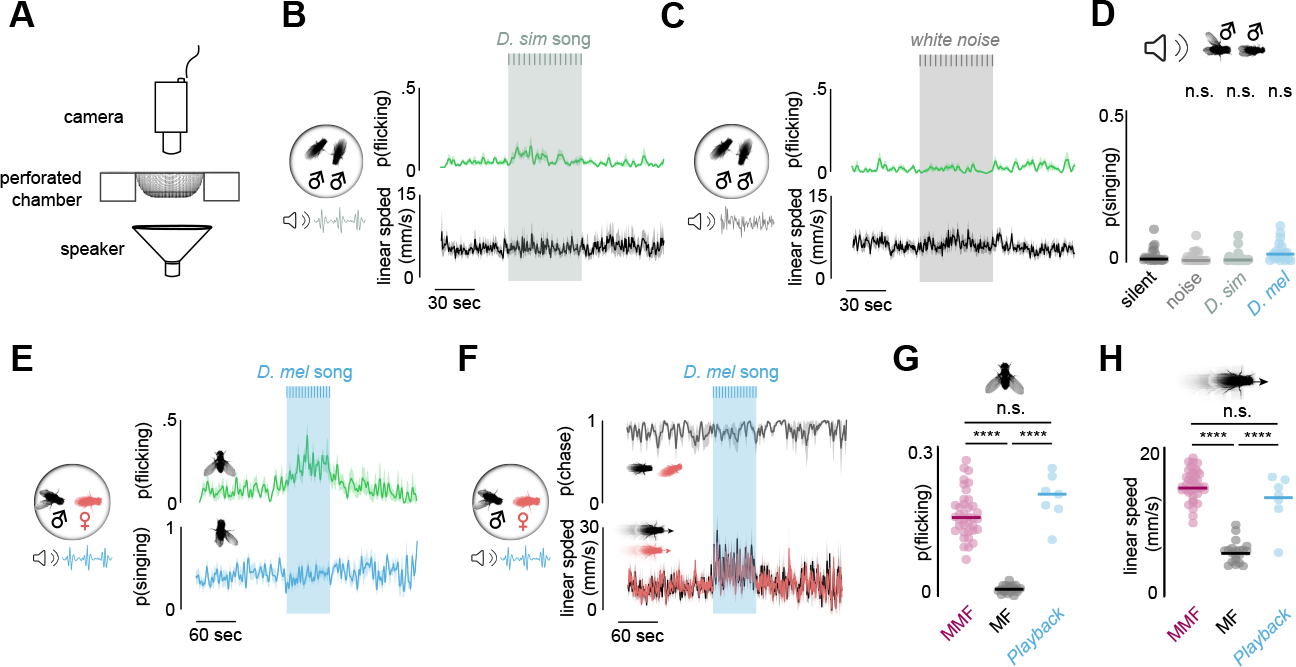
Quantification for behavioral assays with sound playback, related to Figure 2. (**A**) Schematic of the behavioral setup used for song playback of freely moving animals. (**B** and **C**) Probability of males exhibiting bilateral wing flicks (top) and male linear speed (bottom) prior to, during, and after 1 minute of synthetic *D. simulans* song playback (B) or white noise (C). Shaded region denotes playback period, darker gray hatches denote individual trains of pulse song. (**D**) Probability of males exhibiting unilateral wing extensions when paired with another male. during playback of no sound (black), white noise (gray), synthetic *D. simulans* song (light gray), and synthetic *D. melanogaster* song (blue). (**E**) Probability of males exhibiting unilateral wing extensions (top) and chasing the female (bottom) prior to, during, and after 1 minute of synthetic *D. simulans* song playback to courting MF pairs. (**F**) Probability of males chasing (top) and linear speed of males and females (bottom) prior to, during, and after 1 minute of synthetic *D. melanogaster* song playback to courting MF pairs. Note that playback induces agonistic actions but does not suppress courtship pursuit (same assays as shown in E). (**G** and **H**) Probability of males performing bilateral wing flicks (G) and the average linear speed (H) of males in MMF triads, MF pairs, and MF pairs with D. melanogaster pulse song playback. Shaded lines show mean ± s.e.m.; Lines with dots represent mean and individual animals. n.s., P > 0.05; ****P < 0.0001. Details of statistical analyses and sample sizes are given in **TableS2**.

**Figure S4.**
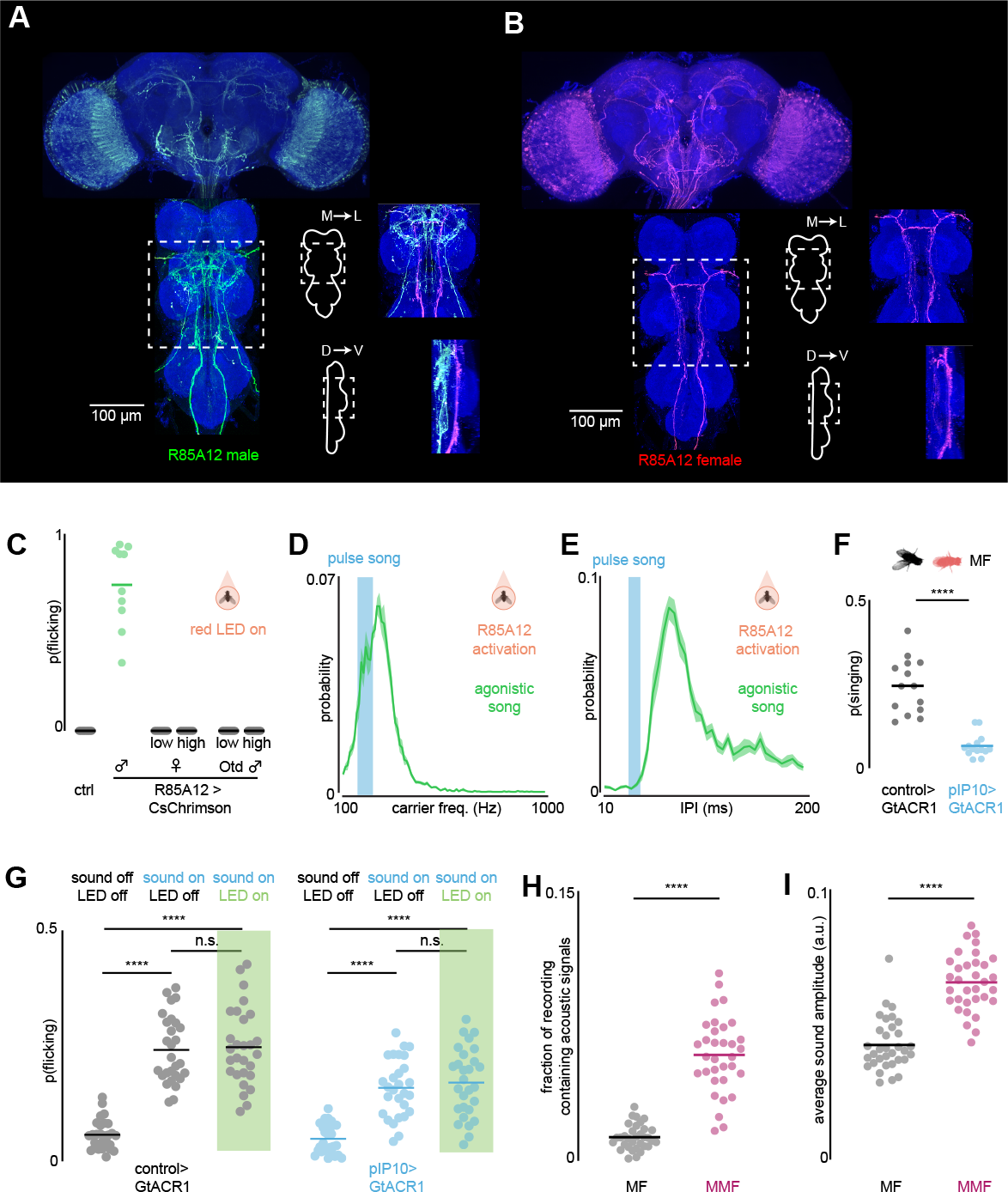
Sexually dimorphic neurons in the male ventral nerve cord (VNC) control agonistic song, related to Figure 3. (**A** and **B**) Immunostaining of neurons labeled by R85A12-Gal4 in males (A) and females (B). Insets show maximum fluorescence projections in the VNC region outlined by the white box along the medial-lateral axis (top) or dorsal-ventral axis (bottom); sexually monomorphic neurites found in both males and females are shown in magenta, and male-specific neurites in green. Note that while male-specific neurites project along the dorsal side of the VNC, sexually monomorphic neurites project along the ventral side, rendering them readily anatomically separable. (**C**) Probability that animals performed bilateral wing flicks during red light stimulation in control males, males in which R85A12 drives CsChrimson expression, females in which R85A12 drives expressing CsChrimson expression, or males in which R85A12 drives CsChrimson expression but intersected by Orthodenticle (Otd), restricting expression to the brain. “Low” reflects the standard light intensity. Given the absence of bilateral wing flicks in females and Otd-intersected males, experiments were repeated at higher light intensities (see Methods). Note that males did not perform bilateral wing flicks when the expression was restricted to the brain, indicating that the causal neurons reside locally within the VNC. (**D** and **E**) Distribution of carrier frequencies (E) and inter-pulse-intervals (F) of the agonistic song elicited by optogenetic activation of neurons labeled by R85A12 in males. Blue line shows the canonical carrier frequency and inter-pulse-interval of D. melanogaster pulse song. (**F**) Fraction of time control males or males in which the pIP10 descending neurons were optogenetically silenced via expression of GtACR1 spent performing unilateral wing extensions during MF assays. (**G**) Fraction of time control males or males in which the pIP10 descending neurons express GtACR1 spent performing bilateral wing flicks when two males were paired in a perforated chamber with no sound, with playback of pulse song but no optogenetic silencing, or with playback of pulse song along with optogenetic silencing (green bar). Note that pIP10 silencing diminished unilateral wing extensions but not the bilateral wing flicks induced by song playback. (**H**) Fraction of acoustic recordings from MF pairs versus MMF triads that were significantly above background noise levels. (**I**) Average signal amplitudes of acoustic recordings from MF pairs versus MMF triads. Shaded lines show mean ± s.e.m.; Lines with dots represent mean and individual animals. n.s., P > 0.05; ****P < 0.0001. Details of statistical analyses and sample sizes are given in **TableS2**.

**Figure S5.**
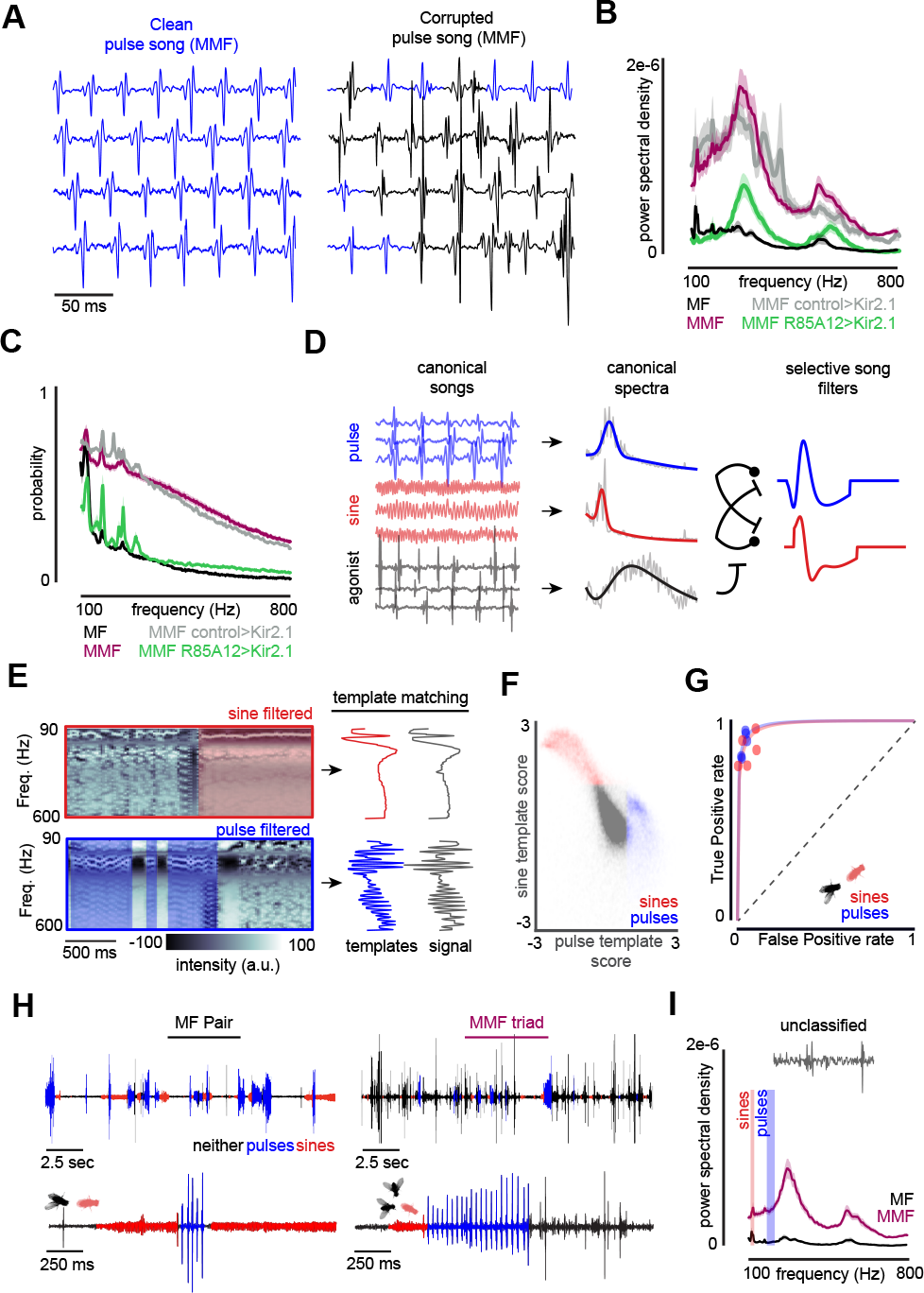
Acoustic signals are corrupted during competition, related to Figure 3. (**A**) Example traces of acoustic recordings from MMF triads showing bouts of unadulterated pulse song (left) and corrupted pulse song (right). Blue regions indicate epochs of pure unadulterated pulse song in both. (**B** and **C**) Average power spectra (B) and the distribution of frequencies (C; see Methods for details) in acoustic recordings from MF assays (black) and MMF assays with wild-type males (purple), R85A12>Kir2.1 males (green) or with >Kir2.1 control males. (**D** and **E**) Schematic depicting a frequency-based pulse classification method. Characteristic power spectra were generated from bouts of pure pulse and sine song recorded from MF assays, as well as bouts of pure agonistic song recorded during optogenetic activation of R85A12 neurons (D, left, middle). Selective filters for sine and pulse song were generated by subtracting resultant filters from one another (D, right; see methods for details). Normalized spectrograms from entire acoustic recordings were multiplied by each filter, allowing for robust visual detection of pulse and sine epochs (E, left; blue and red boxes denote periods of pulse and sine song, respectively). For each time point, we subsequently compared the similarity (minimum euclidean distance with dynamic time warping) of the resultant filtered spectra to canonical templates of sine and pulse song to create a pulse and sine score (E, right). (**F**) Z-scored pulse versus sine scores for each time point across a characteristic MF assay. Epochs classified as pulse (blue) or sine (red) song are color coded. Note that most epochs were classified as neither, reflecting the natural discontinuity of a male’s courtship song and the fact that males only sing a fraction of the courtship trial. (**G**) Sensitivity and specificity of frequency-based classifier for sine (red) and pulse (blue) song detection across 5 MF pairs. Dots represent the classification accuracy across individual assays; lines are inferred isosensitivity (d’) curves. (**H**) Example acoustic recordings from MF (left) and MMF (right) assays, with classified bouts of pulse (blue) and sine (red) song overlayed. Black epochs were classified as neither sine nor pulse. Bottom is a zoom-in across 2 seconds of the top trace. Note that in recordings from MMF assays, many epochs are not classified as either sine or pulse due to acoustic overlap from the two males. (**I**) Average power spectra of epochs that were classified as neither pulse nor sine song in acoustic recordings from MF (black) and MMF (purple) assays, demonstrating that the residual signal in MMF assays has a predominant carrier frequency of ∼300Hz, the same as agonistic song, while the residual signal in MF assays is minimal. Shaded lines show mean ± s.e.m. Details of sample sizes are given in **TableS2**.

**Figure S6.**
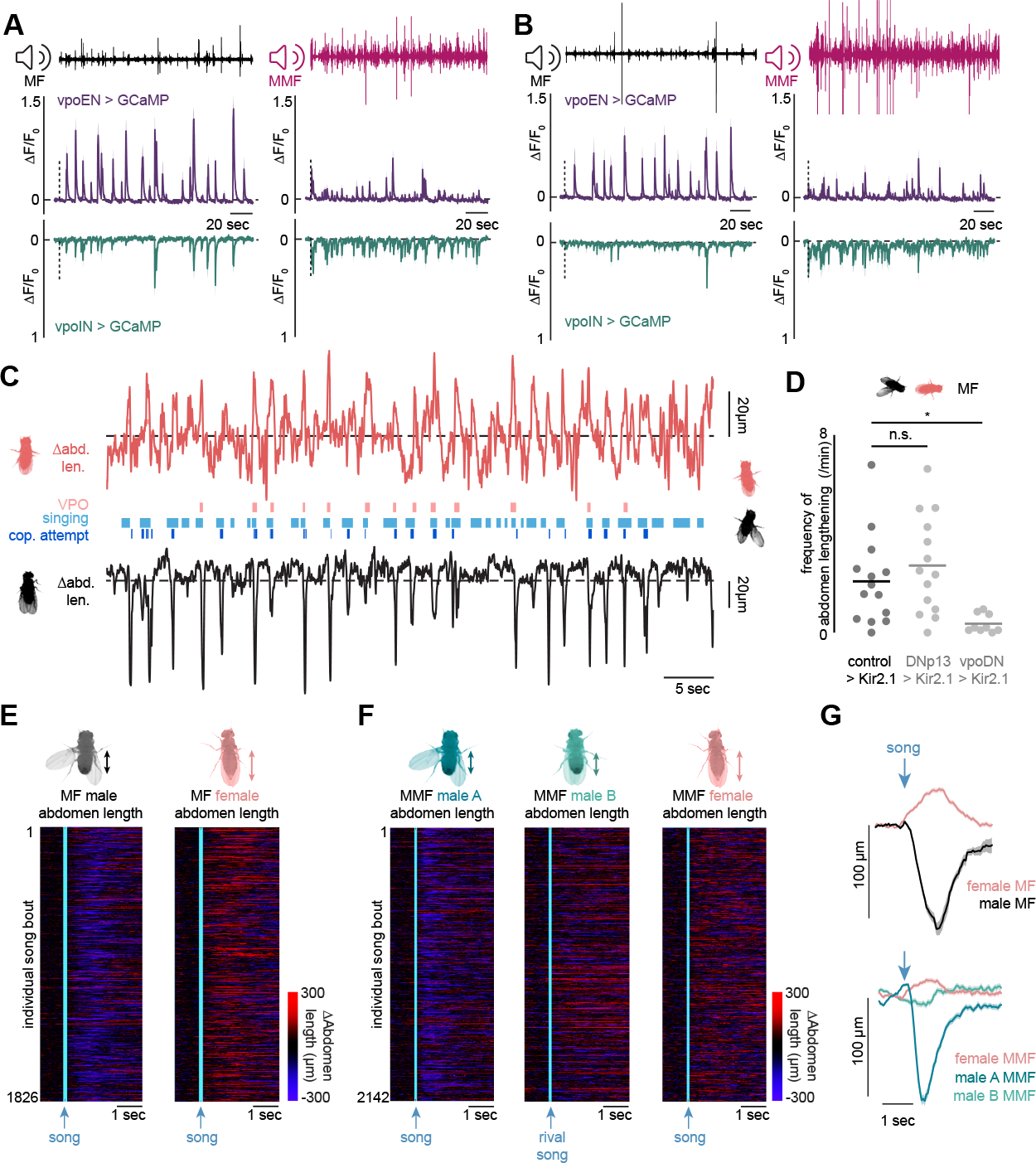
A complex acoustic environment disrupts the behavioral feedback loops between males and females during competitive courtship, related to Figure 4. (**A** and **B**) Average evoked vpoEN (purple) and vpoIN (green) activity (∆F/F_0_) during playback of two additional examples of acoustic recordings from MF (left) and MMF (right). Sound traces on the top show the recording being played back. Horizontal dashed line traces zero, vertical dashed line denotes the onset of sound playback. (**C**) Representative ethogram (2-minute long) of male and female component behaviors in MF pairs, plotted alongside the length of their abdominal segments. Females typically perform vaginal plate openings in response to male courtship songs, while males attempt copulation during vaginal plate openings. (**D**) Frequencies of female abdomen lengthening events in MF assays with control virgin females, or virgin females in which DNp13 or vpoDNs were constitutively silenced by expression of the inward-rectifying potassium channel Kir2.1. DNp13 and vpoDN are descending neurons essential for ovipositor extrusions and vaginal plate openings, respectively. While virgin females typically respond to male courtship song by performing vaginal plate openings, mated females instead reject males by performing ovipositor extrusions^12,69^. Note that silencing of vpoDN but not DNp13 neurons significantly diminished the frequency of abdominal lengthenings, indicating that these epochs predominantly reflect vaginal plate openings. (**E**) Heatmap of male (left) and female (right) abdomen lengths in MF assays, aligned to the onset of the male’s unilateral wing extension across all assays. The male abdomen contracts as he performs a copulation attempt, while the female abdomen lengthens during vaginal plate openings. (**F**) Same as (E), but for MMF assays aligned to when male A performs a unilateral wing extension. (**G**) Top: average female abdomen length (pink) and male abdomen length (black), aligned to the onset of the male’s unilateral wing extensions. Bottom: average female abdomen length (pink), male A abdomen length (blue), and male B abdomen length (teal) aligned to the onset of the male A’s unilateral wing extensions. Thin lines denote individual animals; shaded lines show mean ± s.e.m.; Lines with dots represent mean and individual animals. n.s., P > 0.05; **P < 0.01. Details of statistical analyses and sample sizes are given in **Table S2**.

**Figure S7.**
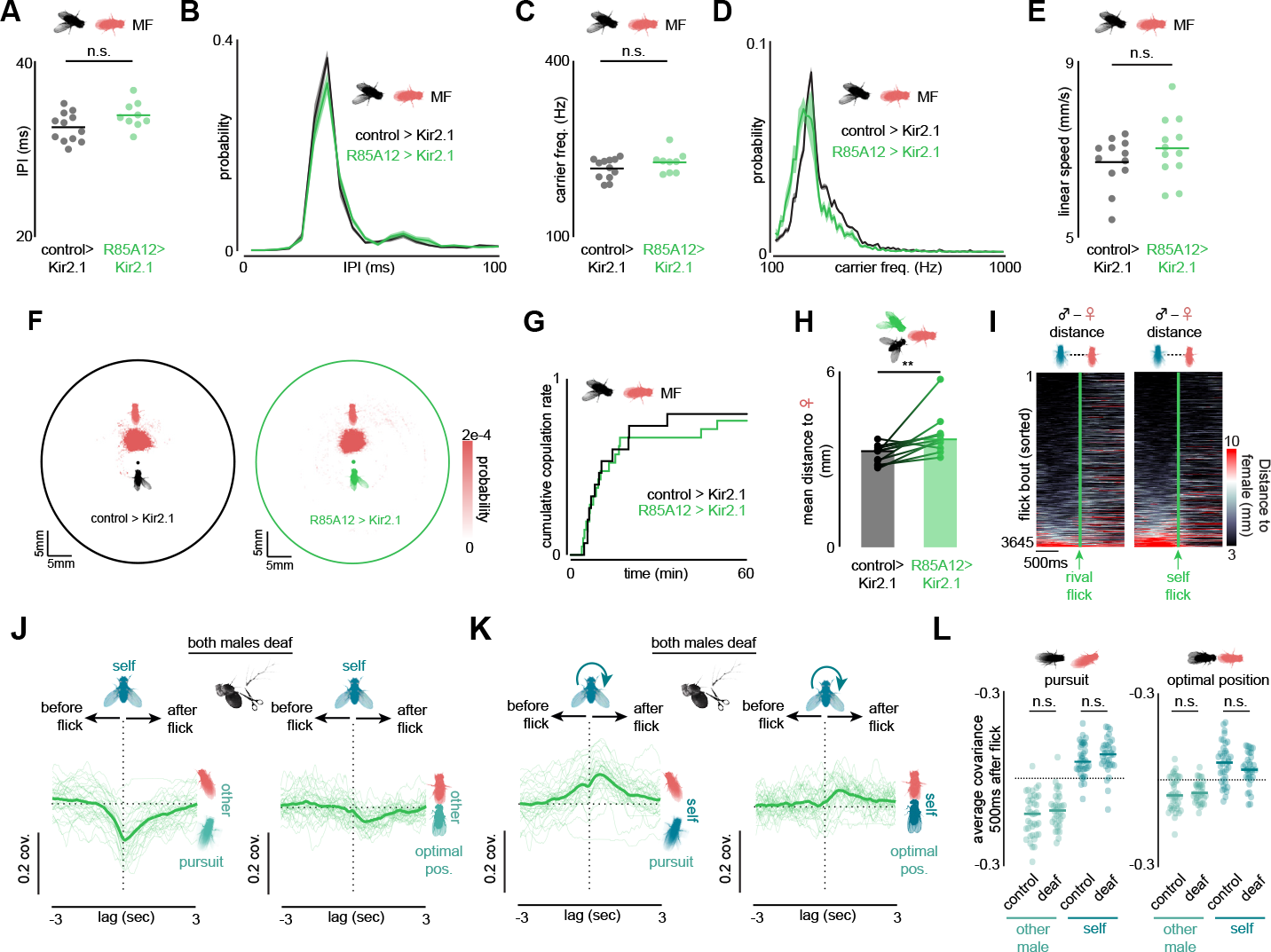
Agonistic wing flicks repel rival males, related to Figure 5. (**A**-**D**) Average values (A, C) and distributions (B, D) of the inter-pulse-interval (A, B) and carrier frequency (C, D) of courtship song recorded from MF assays with control males or males in which R85A12 neurons were constitutively silenced via expression of Kir2.1. (**E**) Average linear speed of control males or males whose R85A12 neurons were constitutively silenced via expression of Kir2.1 during MF assays. (**F**) Heat maps showing the position of the female relative to a male (central dot) during MF assays with control males (left) or males in which R85A12 neurons were constitutively silenced (right). Regardless of genotype, males efficiently center the female in their field of view and track her closely. (**G**) Cumulative fraction of copulations in one-hour MF assays, when a female was paired with a control male (black) or a male in which R85A12 neurons were constitutively silenced via expression of Kir2.1 (green). Despite their decreased mating success in competitive environments (see Figure 5F), R85A12-si-lended males effectively copulate with females in the absence of rivals. (**H**) Paired comparison of the average distance from the female to each of the two males in an MMF triad, in which one male is a control male and the other male has R85A12 neurons constitutively silenced. (**I**) Heatmap showing the average distance of a male to the female in wild-type MMF assays, aligned to when his rival performs a bilateral wing flick (left) or when the male himself performs a bilateral wing flick (right). (**J**) The cross-co-variance between one male’s bilateral wing flicks and the other male’s pursuit of the female (left) or the other male’s occupation of the optimal copulatory position behind the female (right) in MMF triads when both males were deafened by having their aristae removed. Positive lags indicate that bilateral wing flicks lead. (**K**) Same as (J) but showing the cross-covariance between one male’s bilateral wing flicks and his own pursuit (left) or position relative to the female. (**L**) Comparison of the average covariance in the 500 ms after a wing-flick and male pursuit (left) and occupancy of the optimal position (right) shown for both the flicking male (self) and his rival (other), in normal MMF triads, (data from Figure 5D and 5E), and deaf male MMF triads (data in J and K). Shaded lines show mean ± s.e.m.; lines with dots represent mean and individual animals; thin lines denote individual animals; n.s., P > 0.05; **P < 0.01. Details of statistical analyses and sample sizes are given in **TableS2**.

**Figure S8.**
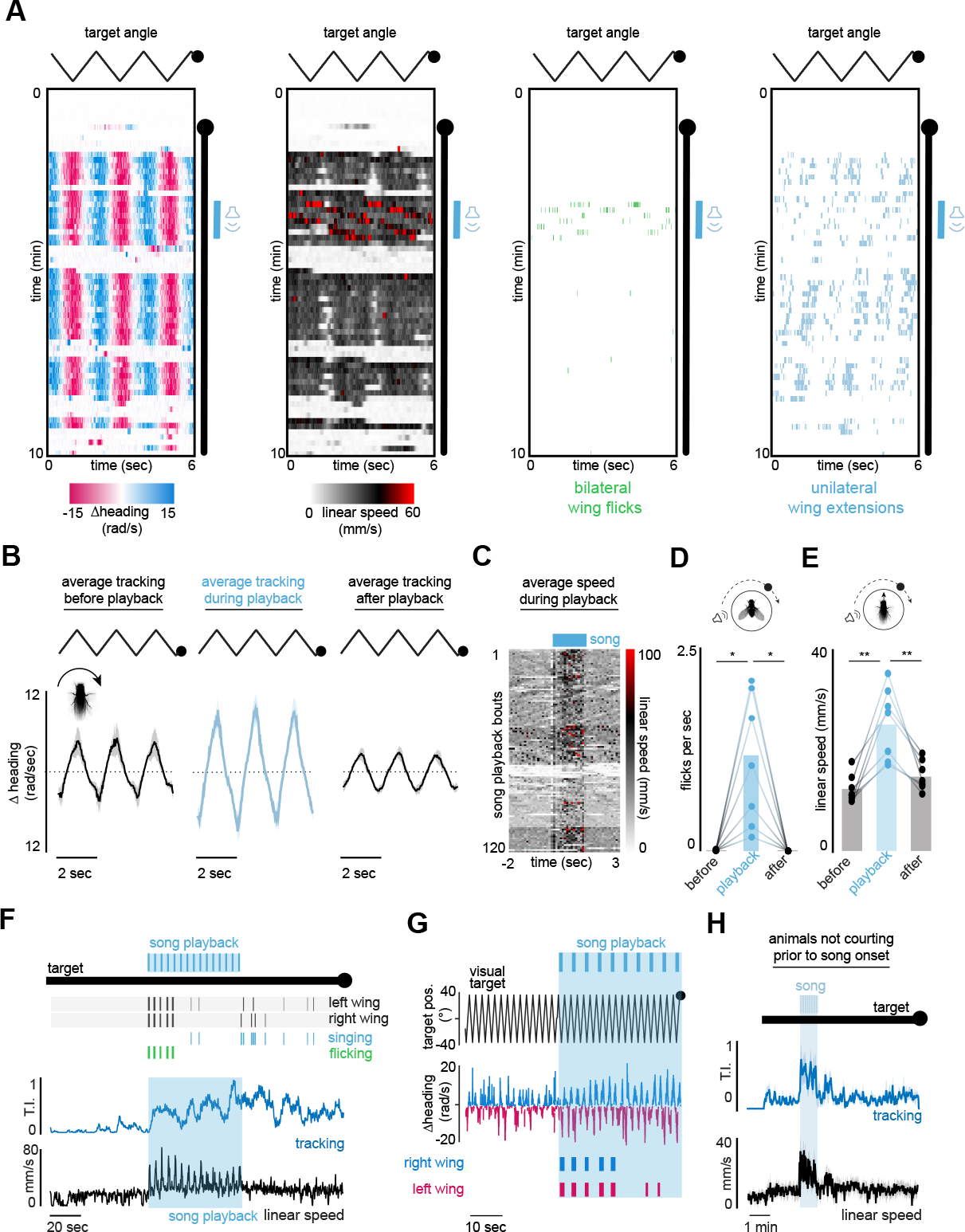
Pulse song playback can both drive agonistic displays and enhance courtship pursuit, related to Figure 6. (A) Representative example of a tethered male’s turning speed (left), linear speed (middle left), bilateral wing flicks (middle right), and unilateral wing extensions (right) over the course of one 10-minute virtual-reality trial. Black line indicates where the visual target was oscillating back and forth in front of the male; blue line indicates period of pulse-song playback. Each bin in the heat map reflects a single frame; all frames were aligned to the oscillating visual stimulus; time progresses from left to right over 6 sec, followed by from top to bottom. (**B**) Average male turning velocity in response to the oscillating visual target, averaged across 3 consecutive cycles, prior to, during, and after song playback. (**C**) Heat map of male linear speed across animals aligned to the onset of individual pulse trains. (**D**) Number of flicks performed per second by tethered males before, during, or after playback of pulse song. (**E**) Average linear speed of tethered males before, during, or after playback of pulse song. (**F**) Representative behavior of a tethered male prior to, during, and after the playback of conspecific courtship song. Black hatch marks denote left and right wing extensions; light blue hatch marks indicate bouts of unilateral wing extensions; green hatch marks indicate bouts of bilateral wing flicks. The male’s tracking index (see methods) is plotted in blue, and his linear speed in black. The male transitions from courting the fictive female by singing to charging towards it while flicking upon song-playback. Note that the male did not exhibit courtship prior to song playback, indicating that agonistic actions can be evoked even if males were not courting. (**G**) Zoomed in view of the example as (E) around the onset of the acoustic stimulus showing the induction of visual pursuit (left-right turning) following playback of pulse song. (**H**) Average male tracking index, aligned to the onset of song playback, across males that did not exhibit visual pursuit prior to the onset of song playback. Shaded lines show mean ± s.e.m. Details of sample sizes are given in **TableS2**.

**Figure S9.**
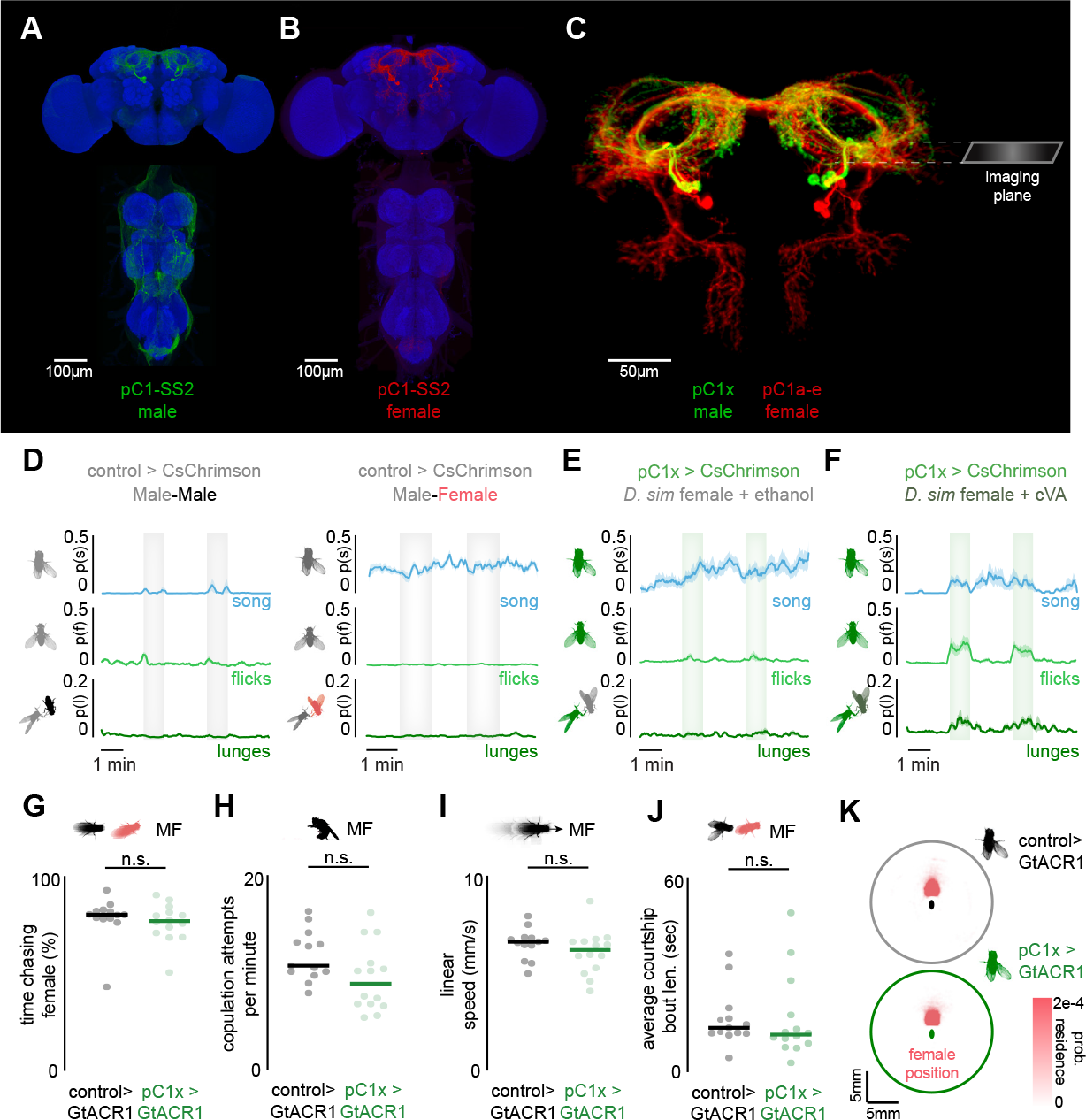
Sexually dimorphic pC1x neurons regulate agonistic but not courtship behaviors, related to Figure 6. ***(**A**-**C**) Examples of immunostained neurons labeled by pC1-SS2 in the brain and VNC of males (A) and females (B). (C) shows a zoomed in view of the neurons labeled in the central brain in males (green) and females (red). The approximate location of imaged neurites is denoted by the imaging plane (right). Note that females possess one more neuron (pC1a-e)^14,57,73,76^. Images adapted from the Janelia FlyLight split-GAL4 collection. (**D**) Comparison of component behaviors in control males. left: Average probability of control males performing unilateral wing extensions (top), bilateral wing flicks (middle), or lunge attacks (bottom) paired with a wingless mute male prior to, during, and after two 1 min bouts of red-light stimulation (shaded regions). right: Same as (left), but for control males paired with a female. (**E** and **F**) Same as (D), but for males expressing CsChrimsonin pC1x neurons when paired with a *D. simulans* female (E) or a *D. simulans* female perfumed with the male pheromone cVA. Note that *D. simulans* females possess the same cuticular pheromones as *D. melanogaster* males, but do not produce cVA. (**G**-**K**) Comparison of the courtship performance of control males and males in which pC1x neurons were optogenetically silenced via expression of GtACR1 in MF assays, showing the fraction of time males spent chasing female (G), the frequency of their copulation attempts (H), their average linear speed (I), the average length of their courtship bouts (J), and the positional occupancy of the female relative to the male (K). Shaded lines show mean ± s.e.m.; Lines with dots represent mean and individual animals. n.s., P > 0.05. Details of statistical analyses and sample sizes are given in **Table S2**.

**Figure S10.**
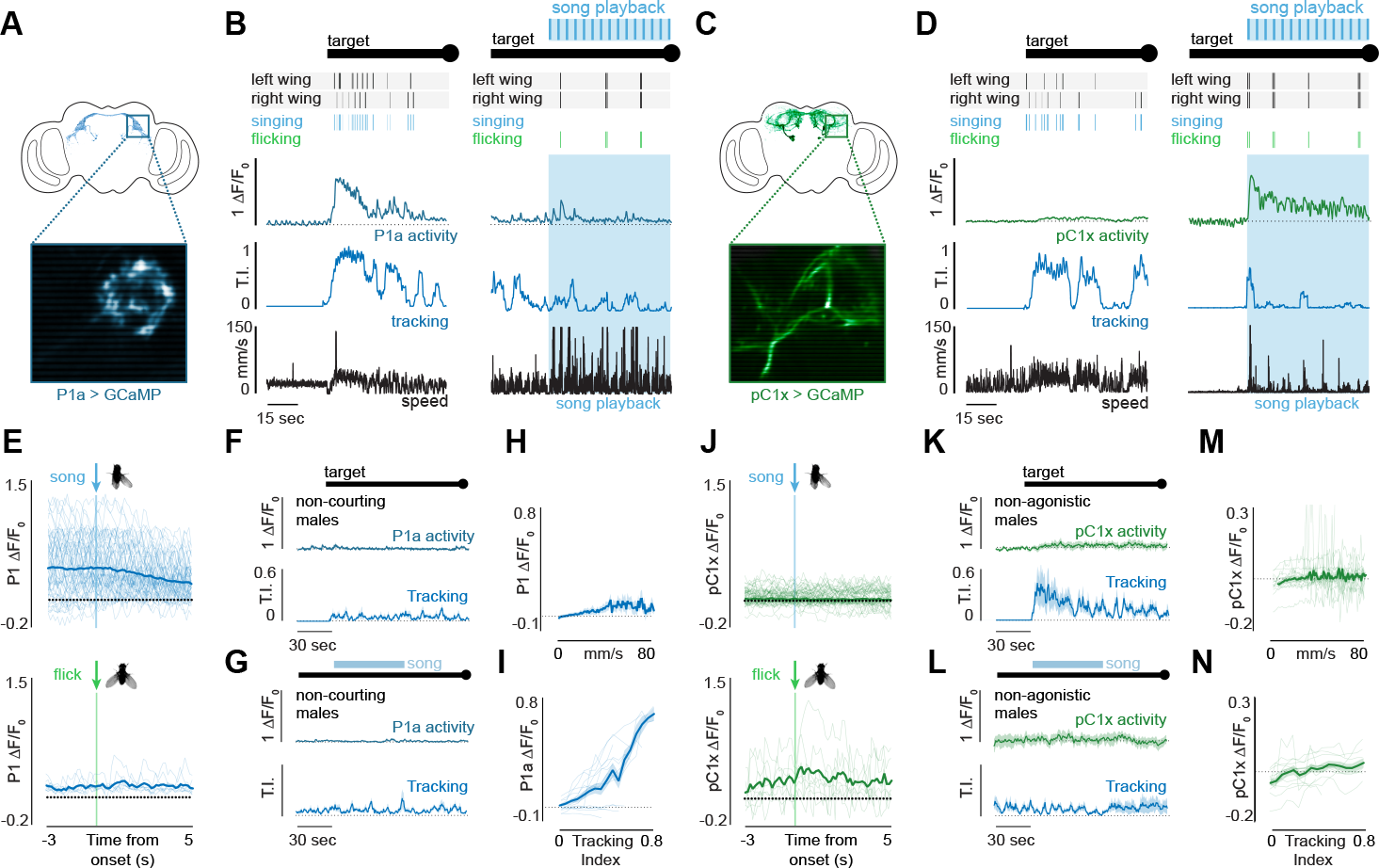
P1 and pC1x activity are correlated with different social behaviors, related to Figure 6. ***(**A**) Schematic showing the imaging plane used for recording from P1a neurons during tethered behavior. (**B**) Characteristic examples of recorded P1a activity (∆F/F_0_) during courtship (left) versus during agonistic behaviors (right). Black hatch marks denote left and right wing extensions; light blue hatch marks indicate bouts of unilateral wing-extensions (singing); green hatch marks indicate bouts of bilateral wing flicks. The male’s tracking index (T.I., see methods) is plotted in blue, and his linear velocity in black. (**C** and **D**) Same as A and B, but showing a characteristic example of the activity of pC1x neurons (∆F/F_0_, green) during courtship (right) or agonistic behavior (left). (**E**) Average activity (∆F/F_0_) of P1a neurons, aligned to the onset of unilateral wing extensions (top) or bilateral wing flicks in tethered males. Thin lines denote individual trials. (**F**) Average P1a activity (∆F/F_0_, top) and tracking index (bottom) of tethered males that did not exhibit any courtship (wing extensions, pursuit) towards the visual target, aligned to either the onset of visual motion (black line, left) or to the onset of pulse song playback (blue box, right). P1a neurons are not directly activated by song but correlate with enhanced pursuit that occurs frequently (but not invariably) with song playback. (**H** and **I**) Activity of P1a neurons (average ∆F/F_0_) plotted against the male’s linear speed (H) or tracking index (I). (**J, K, L, M**, and **N**) Same as E-I, but for the recorded activity of pC1x neurons. Shaded lines show mean ± s.e.m.; thin lines denote individual animals. Details of sample sizes are given in **TableS2**.

## Notes

### Competing Interest Statement

The authors have declared no competing interest.

