## Supplementary material for "Male-male interactions shape mate selection in *Drosophila*": TableS1

### Table S1
### Genotypes by figure

| Figure | Manipulated sex | Name | Full genotype |
|---|---|---|---|
| 1B-D, S1B-D | N/A | wild type | Wild-type Canton-S strain |
| 1E | N/A | wild type | Wild-type Canton-S, CO-4N, and ZH42 strains |
| 1F-I | N/A | wild type | Wild-type Canton-S strain |
| 2B-N, S2A-M, S3B-H | N/A | wild type | Wild-type Canton-S strain |
| 3A, S4A | male | R85A12>CsChrimson | 20XUAS-CsChrimson-mVenus @ attP18; +/+; R85A12-Gal4 @ attp2/+ |
| S4B | female | R85A12>CsChrimson | 20XUAS-CsChrimson-mVenus @ attP18; +/+; R85A12-Gal4 @ attp2/+ |
| 3B | male | control>CsChrimson | w+; +/20XUAS-CsChrimson-mCherry @ su(Hw)attP5; Gal4 @ attp2/+ |
| | | R85A12>CsChrimson | w+; +/20XUAS-CsChrimson-mCherry @ su(Hw)attP5; R85A12-Gal4 @ attp2/+ |
| S4C | male | control>CsChrimson | w+; +/20XUAS-CsChrimson-mCherry @ su(Hw)attP5; Gal4 @ attp2/+ |
| | | R85A12>CsChrimson | w+; +/20XUAS-CsChrimson-mCherry @ su(Hw)attP5; R85A12-Gal4 @ attp2/+ |
| | | R85A12∩Otd>CsChrimson | 20xUAS-FRT>STOP-FRT>-CsChrimson-mVenus @ attp18; +/otd-nls:FLPe @ attP40; R85A12-Gal4 @ attp2/TM6B |
| | female | R85A12>CsChrimson | w+; +/20XUAS-CsChrimson-mCherry @ su(Hw)attP5; R85A12-Gal4 @attp2/+ |
| 3C-D | male | control>Kir2.1 | w*; +/+; Gal4 @ attP2/5XUAS-Kir2.1.EGFP |
| | | R85A12>Kir2.1 | w*; +/+; R85A12-Gal4 @ attP2/5XUAS-Kir2.1.EGFP |
| 3E, S4D-E | male | R85A12>CsChrimson | w+; +/20XUAS-CsChrimson-mCherry @ su(Hw)attP5; R85A12-Gal4 @attp2/+ |
| S4F-G | male | control>GtACR1 | w+; p65.AD @ attP40/+; Gal4.DBD @ attp2/UAS-GtACR1.d.EYFP @ attP2 |
| | | pIP10>GtACR1 | w+; VT0405560-p65.AD/+; VT043047-Gal4.DBD @ attp2/UAS-GtACR1.d.EYFP @ attP2 |
| 3F-G, 3I, 3K, S4H-I, S5A, S5D-I | N/A | wild type | Wild-type Canton-S strain |
| 3H, 3J, 3L | male | control>Kir2.1 | w*; +/+; Gal4 @ attP2/5XUAS-Kir2.1.EGFP |
| | | R85A12>Kir2.1 | w*; +/+; R85A12-Gal4 @ attP2/5XUAS-Kir2.1.EGFP |
| S5B-C | male | Wild type (MF/MMF) | Wild-type Canton-S strain |
| | | control>Kir2.1 | w*; +/+; Gal4 @ attP2/5XUAS-Kir2.1.EGFP |
| | | R85A12>Kir2.1 | w*; +/+; R85A12-Gal4 @ attP2/5XUAS-Kir2.1.EGFP |
| 4B-D, S6A-B | female | vpoEN>GCaMP | w1118/w+; VT017181-p65.AD @ attP40/+; VT002215-Gal4.DBD @ |

| | | | |
|---|---|---|---|
| | | | attP2/20XUAS-IV-Syn21-Op-jGCaMP7f @ VK00005, 20XUAS-IV-Syn21-Op-jGCaMP7f @ attP2 |
| | | vpoIN>GCaMP | w1118/w+; R12D09-p65.AD @ attP40/+; VT008473-Gal4.DBD @ attP2/20XUAS-IV-Syn21-Op-jGCaMP7f @ VK00005, 20XUAS-IV-Syn21-Op-jGCaMP7f @ attP2 |
| 4E | female | vpoEN>GCaMP | w1118/w+; VT017181-p65.AD @ attP40/+; VT002215-Gal4.DBD @ attP2/20XUAS-IV-Syn21-Op-jGCaMP7f @ VK00005, 20XUAS-IV-Syn21-Op-jGCaMP7f @ attP2 |
| 4F-H, S6C, S6E-G | N/A | wild type | Wild-type Canton-S strain |
| S6D | female | control>Kir2.1 | w*; p65.AD @ attp40/+; Gal4.DBD @ attP2/5XUAS-Kir2.1.EGFP |
| | | vpoDN>Kir2.1 | w*; R31D07-p65.AD @ attP40/+; R52F12-Gal4.DBD @ attp2/5XUAS-Kir2.1.EGFP |
| | | DNp13>Kir2.1 | w*; VT038159-p65.AD @ attP40/+; R20C08-Gal4.DBD @ attP2/5XUAS-Kir2.1.EGFP |
| 5A-I | N/A | wild type | Canton-S strain |
| 5J-K, S7A-H | male | control>Kir2.1 | w*; +/+; Gal4 @ attP2/5XUAS-Kir2.1.EGFP |
| | | R85A12>Kir2.1 | w*; +/+; R85A12-Gal4 @ attP2/5XUAS-Kir2.1.EGFP |
| S7I-L | N/A | wild type | Wild-type Canton-S strain |
| 6B, 6C, S8A | N/A | pC1-SS2>GCaMP | w1118; +/+; VT002064-p65.AD @ attP2, dsx-DBD/20XUAS-IVS-jGCaMP7s @ VK00005 |
| 6D, S8B-H | N/A | P1a>GCaMP | w1118; 15A01-p65.AD @ attp40/+; 71G01-Gal4.DBD @ attP2/20XUAS-IVS-jGCaMP7s @ VK00005 |
| | | pC1-SS2>GCaMP | w1118; +/+; VT002064-p65.AD @ attP2, dsx-DBD/20XUAS-IVS-jGCaMP7s @ VK00005 |
| | | wild type | Wild-type Canton-S strain |
| 6E, S9A-C | male, female | pC1-SS2>CsChrimson | w1118, 20xUAS-CsChrimson-mVenus, UAS-syt-HA @ attP18; +/+; VT002064-p65.AD @ attP2, dsx-DBD/+ |
| 6F | male | P1a>CsChrimson | w1118, 20xUAS-CsChrimson-mVenus @ attP18; 15A01-p65.AD @ attp40/+; 71G01-Gal4.DBD @ attP2/+ |
| | | pC1-SS2>CsChrimson | w1118, 20xUAS-CsChrimson-mVenus @ attP18; +/+; VT002064-p65.AD @ attP2, dsx-DBD/+ |
| 6G, S9D | male | control>CsChrimson | w1118, 20xUAS-CsChrimson-mVenus @ attP18; p65.AD @ attp40/+; Gal4.DBD @ attP2/+ |
| | | P1a>CsChrimson | w1118, 20xUAS-CsChrimson-mVenus @ attP18; 15A01-p65.AD @ attp40/+; 71G01-Gal4.DBD @ attP2/+ |
| 6H | male | pC1-SS2>CsChrimson | w1118, 20xUAS-CsChrimson-mVenus @ attP18; +/+; VT002064-p65.AD @ attP2, dsx-DBD/+ |

| | | | |
|---|---|---|---|
| S9E-F | male | pC1-SS2>CsChrimson | w1118, 20xUAS-CsChrimson-mVenus @ attP18; +/+; VT002064-p65.AD @ attP2, dsx-DBD/+ |
| S9G-K | male | control>GtACR1 | w+; p65.AD @ attP40/+; Gal4.DBD @ attp2/UAS-GtACR1.d.EYFP @ attP2 |
| | | pC1-SS2>GtACR1 | w+; +/+; VT002064-p65.AD @ attP2, dsx-DBD/UAS-GtACR1.d.EYFP @ attP2 |
| 6I-J, S10A-B, S10E-I | male | P1a>GCaMP | w1118; 15A01-p65.AD @ attp40/+; 71G01-Gal4.DBD @ attP2/20XUAS-IVS-jGCaMP7s @ VK00005 |
| 6K-L, S10C-D, S10J-N | male | pC1-SS2>GCaMP | w1118; +/+; VT002064-p65.AD @ attP2, dsx-DBD/20XUAS-IVS-jGCaMP7s @ VK00005 |
| 6M-N | male | control>GtACR1 | w+; p65.AD @ attP40/+; Gal4.DBD @ attp2/UAS-GtACR1.d.EYFP @ attP2 |
| | | pC1-SS2>GtACR1 | w+; +/+; VT002064-p65.AD @ attP2, dsx-DBD/UAS-GtACR1.d.EYFP @ attP2 |
| 6O-Q | male | pC1-SS2>CsChrimson | w1118, 20xUAS-CsChrimson-mVenus @ attP18; +/+; VT002064-p65.AD @ attP2, dsx-DBD/+ |
