## Supplementary material for "Male-male interactions shape mate selection in *Drosophila*": TableS2

### Table S2
### Statistics and Sample Sizes

| Figure | Group | Sample size | Statistical tests | p value | Additional notes |
|---|---|---|---|---|---|
| **1B** | MMF | n = 18 triads, n = 36 males | Friedman test followed by Dunn's multiple comparison test | pursuing female: p>0.9999 | Winner vs Loser |
| | | | | singing: p>0.9999 | Winner vs Loser |
| | | | | attempt copulating: p>0.9999 | Winner vs Loser |
| | | | | distance to female: p>0.9999 | Winner vs Loser |
| | | | | Speed: p>0.9999 | Winner vs Loser |
| | | | | body size: p>0.9999 | Winner vs Loser |
| **1C, 1D** | MMF | n = 18 triads, n = 36 males | N/A | N/A | |
| **1E** | MF | n = 96 pairs | N/A | N/A | |
| **1F** | normal male (control) | n = 26 pairs | N/A | N/A | |
| | wingless male | n = 25 pairs | Log-rank (Mantel-Cox) test | chi square = 42.03, p < 0.0001 | |
| | wingless male + sound playback | n = 27 pairs | Log-rank (Mantel-Cox) test | chi square = 0.8866, p = 0.3464 | |
| **1G** | top | n = 68 triads | chi-square test | chi square = 5.882, p = 0.0153 | |
| | bottom | n = 30 triads | chi-square test | chi square = 0.000, p > 0.9999 | |
| **1H** | winged versus wingless competition triad, wingless copulated | n = 24 triads | N/A | N/A | |
| **1I** | winged versus wingless competition triad, winged copulated | n = 44 triads | One-way ANOVA followed by Šidák's multiple comparisons test with pooled variance | p=0.3639 | Winged vs. wingless distance: wingless copulated |
| | | | | p<0.0001 | Winged distance (wingless copulated) vs. wingless distance (winged copulated) |
| | | | | p=0.4082 | Winged distance (wingless copulated) vs. winged distance (winged copulated) |
| | winged versus wingless competition triad, wingless copulated | n = 24 triads | | p<0.0001 | Wingless distance (wingless copulated) vs. wingless distance (winged copulated) |
| | | | | p=0.9998 | Wingless distance (wingless copulated) vs. winged distance (winged copulated) |
| | | | | p<0.0001 | Winged vs. wingless distance: winged copulated |
| **2B** | MF | n = 20 pairs | Brown-Forsythe and Welch ANOVA tests, followed by Dunnett's T3 multiple | pursuing female: p<0.0001 | MF vs MMF |
| | MMF | n = 18 triads, n = 36 males | | | |
| | MF | n = 20 pairs | | singing: p<0.0001 | MF vs MMF |
| | MMF | n = 18 triads, n = 36 males | | | |
| | MF | n = 20 pairs | | | MF vs MMF |

| | | | | | |
|---|---|---|---|---|---|
| | MMF | n = 18 triads, n = 36 males | comparisons test | disengaged: p<0.0001 | |
| | MF | n = 20 pairs | | male speed: p<0.0001 | MF vs MMF |
| | MMF | n = 18 triads, n = 36 males | | | |
| | MF | n = 20 pairs | | flicking: p<0.0001 | MF vs MMF |
| | MMF | n = 18 triads, n = 36 males | | | |
| **2E** | MMF | n = 18 triads, n = 36 males | N/A | N/A | |
| **2F** | MMF | n = 18 triads, n = 36 males | N/A | N/A | |
| **2G** | MMF | n = 18 triads, n = 36 males | N/A | N/A | |
| **2H** | MMF | n = 18 triads, n = 36 males | N/A | N/A | |
| **2I** | MMF | n = 18 triads, n = 36 males | N/A | N/A | |
| **2J** | MMF | n = 18 triads, n = 36 males | N/A | N/A | |
| **2K** | MMF | n = 18 triads, n = 36 males | N/A | N/A | |
| **2L** | no sound | n = 10 trials, n = 20 males | Brown-Forsythe and Welch ANOVA tests, followed by Dunnett's T3 multiple comparisons test | N/A | |
| | noise playback | n = 9 trials, n = 18 males | | p = 0.4872 | noise vs. no sound |
| | *D. simulans* song playback | n = 9 trials, n = 18 males | | p = 0.2733 | *D.sim* vs. no sound |
| | *D. melanogaster* song playback | n = 10 trials, n = 20 males | | p<0.0001 | *D.mel* vs. no sound |
| **2M** | normal (control) | n = 18 triads, 36 males | One-way ANOVA followed by Dunnett's multiple comparisons test | N/A | |
| | deaf | n = 17 triads, 34 males | | p = 0.1660 | Deaf vs. control |
| | deaf and anosmic | n = 16 triads, 32 males | | p < 0.0001 | Deaf/mute vs. control |
| **2N** | normal (control) | n = 18 triads, 36 males | One-way ANOVA followed by Dunnett's multiple comparisons test | N/A | |
| | deaf | n = 17 triads, 34 males | | p = 0.2939 | Deaf vs. control |
| | deaf and anosmic | n = 16 triads, 32 males | | p < 0.0001 | Deaf/mute vs. control |
| **3B** | control>CsChrimson | n = 10 males | Mann-Whitney test | p < 0.0001 | |
| | R85A12>CsChrimson | n = 10 males | | | |
| **3C** | control>Kir2.1 MMF | n = 9 triads, 18 males | Welch's two-tailed t-test | p < 0.0001 | |
| | R85A12>Kir2.1 MMF | n = 9 triads, 18 males | | | |
| **3D** | control>Kir2.1 MF | n = 12 pairs | Student's two-tailed t-test | p = 0.4722 | |
| | R85A12>Kir2.1 MF | n = 12 pairs | | | |
| **3I** | MF | n = 33 pairs | N/A | N/A | |
| | MMF | n = 33 triads | | | |
| **3J** | control>Kir2.1 MMF | n = 10 triads | N/A | N/A | |
| | R85A12>Kir2.1 MMF | n = 14 triads | | | |
| **3K** | MF | n = 33 pairs | Welch's two-tailed t-test | p < 0.0001 | |
| | MMF | n = 33 triads | | | |
| **3L** | control>Kir2.1 MMF | n = 10 triads | Welch's two-tailed t-test | p < 0.0001 | |
| | R85A12>Kir2.1 MMF | n = 14 triads | | | |
| **4B** | vpoEN>GCaMP | n = 7 females | N/A | N/A | |
| | vpoIN>GCaMP | n = 9 females | | | |
| **4C** | vpoEN>GCaMP | n = 11 females | N/A | N/A | |
| | vpoIN>GCaMP | n = 7 females | | | |
| **4D** | vpoEN>GCaMP | n = 25 females | N/A | N/A | |
| | vpoIN>GCaMP | n = 23 females | N/A | N/A | |
| **4E** | vpoEN>GCaMP | n = 7 females | N/A | N/A | |

| | | | | |
|---|---|---|---|---|
| **4G** | MF | n = 20 pairs | N/A | N/A |
| | MMF | n = 18 triads, n = 36 males | | |
| **4H** | MF sound playback | n = 44 pairs | Log-rank (Mantel-Cox) test | chi square = 22.38, p < 0.0001 |
| | MMF sound playback | n = 35 pairs | | |
| **5A** | MMF | n = 33 triads | N/A | N/A |
| **5B** | MF | n = 33 pairs | N/A | N/A |
| | MMF | n = 33 triads | | |
| **5C** | MMF | n = 26 triads | N/A | N/A |
| **5D** | MMF | n = 18 triads, n = 36 males | N/A | N/A |
| **5F (left)** | MMF | n = 2295 frames from 301 bouts of flicking across one assay | N/A | N/A |
| **5F (right)** | MMF | n = 18 triads, n = 36 males | N/A | N/A |
| **5G** | MMF | n = 18 triads, 36 males | Repeated-Measures one-way ANOVA followed by Šidák's multiple comparisons test | Rival behind, before vs. after flick: p < 0.00001 |
| | | | | Rival in front, before vs. after flick: p = 0.6524 |
| **5H** | MMF | n = 18 triads, n = 36 males | N/A | N/A |
| **5I** | MMF | n = 18 triads, n = 36 males | N/A | N/A |
| **5J** | competition triads | n = 11 triads | N/A | N/A |
| **5K** | competition triads | n = 64 triads | chi-square test | chi square = 27.56, p < 0.0001 |
| **6D** | Tethered males | n = 7 males | N/A | N/A |
| **6F** | P1a > CsChrimson Solitary males | n = 24 males | N/A | N/A |
| | pC1x > CsChrimson Solitary males | n = 14 males | N/A | N/A |
| **6G, S9D (left)** | control>CsChrimson with wingless male | n = 10 pairs | N/A | N/A |
| | P1a>CsChrimson with wingless male | n = 10 pairs | N/A | N/A |
| **6H, S9D (left)** | control>CsChrimson with wingless male | n = 10 pairs | N/A | N/A |
| | pC1x>CsChrimson with wingless male | n = 10 pairs | N/A | N/A |
| **6I, 6J** | P1a>GCaMP | n = 8 males | N/A | N/A |
| **6K, 6L** | pC1x>GCaMP | n = 5 males | N/A | N/A |
| **6M** | control>GtACR1 MMF | n = 15 triads, 30 males | Welch's two-tailed t-test | p < 0.0001 |
| | pC1x>GtACR1 MMF | n = 14 triads, 28 males | | |
| **6N** | control>GtACR1 MF | n = 13 pairs | Student's two-tailed t test | p = 0.0626 |
| | pC1x>GtACR1 MF | n = 14 pairs | | |
| **6O, S9D (right)** | control>CsChrimson MF | n = 9 pairs | N/A | N/A |
| | pC1x>CsChrimson MF | n = 8 pairs | | |
| **6P** | pC1x>CsChrimson with *D. melanogaster* female (control) | n = 8 pairs | Kruskal-Wallis test followed by Dunn's multiple comparisons test | N/A |
| | pC1x>CsChrimson with *D. simulans* female | n = 10 pairs | | *D. sim.* (mock) vs. *D. mel.:* p = 0.1192 |
| | pC1x>CsChrimson with *D. simulans* female + cVA | n = 7 pairs | | *D. sim.* (cVA) *vs. D. mel.:* p<0.0001 |

| | | | | | |
|---|---|---|---|---|---|
| **6Q** | pC1x>CsChrimson with *D. melanogaster* female (control) | n = 8 pairs | Kruskal-Wallis test followed by Dunn's multiple comparisons test | N/A | |
| | pC1x>CsChrimson with *D. simulans* female | n = 10 pairs | | *D. sim.* (mock) vs. *D. mel.:* p = 0.6316 | |
| | pC1x>CsChrimson with *D. simulans* female + cVA | n = 7 pairs | | *D. sim.* (cVA) *vs. D. mel.:* p=0.0002 | |
| **S1B** | MF | 2 pairs (20,000 frames per behavior) | N/A | N/A | |
| | MMF | 2 triads, 4 males (20,000 frames per behavior) | N/A | N/A | |
| **S1D** | winged versus wingless competition triad | n = 68 triads | N/A | N/A | |
| | one wingless male and one female | n = 38 pairs | N/A | N/A | |
| | two wingless males and one female | n = 41 triads | N/A | N/A | |
| **S2A** | MF | n = 20 pairs | Welch's two-tailed t-test | speed: p < 0.0001 | |
| | MMF | n = 18 triads | | female being pursued: p < 0.0001 | |
| **S2B** | MF | n = 20 pairs | Student's two-tailed t-test | p = 0.0004 | |
| | MMF | n = 18 triads, 36 males | | | |
| **S2C** | MMF | n = 18 triads, 36 males | One-way ANOVA followed by Šidák's multiple comparisons test | Both pursuing female vs chance overlap: p < 0.0001 | Chance overlap quantified as squared probability of either male in behavior |
| | | | | Both singing vs chance overlap: p = 0.9984 | |
| | | | | Both disengaged vs chance overlap: p = 0.8593 | |
| | | | | Both attempting copulation vs chance overlap: p = 0.8908 | |
| | | | | Both in optimal position vs chance overlap: p <0.0001 | |
| | | | | Both flicking vs. chance overlap: p>0.9999 | |
| **S2D** | MF | n = 20 pairs | N/A | N/A | |
| | MMF | n = 18 triads, 36 males | N/A | N/A | |
| **S2E** | MMF | n = 18 triads, 36 males | N/A | N/A | |
| **S2F** | MMF | n = 18 triads, 36 males | N/A | N/A | |

| | | | | | |
|---|---|---|---|---|---|
| **S2G** | MMF | n = 18 triads, 36 males | N/A | N/A | |
| **S2H-K** | MMF | n = 240 song bouts from 18 triads, 36 males | Repeated-Measures one-way ANOVA with Geisser-Greenhouse correction, followed by Šidák's multiple comparisons test with individual variances computed for each comparison | Facing angle, before vs after rival song: p < 0.0001 | |
| | | | | Distance to female, before vs after rival song: p < 0.0001 | |
| **S2L-M** | MMF | n = 2142 song bouts from 18 triads, 36 males | Student's paired two-tailed t-test, before vs. after rival flicks | p<0.0001 | |
| **S3B** | *D. simulans* song playback | n = 9 trials, n = 18 males | N/A | N/A | |
| **S3C** | white noise playback | n = 9 trials, n = 18 males | N/A | N/A | |
| **S3D** | no sound | n = 10 trials, n = 20 males | Kruskal-Wallis test followed by Dunn's multiple comparisons test | N/A | |
| | noise playback | n = 9 trials, n = 18 males | | p = 0.3470 | noise vs. no sound |
| | *D. simulans* song playback | n = 9 trials, n = 18 males | | p = 0.4171 | *D.sim* vs. no sound |
| | *D. melanogaster* song playback | n = 10 trials, n = 20 males | | p = 0.6259 | *D.mel* vs. no sound |
| **S3E, S3F** | *D. melanogaster* song playback to MF | n = 7 pairs | N/A | N/A | |
| **S3G** | MMF | n = 18 triads, 36 males | One-way ANOVA followed by Tukey's multiple comparisons test | MMF vs MF: p<0.0001 | |
| | MF | n = 20 pairs | | MMF vs Playback: p = 0.0995 | |
| | *D. melanogaster* song playback to MF | n = 7 pairs | | MMF vs MF: p<0.0001 | |
| **S3H** | MMF | n = 18 triads, 36 males | One-way ANOVA followed by Tukey's multiple comparisons test | MMF vs MF: p<0.0001 | |
| | MF | n = 20 pairs | | MMF vs Playback: p = 0.0530 | |
| | *D. melanogaster* song playback to MF | n = 7 pairs | | MMF vs MF: p<0.0001 | |
| **S4C** | control>CsChrimson male (control) | n = 11 males | Kruskal-Wallis test followed by Dunn's multiple comparisons test | N/A | |
| | R85A12>CsChrimson male | n = 10 males | | p < 0.0001 | R85A12>CsChrimson male vs. control |
| | R85A12∩Otd>CsChrimson male | n = 10 males | | p > 0.9999 | R85A12∩Otd>CsChrimson male vs. control |
| | R85A12∩Otd>CsChrimson male, high light | n = 10 males | | p > 0.9999 | R85A12∩Otd>CsChrimson male, high light vs. control |
| | R85A12>CsChrimson female | n = 8 females | | p > 0.9999 | R85A12>CsChrimson female vs. control |
| | R85A12>CsChrimson female, high light | n = 8 females | | p > 0.9999 | R85A12>CsChrimson female, high light vs. control |
| **S4D, S4E** | R85A12>CsChrimson | n = 15 males | N/A | N/A | |
| **S4F** | control>GtACR1 MF | n = 14 pairs | Student's two-tailed t-test | p < 0.0001 | |
| | pIP10>GtACR1 MF | n = 14 pairs | | | |

| | | | | | |
|---|---|---|---|---|---|
| **S4G** | control>GtACR1 MM | n = 14 trials, 28 males | Repeated measures one-way ANOVA followed by Tukey's multiple comparisons test | sound off, light off vs sound on, light off: p < 0.0001 | |
| | | | | sound off, light off vs sound on, light on: p < 0.0001 | |
| | | | | sound on, light off vs sound on, light on: p = 0.9076 | |
| | pIP10>GtACR1 MM | n = 14 trials, 28 males | | sound off, light off vs sound on, light off: p < 0.0001 | |
| | | | | sound off, light off vs sound on, light on: p < 0.0001 | |
| | | | | sound on, light off vs sound on, light on: p = 0.5156 | |
| **S4H** | MF | n = 33 pairs | Student's two-tailed t-test | p < 0.0001 | |
| | MMF | n = 33 triads | | | |
| **S4I** | MF | n = 33 pairs | Welch's two-tailed t-test | p < 0.0001 | |
| | MMF | n = 33 triads | | | |
| **S5A** | MMF | n = 4 triads | N/A | N/A | |
| **S5B-C** | MF | n = 33 pairs | N/A | N/A | |
| | MMF | n = 33 triads | N/A | N/A | |
| | control>Kir2.1 MMF | n = 10 triads | N/A | N/A | |
| | R85A12>Kir2.1 MMF | n = 14 triads | N/A | N/A | |
| **S5G** | pulse song classification accuracy | n = 5 pairs | N/A | N/A | |
| | sine song classification accuracy | n = 5 pairs | N/A | N/A | |
| **S5I** | MF | n = 33 pairs | N/A | N/A | |
| | MMF | n = 33 triads | N/A | N/A | |
| **S6A** | vpoEN>GCaMP | n = 7 females | N/A | N/A | |
| | vpoIN>GCaMP | n = 8 females | N/A | N/A | |
| **S6B** | vpoEN>GCaMP | n = 7 females | N/A | N/A | |
| | vpoIN>GCaMP | n = 8 females | N/A | N/A | |
| **S6D** | control>Kir2.1 MF (control) | n = 14 females | Kruskal-Wallis test followed by Dunn's multiple comparisons test | N/A | |
| | DNp13>Kir2.1 MF | n = 14 females | | p = 0.8233 | DNp13 > Kir2.1 vs. control |
| | vpoDN>Kir2.1 MF | n = 9 females | | p = 0.0230 | vpoDN > Kir2.1 vs. control |
| **S6E, S6G (top)** | MF | n = 1826 song bouts from 20 pairs | N/A | N/A | |
| **S6E, S6G (bottom)** | MMF | n = 2142 song bouts from 18 triads, 36 males | N/A | N/A | |
| **S7A** | control>Kir2.1 MF | n = 12 pairs | Student's two-tailed t-test | p = 0.0659 | |
| | R85A12>Kir2.1 MF | n = 9 pairs | | | |
| **S7B** | control>Kir2.1 MF | n = 12 pairs | N/A | N/A | |
| | R85A12>Kir2.1 MF | n = 9 pairs | | | |
| **S7C** | control>Kir2.1 MF | n = 12 pairs | Student's two-tailed t-test | p = 0.1908 | |
| | R85A12>Kir2.1 MF | n = 9 pairs | | | |

| | | | | |
|---|---|---|---|---|
| **S7D** | control>Kir2.1 MF | n = 12 pairs | N/A | N/A |
| | R85A12>Kir2.1 MF | n = 9 pairs | | |
| **S7E** | control>Kir2.1 MF | n = 12 pairs | Student's two-tailed t-test | p = 0.2461 |
| | R85A12>Kir2.1 MF | n = 12 pairs | | |
| **S7F** | control>Kir2.1 MF | n = 12 pairs | N/A | N/A |
| | R85A12>Kir2.1 MF | n = 12 pairs | | |
| **S7G** | control>Kir2.1 MF | n = 15 pairs | Log-rank (Mantel-Cox) test | chi square = 0.05062, p = 0.8220 |
| | R85A12>Kir2.1 MF | n = 21 pairs | | |
| **S7H** | competition triads | n = 11 triads | Mann-Whitney test | p = 0.0041 |
| **S7I** | MMF | n = 3645 flicking bouts from 18 triads, 36 males | N/A | N/A |
| **S7J-K** | deaf male MMF | n = 17 triads, 34 males | N/A | N/A |
| **S7L** | normal MMF | n = 18 triads, 36 males | N/A | N/A |
| | deaf male MMF | n = 17 triads, 34 males | N/A | N/A |
| **S8B-C** | Tethered males | n = 7 males | N/A | N/A |
| **S8D** | Tethered males | n = 7 males | Friedman test followed by Dunn's multiple comparison test | Before vs. During: p = 0.0283 |
| | | | | After vs. During: p = 0.0122 |
| **S8E** | Tethered males | n = 7 males | Repeated-Measures one-way ANOVA followed by Dunn's multiple comparison test | Before vs. During: p = 0.0045 |
| **S8H** | Tethered males not courting prior to song | n = 7 males | N/A | N/A |
| **S9D** | control>CsChrimson, wingless male target | n = 10 pairs | N/A | After vs. During: p = 0.0078 |
| | control>CsChrimson, female target | n = 9 pairs | N/A | N/A |
| **S9E** | pC1x>CsChrimson, mock perfumed *D. sim* female target | n = 10 pairs | N/A | N/A |
| **S9F** | pC1x>CsChrimson, cVA perfumed *D. sim* female target | n = 7 pairs | N/A | N/A |
| **S9G-J** | control>GtACR1 | n = 13 pairs | Brown-Forsythe and Welch ANOVA tests, followed by Dunnett's T3 multiple comparisons test | Time chasing: p = 0.9644 |
| | | | | Copulation attempts: p = 0.1411 |
| | pC1x>GtACR1 | n = 14 pairs | | Male speed: p = 0.6825 |
| | | | | Courtship bout length: p > 0.9999 |
| **S9K** | control>GtACR1 | n = 13 pairs | N/A | N/A |
| | pC1x>GtACR1 | n = 14 pairs | N/A | N/A |
| **S10E (top)** | P1a>GCaMP | n = 57 unilateral wing extensions across 8 males | N/A | N/A |
| **S10E (bottom)** | P1a>GCaMP | n = 7 bilateral wing flicks across 8 males | N/A | N/A |
| **S10F-G** | P1a>GCaMP | n = 6 males | N/A | N/A |
| **S10H-I** | P1a>GCaMP | n = 14 males | Pearson's correlation | Tracking vs activity: r = |

| | | | | |
|---|---|---|---|---|
| | | | | 0.7628, p < 0.00001 |
| | | | | Speed vs activity: r = 0.1308, p < 0.00001 |
| **S10J (top)** | pC1x>GCaMP | n = 53 unilateral wing extensions across 5 males | N/A | N/A |
| **S10J (bottom)** | pC1x>GCaMP | n = 8 bilateral wing flicks across 5 males | N/A | N/A |
| **S10K-L** | pC1x>GCaMP | n = 8 males | N/A | N/A |
| **S10M-N** | pC1x>GCaMP | n = 13 males | Pearson's correlation | Tracking vs activity: r = 0.3467, p < 0.00001 |
| | | | | Speed vs activity: r = 0.1060, p < 0.00001 |
